# Non-canonical DNA and sequencing challenges in bird genomes

**DOI:** 10.1101/2025.10.17.683159

**Authors:** Linnéa Smeds, Jacob Sieg, Simona Secomandi, Chul Lee, Marco Sollitto, Jack A. Medico, Huiqing Zeng, Francesca Chiaromonte, Erich D. Jarvis, Giulio Formenti, Kateryna D. Makova

## Abstract

Non-canonical (non-B) DNA motifs can fold into structures such as G-quadruplexes (G4s), triplexes, and Z-DNA that regulate gene expression and promote mutagenesis in mammals. Yet they remain undercharacterized in other vertebrates, partly because they are difficult to sequence and often missing from draft assemblies. Using the telomere-to-telomere (T2T) zebra finch genome, the near-complete chicken genome, and seven additional high-quality assemblies spanning >100 million years of avian evolution, we present a genome-wide analysis of non-B DNA in birds. Non-B motifs cover 7.6% of the zebra finch and chicken genomes and are disproportionately enriched on their small, gene-rich dot chromosomes (e.g., 21.6% and 33.7% coverage, respectively). As in mammals, G4s are overrepresented at promoters and 5′UTRs (up to 6-fold). Introns on bird dot chromosomes show >9-fold enrichment of G4s and direct repeats. We provide experimental evidence of non-B DNA structure formation. Circular Dichroism spectra indicate that four common zebra finch G4 motifs fold *in vitro*. Genome-wide permanganate/S1 footprinting (PDAL-Seq) in a zebra finch cell line detects single-stranded DNA associated with non-B DNA motifs at transcribed and regulatory regions and at tandem repeats, telomeric and centromeric satellites, and Z-DNA-rich Ngaro retrotransposons. Finally, we find a negative relationship between non-B DNA coverage and PacBio HiFi sequencing depth on zebra finch dot chromosomes, potentially explaining in part why these chromosomes have resisted complete assembly. Our results suggest that non-B DNA contributes to gene regulation and repeat dynamics in birds and inform strategies for completing bird genomes.

## Introduction

Recent improvements in genome assembly, driven by advances in long-read sequencing and computational algorithms, have advanced the goal of generating *complete and gapless* reference genomes, i.e., telomere-to-telomere (T2T) assemblies (Rhie et al. 2021). The first published T2T genome was that of a human female (Nurk et al. 2022), followed by the complete human Y chromosome (Rhie et al. 2023), as well as the sex chromosomes (Makova et al. 2024) and the autosomes (Yoo et al. 2025) of several ape species. T2T genomes have proven to be essential for studies of segmental duplications (Vollger et al. 2022; Jeong et al. 2025), transposable elements (Hoyt et al. 2022), satellites (Altemose et al. 2022), and non-B DNA motifs (Smeds et al. 2025; Mohanty et al. 2025). Studies have shown that non-B DNA motifs—sequences that can form non-canonical, or non-B, DNA—are greatly enriched in newly added sequences in ape T2T genomes compared with previous assemblies (Smeds et al. 2025). Non-B DNA structures have recently emerged as novel functional elements (Guiblet et al. 2021b) and drivers of genome evolution (Makova and Weissensteiner 2023). Determining the complete repertoire of predicted non-B DNA motifs is crucial for understanding the roles of these structures in genomes, ranging from mutation hotspots that facilitate genome instability (Guiblet et al. 2021a; Makova and Weissensteiner 2023) to regulators of gene expression (Wang and Vasquez 2023; Esnault et al. 2025). Despite substantial progress in analyzing non-B DNA in human and great ape T2T genomes (Smeds et al. 2025; Mohanty et al. 2025), the occurrence of non-B DNA motifs in complete genomes of other species remains largely unexplored, in part due to the lack of T2T assemblies. Moreover, most of our knowledge of non-B DNA function comes from analyses of genomes of mammals and of model organisms, such as yeast (Wang and Vasquez 2023). Non-B DNA functions in the genomes of other species are still critically understudied.

Bird genomes have been included in several large-scale evolutionary studies of non-B DNA motifs, investigating either certain motif types or particular genomic regions. For example, a recent study investigated non-B DNA in UnTranslated Regions (UTRs) across hundreds of eukaryote species, including 23 bird species, and reported that G-quadruplexes (G4s) were the predominant non-B DNA motif type in bird UTRs, just as in mammals and reptiles (Gummadi et al. 2024). Another study analyzing over 150 bird species found that Z-DNA motifs were most common in bird promoter regions, and established a negative relationship between Z-DNA motif density and developmental time (Wang et al. 2024). Another study annotated non-B DNA motifs in promoters from over a thousand species across the tree of life, including six bird species, and compared them across large taxonomic groups (Yella and Vanaja 2023). However, to our knowledge, a detailed study of the non-B DNA motif landscape across bird genomes and among their chromosomes has been lacking.

More than 2,000 bird genomes are now publicly available (NCBI 2026). Yet almost none of them have been assembled to completeness. Bird genomes are smaller than mammalian genomes (∼1 Gbp vs. ∼3 Gbp for the haploid genome), with a much lower repeat content (∼10% vs. ∼25-50%) (International Chicken Genome Sequencing Consortium 2004; Warren et al. 2010; Liao et al. 2023). Nevertheless, they have proven challenging to sequence in full, likely due to the unique organization and content of their genomes, which consist of large *macrochromosomes* as well as multiple small GC-rich *microchromosomes* that contain many genes and have high recombination rates (Ellegren 2013). Most bird genome assemblies released in the 2010s used short-read technologies, which have difficulty sequencing templates with high GC content (Benjamini and Speed 2012). As a result, a substantial fraction of the bird microchromosomes have been almost entirely missing from the assemblies (Peona et al. 2018). Long-read sequencing technologies have led to a vast increase in high-quality chromosome-level assemblies for birds (Bravo et al. 2021). Such assemblies are available for 374 bird species on NCBI at the time of writing (NCBI 2026). While long-read technologies do not have the same GC bias as short-read ones, sequencing and assembling small bird chromosomes have remained difficult even with long reads (Peona et al. 2021; Barros et al. 2022). Indeed, the first bird genome assembly with all microchromosomes represented—that of a chicken (*Gallus gallus*)—was published only recently (Huang et al. 2023). This publication suggested that small chromosomes should be further divided into microchromosomes and *dot chromosomes*, the latter of which are the most extreme in terms of small size, high GC content, and enrichment for housekeeping genes (Huang et al. 2023).

Recently, analyzing the newly published T2T genome of zebra finch (*Taeniopygia guttata*), we found that non-B DNA motif coverage increases as chromosome size decreases and that introns in the euchromatic compartment of dot chromosomes have a particularly high non-B DNA enrichment (Formenti et al. 2026). Here, we expand these results and present a comprehensive analysis of the non-B DNA landscape in zebra finch, chicken, and several other bird species. For zebra finch, we study the distribution of non-B DNA in functionally important regions (e.g., different genic compartments) in detail, as well as in the sequences added to the T2T vs. the previous assembly (e.g., centromeres and repeats). We compare our results to the non-B DNA motif coverage in the nearly complete chicken genome (which is missing some telomeres and parts of the W chromosome (Huang et al. 2023), and potentially some genes (Ren et al. 2025)) as well as in seven other high-quality assemblies spanning ∼108 million years (MY) across the bird phylogeny (Kumar et al. 2017). Using Circular Dichroism (CD) analysis, an experimental spectroscopy technique, we examine the *in vitro* folding of the most common G4 sequences in the zebra finch genome. We further apply an *in vivo* technique, Permanganate/S1 Footprinting with Direct Adapter Ligation and Sequencing (PDAL-Seq, (Lahnsteiner et al. 2024)), to a zebra finch cell line, in order to detect the formation of single-stranded DNA (ssDNA) present at many non-B DNA structures genome-wide. Additionally, we analyze G4 ChIP-seq data, providing information about G4 folding *in vivo*, for chicken. Our results allow us to formulate hypotheses about functions of non-B DNA in bird genomes. Finally, we offer a potential explanation as to why the smallest bird chromosomes have been notoriously challenging to sequence.

## Results

### Non-B DNA motifs are distributed unevenly across the zebra finch genome, with contrasting coverage among chromosome categories

We annotated seven motif types with the potential to form non-B DNA structures in the recently available zebra finch diploid T2T assembly (Formenti et al. 2026) and the near-complete chicken assembly (Huang et al. 2023). These included A-phased repeats, direct repeats, short tandem repeats, inverted repeats, triplex motifs, G4s, and Z-DNA (Fig. 1A, see Methods). Compared to previous studies (Formenti et al. 2026; Smeds et al. 2025), here we applied novel algorithms for annotating G4s and Z-DNA (Mohanty et al. 2025; Petrovič et al. 2025), using scoring thresholds based on experimental data rather than solely on pattern match. We also used stricter thresholds for repeat and spacer lengths for the other motifs, specifically shortening the repeat and spacer length for inverted repeats (maximum lengths of 30 bp and 10 bp, respectively) (Brázda et al. 2016), and requiring 100% homopurine content and shorter spacer length (10 bp) for mirror repeats to be considered triplex-forming (Belotserkovskii et al. 1990). As a result, we conservatively focused on motifs most likely to form non-B DNA structures (see Table S1 for a detailed comparison of our results with those obtained with earlier methods). Taken together, the non-B DNA motif coverage across the genome was 7.6% for both zebra finch and chicken (Tables S1 and S2).

**Figure 1.**
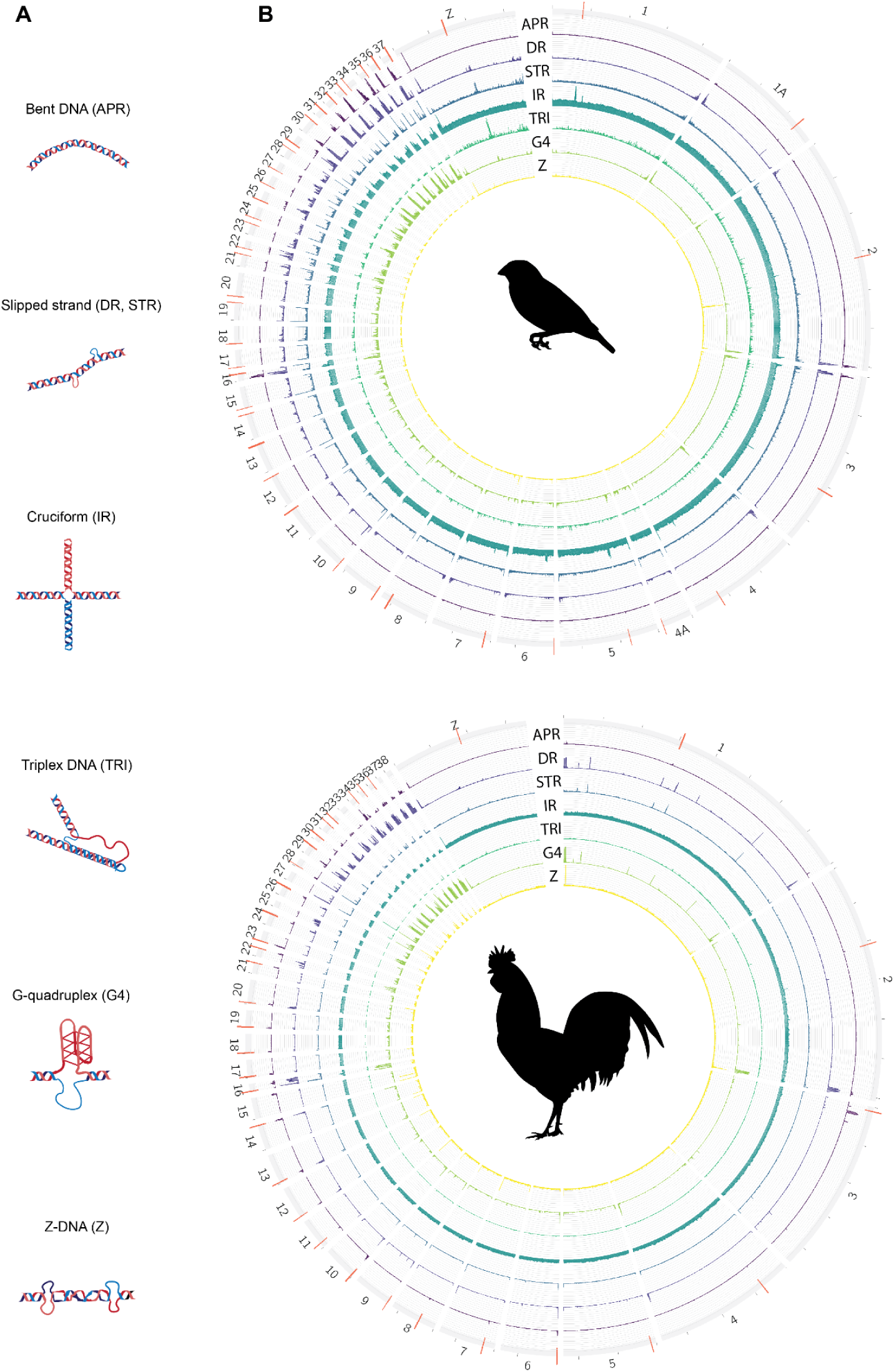
Examples of non-B DNA structures and their motif landscape in the zebra finch and chicken genomes. *(A)* Non-B DNA structures: bent DNA formed by A-phased repeats (APR), slipped-strand structures formed by direct repeats (DR) or short tandem repeats (STR), cruciforms formed by inverted repeats (IR), triplex DNA formed by homopurine mirror repeats (TRI), G-quadruplexes (G4) formed by guanine stems separated by loops, and Z-DNA (Z) formed by purine-pyrimidine stretches. *(B)* Circos plots with non-B motif coverage for the zebra finch primary haplotype (top) and the chicken assembly, which only has one haplotype publicly available (bottom). Abbreviations as in A. Bird silhouettes are from https://www.phylopic.org.

The zebra finch and chicken genomes consist of 11 and 10 macrochromosomes, respectively (including Z in both cases, and W in zebra finch), 19 microchromosomes, and 11 dot chromosomes (including W in chicken) (Formenti et al. 2026; Huang et al. 2023). The chromosomal distribution of non-B DNA motifs was, as we previously noted (Formenti et al. 2026), strikingly skewed towards high densities on small chromosomes, especially for the dot chromosomes, where an overall motif coverage was 21.6% for zebra finch and 33.7% for chicken (Figs. 1B, 2A, and S1, and Tables S1-S4), consistent with shorter and more GC-rich dot chromosomes in chicken than zebra finch. While this difference may be explained by the divergence time between the two species (∼90 MY (Kumar et al. 2017)), it should be interpreted with caution because chicken micro- and dot chromosomes still lack some telomeres and are therefore incomplete (Formenti et al. 2026).

**Figure 2.**
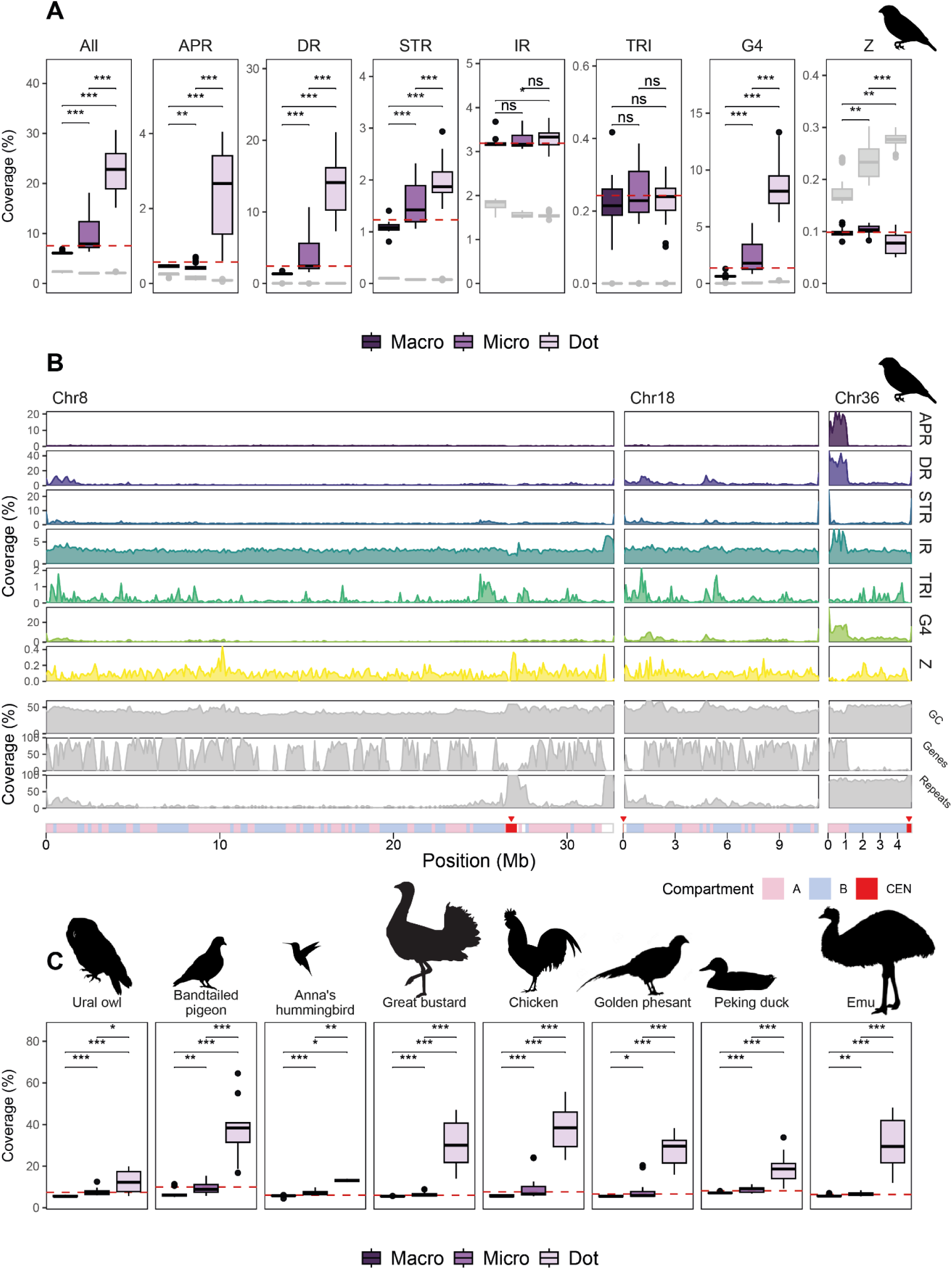
Non-B DNA motif abundance in different chromosome categories in birds. *(A)* Overall differences in various non-B DNA motif coverage among macro-, micro-, and dot chromosomes in the diploid zebra finch T2T genome. Shaded boxes show motif content in randomly shuffled chromosomes with the same GC content as the real chromosomes. “All” means all non-B DNA motif types taken together. \*\**P* <0.01, \*\*\**P* <0.001, Wilcoxon test for observed data only, adjusted for multiple testing using FDR. Comparisons marked with ‘ns’ are not significant. Note that the scale on the y-axis is different for each panel. *(B)* Examples of non-B DNA motif coverage in 100-kb windows along one zebra finch macrochromosome (chr8), one microchromosome (chr18), and one dot chromosome (chr36), along with GC-content, coding gene content, and repeat content. Maternal haplotypes were used in each case. The y-axis scale is different across motif types but is the same for the three chromosomes. The centromeres (red, marked with arrowheads) and compartments (pink: A compartment; and blue: B compartment) are shown below the panels. All chromosomes are shown in Fig. S3. *(C)* Overall non-B motif coverage in eight other bird species spanning more than 100 million years of evolution, comparable to “All” coverage plot for zebra finch in A. Statistical tests as in A. All bird silhouettes except the great bustard (drawn by the authors) are from https://www.phylopic.org.

While we expected to observe a particularly high coverage of G4s due to the high GC content at micro- and dot chromosomes (Fig. S2), we found that not only G4s but also A-phased repeats, direct repeats, and short tandem repeats had significant enrichment at dot chromosomes compared to macrochromosomes, whereas Z-DNA showed a significant depletion (Fig. 2A). To test to what extent different motifs are expected to occur at random in sequences with different GC content, we also annotated non-B DNA in simulated genomes where the nucleotides were shuffled around within each chromosome (Fig. 2A, shaded boxes). On dot chromosomes, for all motif types except Z-DNA, the observed motif content was considerably higher compared to the shuffled sequences with similar GC content. Microchromosomes sometimes had intermediate coverage when compared to macro- and dot chromosomes, e.g., for short tandem repeats, but often had coverage more similar to that of macrochromosomes (Fig. 2A). The non-B DNA coverage on the two zebra finch sex chromosomes—the Z and the W—was 6.5% and 6.9%, respectively, which was typical of the macrochromosomes (Table S1). Chicken Z and W had non-B DNA coverage of 6.1% and 23%, respectively, consistent with their classification as a macrochromosome and a dot chromosome, respectively (Huang et al. 2023) (Table S2). Thus, we observed substantial differences in non-B DNA motif coverage among chromosome categories, with starkest contrasts between macro- and dot chromosomes.

Visual inspection of non-B DNA motif landscapes in the zebra finch genome indicated high variability within chromosomes, with peak regions for different motif types sometimes colocated (Fig. 2B). Peaks were generally higher on dot chromosomes compared to macrochromosomes, with a tendency for Z-DNA motif peaks to be near some of the centromeres (Figs. 2B and S3). Interestingly, one high peak in the zebra finch pseudo-autosomal region (PAR, present on both Z and W), was enriched in all non-B DNA motif types except triplex motifs (Fig. S3, last panel). In dot chromosomes (e.g., chr36 shown in Fig. 2B; Fig. S3), A compartments, consisting of euchromatin (Fishman et al. 2019), were enriched in A-phased, direct and inverted repeats, as well as G4s, whereas B compartments, consisting of heterochromatin, were enriched in Z-DNA motifs. G4 enrichment was not always correlated with high GC-content (Fig. 2B, chr36).

To examine whether the skewed non-B DNA motif coverage between chromosome categories is a general feature of bird genomes, we repeated the analysis in seven other bird species with high-quality but yet incomplete genomes: Ural owl (*Strix uralensis*) (Chrysostomakis et al. 2025), band-tailed pigeon (*Patagioenas fasciata,* VGP genome), Anna’s hummingbird (*Calypte anna,* VGP genome) (Rhie et al. 2021), great bustard (*Otis tarda*) (Luo et al. 2023), Pekin duck (*Anas platyrhynchos*), golden pheasant (*Chrysolophus pictus*) (Li et al. 2025), and emu (*Dromaius novaehollandiae,* VGP genome). We chose these species based on assembly quality and phylogenetic position to achieve even spread across the bird tree (Fig. S4) (Kumar et al. 2017). All seven showed the same pattern as zebra finch and chicken, with dot chromosomes displaying the highest non-B DNA motif content (Figs. 2C, S5-S7, Tables S5-S11), strongly suggesting this is a general trend across the bird genomes. However, the absolute non-B DNA motif coverage varied between the added species, which could reflect true differences and/or differences in assembly quality/completeness (see Discussion).

We also assessed overlaps between non-B DNA motif types (i.e., when the same position was annotated as part of two or more motif types; Table S12, Fig. S8). In zebra finch and chicken, 15% and 18% of non-B-DNA-annotated bases overlapped at least two different motif types, and ∼5% overlapped more than two. Genomewide, the most common overlap was between G4s and direct repeats (3.5% of all non-B-DNA-annotated bases, or 5.8 Mb in zebra finch). Interestingly, almost 60% of all overlaps between G4s and direct repeats occurred on the dot chromosomes, even though the combined length of these chromosomes represented only ∼6% of the genome. Overlaps between different motif types in the other bird genomes were largely similar to those observed in the zebra finch and chicken (Fig. S8C-I).

### Non-B DNA sequences are enriched at several functional genomic regions

Non-B DNA motifs were enriched at functional regions in the zebra finch and chicken genomes (Fig. 3). Putative promoter regions (1 kb from the transcription start site, TSS) and 5’UTRs on macro- and microchromosomes showed moderate enrichment in almost all non-B DNA motif types in zebra finch (between 1.1-6.0× compared to genome-wide coverage; except for A-phased repeats) and most motif types in chicken (1.1-3.7×; Fig. 3A). The fold-enrichment was usually higher for microchromosomes compared to macrochromosomes, especially for G4s (6.0× vs. 3.9×, respectively, in zebra finch and 3.7× vs. 2.4×, respectively, in chicken). Dot chromosomes displayed a remarkably different pattern, with significant enrichment at intronic regions, which contained >9× of direct repeats and G4s in each species, as well as 8× of A-phased repeats for zebra finch compared to genome-wide densities (Fig. 3A, bottom panels). Chicken also showed high enrichment in intergenic regions, which could reflect that long non-coding RNAs are not yet annotated in the chicken assembly and are therefore included in the intergenic sequence. When comparing the functional regions to the chromosome category base level (for macro, micro, dot; see arrowheads on the rightmost part of Fig. 3A) rather than to the genome-wide average (Fig. 3A), the differences between categories were less pronounced.

**Figure 3.**
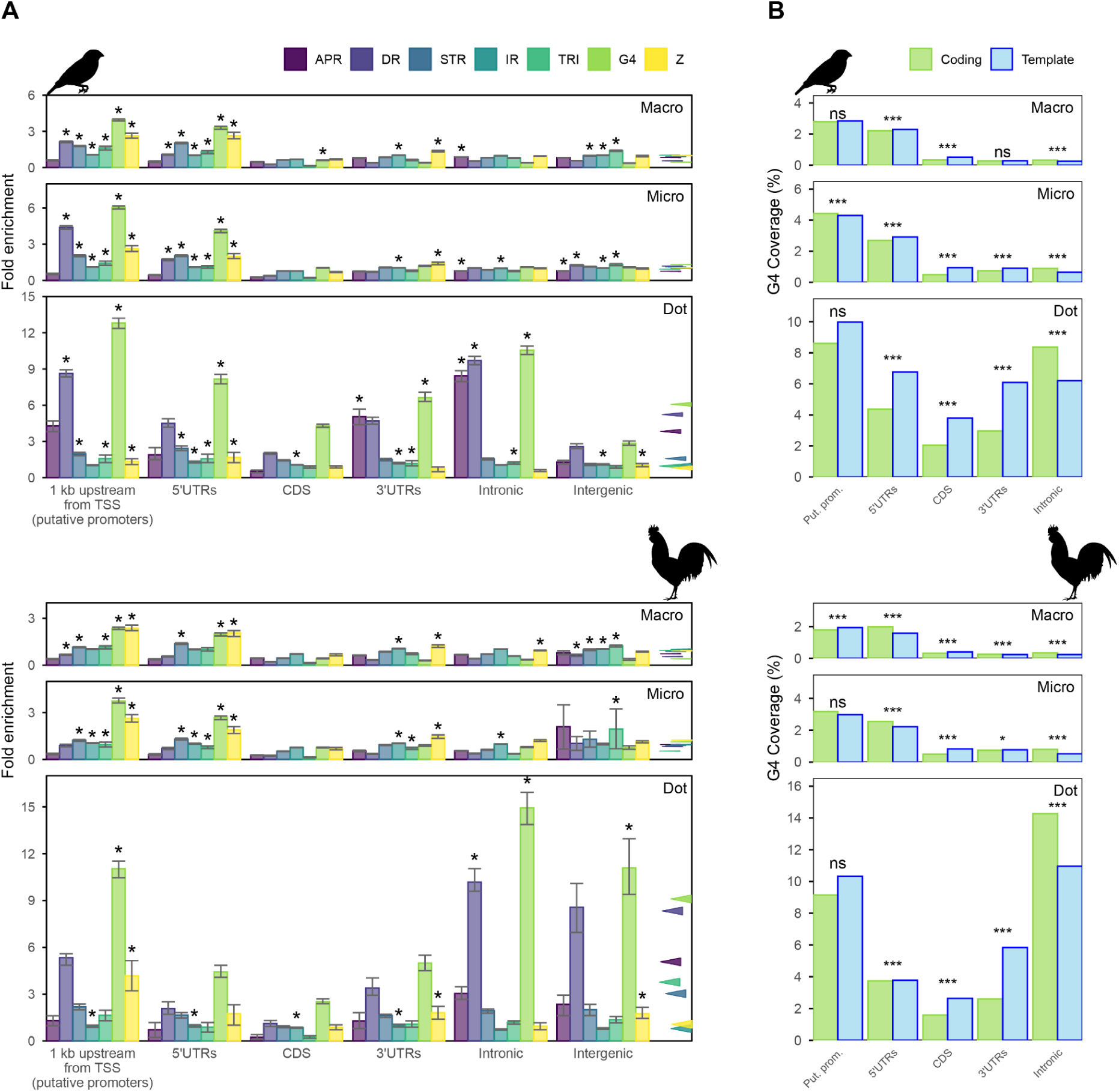
Non-B DNA motif enrichment at zebra finch and chicken functional regions. *(A)* Non-B DNA enrichment at functional regions on macro-, micro-, and dot chromosomes, as compared to genome-wide non-B DNA motif coverage. Average enrichment for the chromosome category (macro, micro, or dot) is shown with arrowheads to the right of the bars. For each bar, an interval was constructed by downsampling the data to 50% 100 times and excluding the two highest and two lowest enrichment values. Bars are marked with “*” if the interval is above the overall enrichment for that chromosome type (macro, micro, or dot; see Methods for details). *(B)* Coverage of G4 motifs on the coding and template strands. Bars are compared using a paired Wilcoxon test on the coverage per gene for each group, corrected with FDR; *** *P*<0.001, ** *P*<0.01, * *P*<0.05, ‘ns’ non-significant. Note that intergenic regions are strand-ignorant and therefore were excluded. Bird silhouettes are from https://www.phylopic.org.

We also assessed which strands the G4s preferentially occurred on, relative to the direction of gene transcription. For protein-coding regions (CDS) and 3’UTRs, there were significantly more G4s annotated on the template than on the coding strand in both zebra finch and chicken (Fig. 3B). However, the opposite was observed for introns. Overall, these observations suggest that G4s are less common at the level of the mRNA.

Despite differences in annotation completeness (Table S13), the non-B DNA enrichment patterns at functional regions in the other bird genomes were remarkably similar to those of zebra finch and chicken, with putative promoters and 5’UTRs enriched especially in G4s for all chromosome categories, and the most extreme enrichment levels on dot chromosomes, sometimes reaching 15-20× (Fig. S9). Dot chromosome introns were especially enriched in direct repeats and G4s—just as in zebra finch and chicken, but often to an even higher degree (up to 20× compared to genome-average for G4s in great bustard, Fig. S9G). Similar to chicken, direct repeats and G4s were enriched at intergenic regions of dot chromosomes in the other bird species, but not in zebra finch.

### Non-B DNA motifs are overrepresented at many tandem repeats and at some transposable elements and satellites

Different repeat classes in the zebra finch T2T genome assembly were analyzed for enrichment or depletion of non-B DNA motifs per chromosome category, in comparison to the whole-genome motif coverage (Fig. 4A). Among transposable elements (TEs), multiple classes of DNA transposons displayed enrichment in several non-B DNA motif types, particularly in direct repeats and G4s on dot chromosomes, though some TE classes (DNA/Kolobok, LTR/Copia) were entirely missing from these chromosomes (Fig. 4A, bottom panel). Among TEs examined, Ngaro elements—a distinct group of retrotransposons (Goodwin and Poulter 2004)—showed the highest relative enrichment, with Z-DNA motifs at approximately 300-fold above the genome-wide average coverage. However, Ngaro elements occupy only a minor part (∼70 kb) of the diploid zebra finch genome. We found that long interspersed nuclear elements (LINEs), short interspersed nuclear elements (SINEs), and long terminal repeats (LTRs) were usually not enriched in non-B DNA motifs (Fig. 4A).

**Figure 4.**
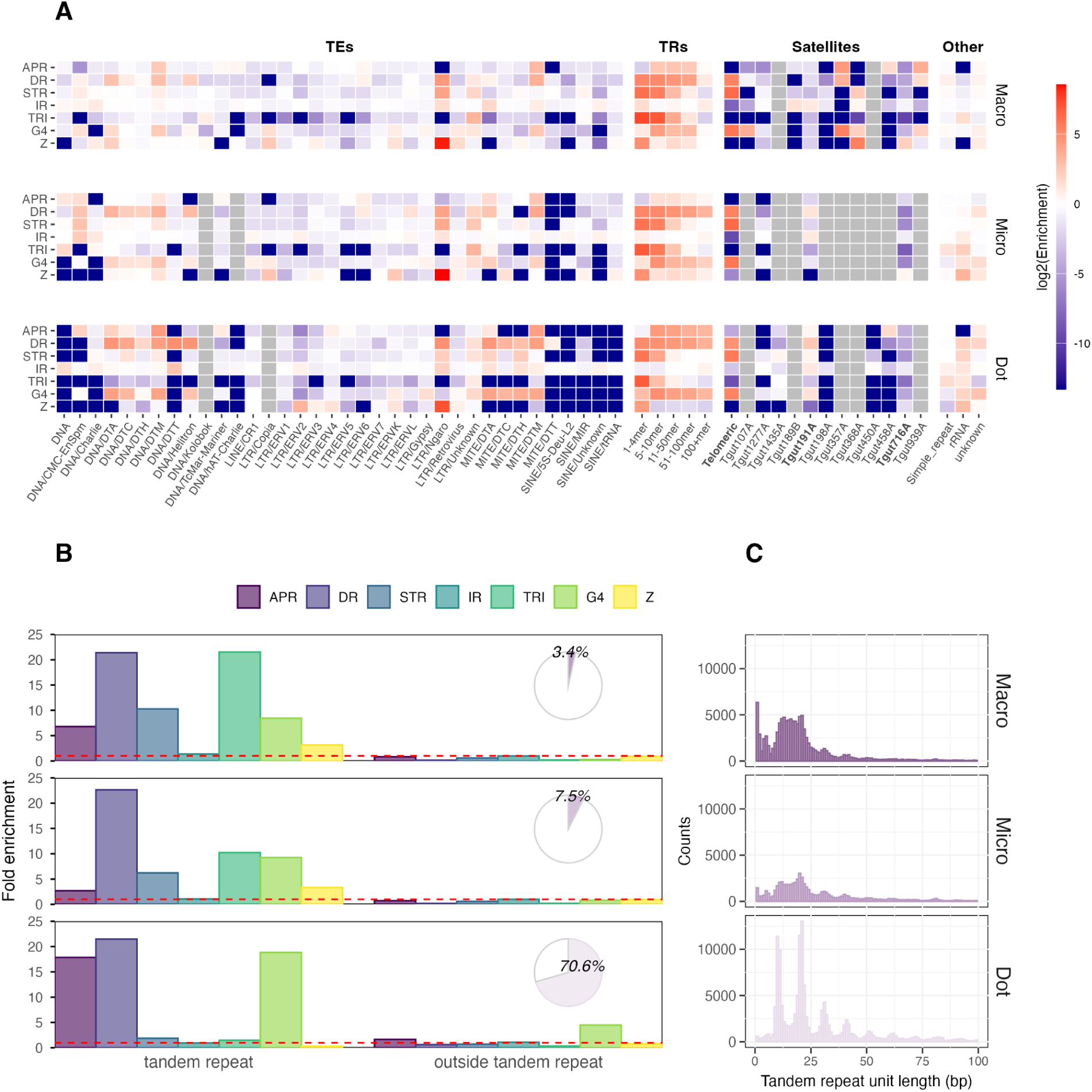
Non-B DNA motif enrichment in zebra finch repetitive regions. **A***(A)* Enrichment of non-B DNA motifs at different repeat classes for each chromosome category. Only repeat classes occupying at least 10 kb in the diploid genome are shown. TE: Transposable elements; TR: tandem repeats. White denotes no enrichment compared to the genome-wide average motif coverage; red and blue denote enrichment and depletion, respectively (color coding on the log scale). Repeat classes that are missing from a chromosome category are marked in gray. Telomeric and centromeric/pericentromeric satellites are marked in bold. *(B)* Enrichment of non-B DNA motifs within introns annotated vs. not annotated as tandem repeats. Inset pie charts show the proportion of introns that overlap with tandem repeats. The dashed red line denotes the genome-wide average. *(C)* Number of annotated tandem repeats in introns for each repeat unit length.

Tandem repeats of all lengths showed enrichment for essentially all non-B DNA motif types (Fig. 4A), and this enrichment was negatively correlated with the repeat unit length (Fig. S10). We further investigated overlaps between introns and tandem repeats. Intron regions annotated as tandem repeats showed notably higher enrichment for non-B DNA motifs than the rest of the introns (Fig. 4B). This enrichment was high across all chromosome categories. We observed that tandem repeats were highly abundant on dot chromosomes (constituting over 70% of introns) but rare on macro- and microchromosomes (constituting 3.4% and 7.5% of introns, respectively; Fig. 4B, inset). An exception was the high non-B DNA motif peak in PAR, which contained two protein-coding genes (*LOC140682075,* transcription factor 4-like, and *LOC140682130*, methyl-CpG-binding domain protein 2-like) with many tandem repeats in their introns (Fig. S11). The most common tandem repeats on dot chromosome introns were minisatellites, especially with repeated units of multiples of 10-11 bp (Fig. 4C).

The telomeres were enriched for direct repeats and G4s (as expected, because the telomeric sequence contains the repeated G4 motif). Satellites showed a highly mosaic pattern of enrichment and depletion in different non-B DNA motif types, with many low-frequency satellites enriched in A-phased repeats, direct repeats, and G4s, but often depleted in short tandem repeats, triplex repeats, and Z-DNA motifs (Fig. S12). One of the most common satellites, Tgut716A, was enriched in Z-DNA motifs on macrochromosomes (Fig. 4A). The other common satellite, Tgut191A, showed only slight enrichment in inverted repeats on macrochromosomes and in direct repeats on micro- and dot chromosomes. Both of these satellites were previously suggested to be associated with centromeres in the zebra finch (Takki et al. 2022), with Tgut716A identified as the dominant centromeric satellite repeat (Formenti et al. 2026). At the same time, Tgut191A appears to play a different role, potentially associated with Z-W conjugation in the PAR region (Formenti et al. 2026). The 18S/28S rRNA, almost exclusively present on dot chromosome pair 37 in the zebra finch (Formenti et al. 2026), was enriched in all motif types except for APRs. The 5S rRNA (named Tgut368A and shown in Fig. 4A), located in two clusters on macrochromosome pair 2 (Formenti et al. 2026), was highly enriched in Z-DNA, STRs, and G4s (45.3×, 4.9×, and 4.4×, respectively, compared to the genome-wide average).

In zebra finch, Z-DNA motifs were significantly overrepresented in six centromeres (2.9-19.9× compared to the genome-wide average; Fig. S13). The strongest overrepresentation was observed for macrochromosome 7, which likely drives the enrichment in satellite Tgut716A, as documented in the repeat analysis above (Fig. 4A). Because the average non-B DNA motif coverage differed strikingly among chromosome categories (Fig. 1B), we also assessed non-B DNA enrichment at each centromere in relation to the rest of the chromosome it was located on, rather than to the genome-wide average coverage, but this did not change the overall patterns (Fig. S14). However, we note that the choice of algorithm for annotating Z-DNA mattered. Using gfa’s pattern match of (RY)_5+_ utilized in several other publications (Gummadi et al. 2024; Wang et al. 2024; Yella and Vanaja 2023; Smeds et al. 2025) resulted in Z-DNA motif enrichment on 77 of 80 centromeres (Fig. S15). This is due to a large number of short (10-bp) purine-pyrimidine stretches that Z-DNA Hunter, which we used here, does not detect (it uses a minimum threshold of 12 bp) (Petrovič et al. 2025). Notably, over 70% of chicken centromeres showed enrichment in non-B DNA motifs, mainly A-phased repeats, direct repeats, short tandem repeats, and G4s, but not Z-DNA (Fig. S16).

### *In vitro* folding of G4s based on methylation and Circular Dichroism analyses

To assess whether G4s fold at functional regions in zebra finch, we used blood 5mC methylation levels as a proxy, because G4 formation is inversely correlated with methylation (Niu et al. 2025; Halder et al. 2010; Mao et al. 2018). Genic regions overlapping with G4s had significantly lower median methylation levels than genic regions not overlapping with G4s (Fig. 5A; methylation patterns for G4s located on the template vs. the coding strand were similar; Fig. S17). This pattern was most prominent at putative promoter regions and 5’UTRs, where the vast majority of CpG sites overlapping a G4 motif were unmethylated, and suggests that G4s can fold in these regions (Fig. 5A). Interestingly, for dot chromosomes, there was less methylation at 5’UTR G4s as compared to the putative promoter G4s, suggesting that their gene regulatory elements are closer to the TSSs, reflecting the compact nature of these chromosomes. We also observed that a significantly smaller proportion of 5’UTRs and putative promoters on the dot chromosome, as compared to macrochromosomes, have CpG sites at G4s, which can be methylated and hence interfere with G4 formation (Fig. S18). This is despite the high overall GC content on the dot chromosomes (Formenti et al. 2026). Therefore, our methylation analysis suggests that G4s are likely to fold and be involved in regulating functional genic regions in the zebra finch genome.

**Figure 5.**
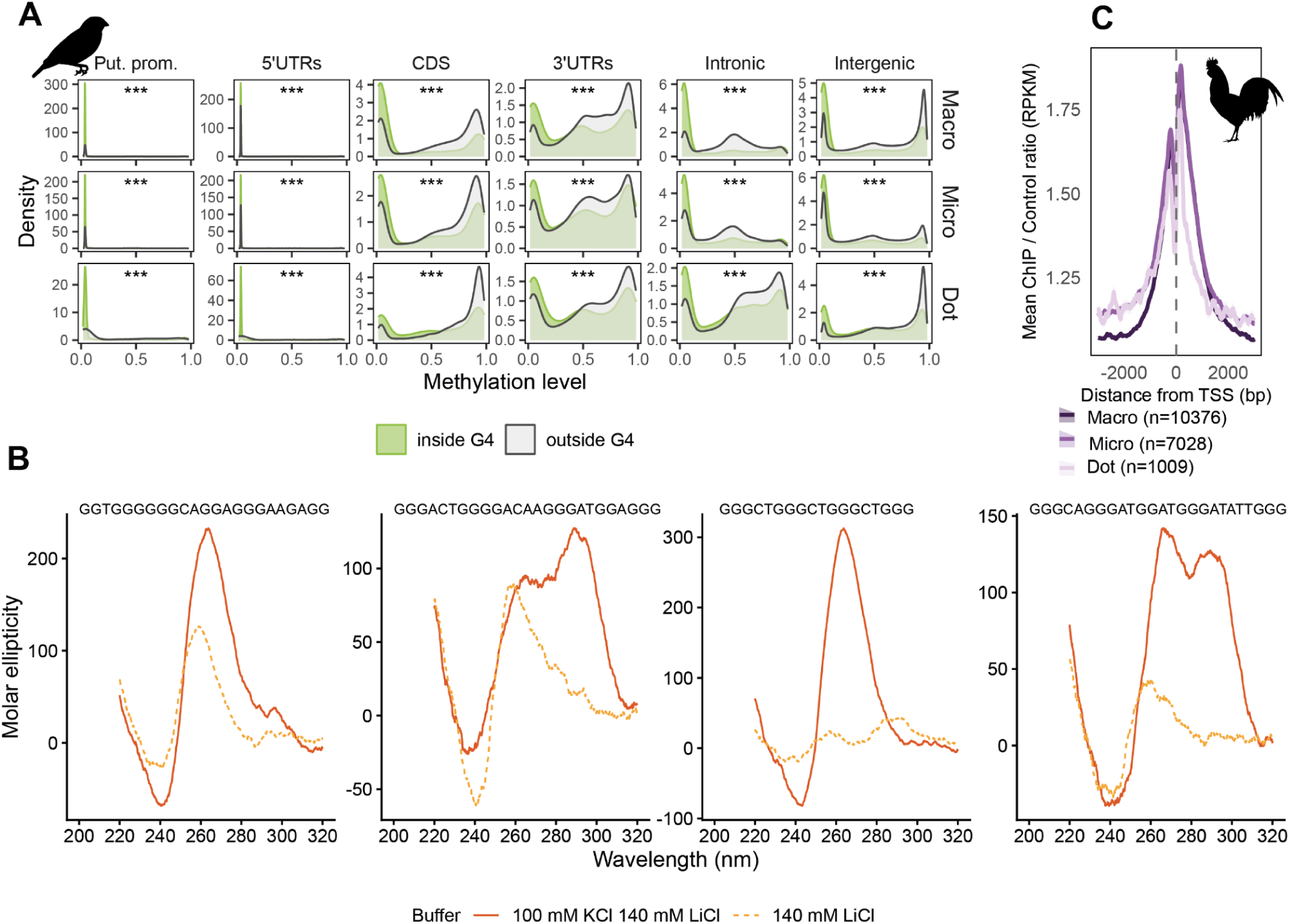
G4 formation. *(A)* Distribution of gene median methylation levels at CpG sites outside (gray) and inside (green) G4s in zebra finch. Gene regions are separated into chromosome categories, and G4s are strand-ignorant (for strand-specific G4s, see Fig. S17). *** *P*<0.001, Mann-Whitney U test. TSS: transcription start site; UTR: untranslated region, CDS: protein-coding region. *(B)* Circular Dichroism (CD) spectra for four commonly occurring G4 motifs in the zebra finch genome, listed in Table 1. *(C)* Ratio of G4 ChIP-seq data from chicken cell line DF1 compared to control around transcription start sites (TSS). Data taken from (Zheng et al. 2020). The number of genes per chromosome category is given in parentheses. Bird silhouettes are from https://www.phylopic.org.

**Table 1.**
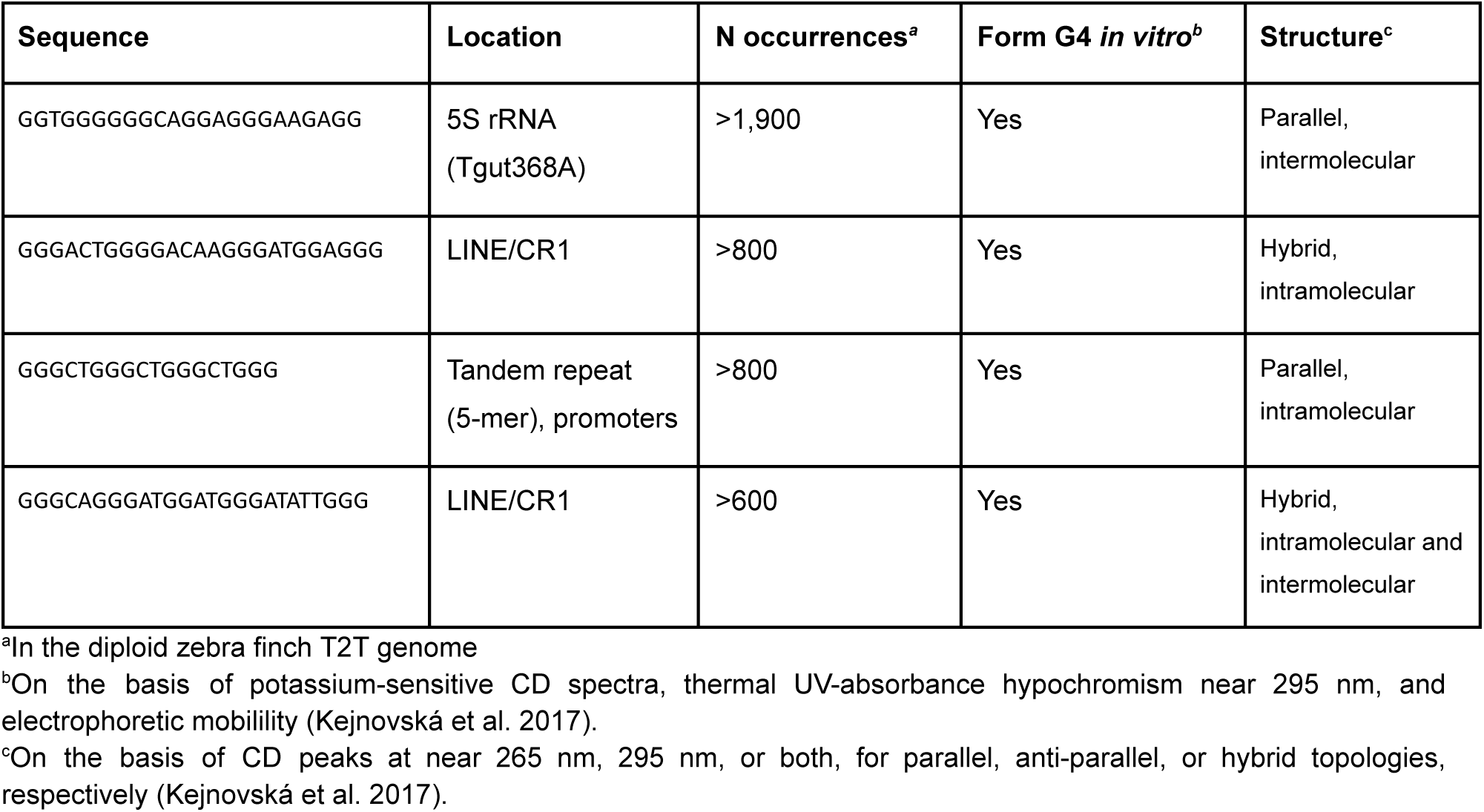
Experimentally verified common G4s in zebra finch.

We summarized the most common G4 motifs in the zebra finch genome (Table S14). The top two motifs, GGGTTAGGGTTAGGGTTAGGG (telomeric) and GGGAAGGGAAGGGAAGGG (in LTR retrostransposons on 73/80 chromosomes), occurring >69,000 and >33,000 times, respectively, in the diploid genome, have already been verified to form G4 structures *in vitro* in previous studies (Esnault et al. 2023; Bedrat et al. 2016; Del Villar-Guerra et al. 2018). We further tested four additional G4 motifs experimentally: one from the 5S rRNA (Tgut368A), two from LINE/CR1 elements (transposable elements unique to birds (Stumph et al. 1981)), and one from a tandem repeat that also had 10% overlap with putative promoters. All four motifs produced circular dichroism (CD) and UV-absorbance spectra that were consistent with known signatures of G4 structure (Mergny et al. 1998; Kejnovská et al. 2017) (Table 1, Figs. 5B and Fig. S19). Two of the tested G4 motifs also showed signs of multiple conformations on native gels (Fig. S20). These results indicate that these common G4 motifs can indeed form G4 structures *in vitro*.

### Formation of non-B DNA in cells

For chicken, we used previously published G4 ChIP-seq data targeting G4 structures with G4P, an artificial G4 probe protein (Zheng et al. 2020). ChIP-seq signal was enriched around the transcription start sites (TSS), indicating G4 formation in these regions, and we observed similar signal patterns across genes on chromosomes from different categories (Fig. 5C, the full heatmap of all genes is shown in Fig. S21). G4 ChIP-seq peaks were 23.8× and 51.3× more common at promoters and 5’UTRs, respectively, than genome-wide (see Fig. S22 for enrichment per class and chromosome category).

For zebra finch, we performed permanganate-S1 footprinting with direct adapter ligation and sequencing (PDAL-Seq). PDAL-Seq identifies loci prone to forming non-B structures in cells by sequencing the single-stranded DNA (ssDNA) associated with such structures (Lahnsteiner et al. 2024; Sieg et al. 2025). PDAL-Seq also detects transcription- and replication-induced ssDNA (Lahnsteiner et al. 2024; Sieg et al. 2025). Briefly, T and C residues at ssDNA loci were fixed by permanganate oxidation in a zebra finch cell line. The DNA was extracted, and ssDNA was subsequently digested by nuclease S1. We then ligated adapters to S1-cut sites and sequenced the DNA immediately adjacent to the original ssDNA site. Thus, PDAL-Seq identified ssDNA associated with loci that form diverse non-canonical structures in zebra finch cells (Fig. 6).

**Figure 6.**
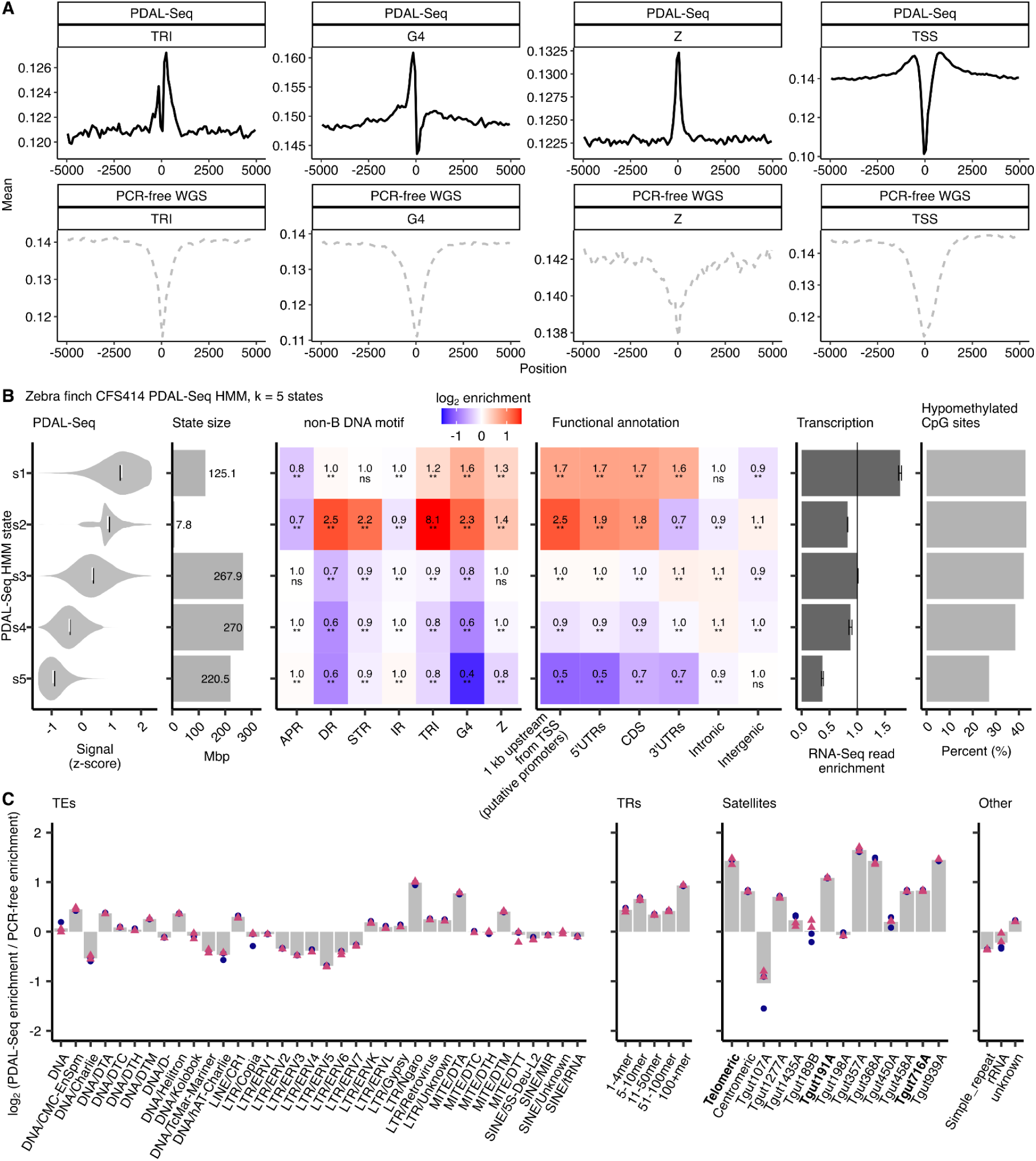
Association between non-B DNA motifs, single-stranded (ss) DNA, and genome function in zebra finch cells determined with permanganate S1 footprinting with direct adapter ligation and sequencing (PDAL-Seq). *(A)* Zebra finch CFS414 cell line PDAL-Seq mean signals around non-B DNA motifs and transcription start sites (TSS) in uniquely mapping, non-repetitive regions. PDAL-Seq data are compared to PCR-free whole-genome DNA sequencing data (WGS) to control for sequencing, mapping, and copy number artifacts. Mean read coverage (CPM per 100 bp bin, both strands summed) is plotted against distance from the feature center (bp). *(B)* CFS414 cell line PDAL-Seq signal in 1-kbp windows for uniquely mapping, non-repetitive regions of the zebra finch genome was fit to a *k* = 5 state Gaussian Hidden Markov Model (Table S15). Rows are hidden states, ordered by mean PDAL-Seq signal. Panels, left to right: z-scored PDAL-Seq signal per state with its mean (black line) and median (white line), x-axis were clipped at 2.5 and 97.5% quantiles; state size in Mbp; log_2_ enrichment of the seven non-B DNA motif classes; log_2_ enrichment of the functional annotations from Figure 3; CFS414 RNA-Seq read enrichment, the bar being the mean of the two libraries (sequence read archive accession numbers SRR17849680 and SRR17849681) and the error bar their spread; and the percentage of CpG sites in the state that are hypomethylated (0–20% 5-methyl-cytosine). Enrichment is observed coverage over a within-chromosome shuffled null, from 100 iterations of 10,000 subsampled annotations. Heatmap fill is clipped at ±log_2_(3) for color only; each tile is labeled with the unclipped enrichment above its significance mark (\**P* ≤ 0.05, \*\**P* ≤ 0.01, ns otherwise), where *P* is the Wilcoxon test above the subsample size and the empirical null below it. *(C)* ssDNA potential in repetitive regions of the CFS414 genome. Enrichment of PDAL-Seq compared to PCR-free whole-genome DNA sequencing at different classes of repeats sharing an x-axis with Figure 4A, except for centromeres which are included with satellites.

Analyzing PDAL-Seq data, we first asked how ssDNA formation varies at non-repetitive regions in zebra finch, and whether this variation was associated with non-B DNA motifs and genic functional elements (Sieg et al. 2025). PDAL-Seq signal was elevated at triplex, G4, and Z DNA motifs, as well as at TSS (Fig. 6A), consistent with non-B DNA structure formation. In contrast, PDAL-Seq signal was depleted at A-phased, direct, short tandem, and inverted repeat motifs, consistent with reduced ssDNA formation along with the presence of sequencing artifacts (Fig. S23). These results are concordant with permanganate/S1 studies in cell lines from mice, humans, and other apes (Kouzine et al. 2017; Sieg et al. 2025).

To further relate ssDNA formation to non-B DNA motifs and genic features genome-wide, we used a hidden Markov model (HMM) to segment the genome into data-driven states of PDAL-Seq signal while accounting for signal in neighboring windows (Yang et al. 2018; Ernst and Kellis 2017; Vu and Ernst 2022). We fit a Gaussian HMM to PDAL-Seq signals in 1-kbp windows after removing regions with high proportions of multimapped reads (Fig. 6A, see Methods). This approach segmented the genome by ssDNA formation and identified associations between PDAL-Seq signal, non-B motifs, and genic functional elements. The HMM state with the highest PDAL-Seq signal was ∼2-fold enriched in G4 motifs, promoters, CDS, UTRs, and RNA-Seq reads (Fig. 6B, State 1), consistent with G4 formation at transcribed genes (Esnault et al. 2025). The HMM also identified a rare high PDAL-Seq state that was strongly enriched in several non-B motifs (direct repeats, short tandem repeats, triplexes, G4s, and Z-DNA) along with promoters and 5’UTRs (Fig. 6B, State 2). These regions were less transcribed than those in State 1, but were hypomethylated, consistent with regulatory elements such as promoters and enhancers. In contrast, the two HMM states with the lowest PDAL-Seq signal were depleted in G4s and mostly consisted of intergenic regions (Fig. 6B, States 4-5). In summary, ssDNA profiles determined in cells confirmed that zebra finch non-B motifs are associated with ssDNA, transcriptionally active loci, and regulatory elements, suggesting that non-B DNA is implicated in gene regulation.

We next examined ssDNA formation in zebra finch repeats (Fig. 6C). We compared PDAL-Seq read enrichment with PCR-free, whole-genome sequencing read enrichment to control for sequencing and mapping artifacts across distinct repeat categories in zebra finch. Among TEs, the Z-DNA-enriched Ngaro element produced the strongest PDAL-Seq signal. Z-DNA produces ssDNA at single-stranded/double-stranded junctions (Kouzine et al. 2017), which may be the source of the ssDNA signal at these TEs. All tandem repeats were enriched in PDAL-Seq signal, indicating that these arrays of non-B motifs are forming noncanonical structures. Some satellites were highly enriched in PDAL-Seq signal, indicative of extensive non-B formation in cells. Telomeric satellites were over twofold enriched in ssDNA, consistent with the role of G4 structure in telomere maintenance (Monsen et al. 2021). Centromeric satellites were also enriched, driven by the ssDNA signal in Tgut191A and Tgut716A. The Tgut368A satellite, whose motifs form G4 structures *in vitro* (Table 1, Fig. 5B), also exhibited high ssDNA levels in cells. Interestingly, this satellite is enriched in both G4 and Z-DNA motifs. Future experiments should distinguish which structures actually form in cells.

### Non-B DNA motif landscape is associated with chromosome compartments in zebra finch

Because dot chromosomes differed significantly in non-B DNA motif coverage compared to micro- and macrochromosomes, we examined their motif distribution in zebra finch in greater detail. Bird chromosomes are partitioned into euchromatic A and heterochromatic B compartments, which have been delineated for the zebra finch genome (Formenti et al. 2026). On dot chromosomes, combined non-B DNA motifs were significantly more enriched in the A compartment than in the B compartment (Fig. 7A). A similar (but less pronounced) pattern was observed for macrochromosomes, but not for microchromosomes (Fig. S24).

**Figure 7.**
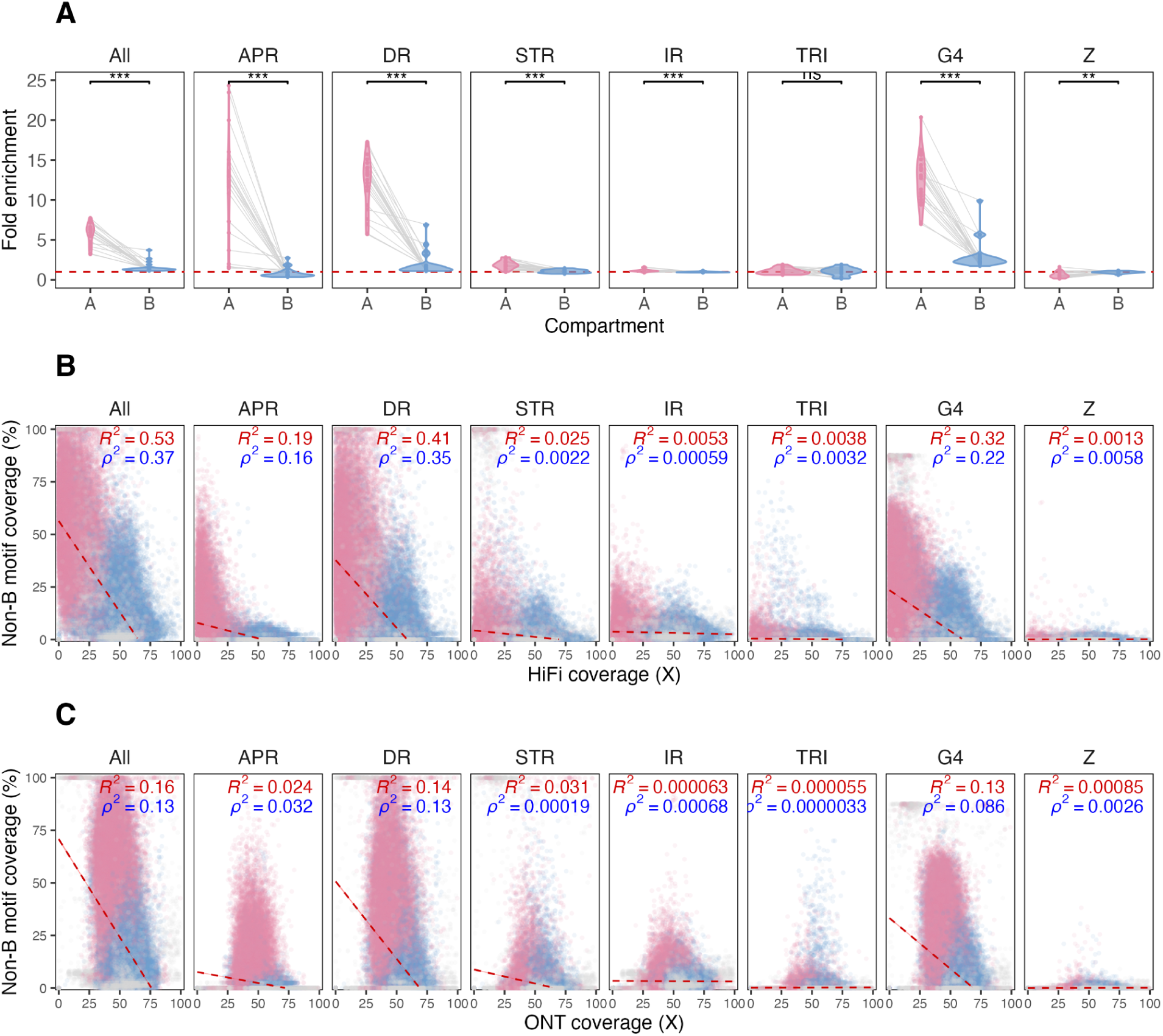
Non-B DNA motif enrichment and sequencing coverage at zebra finch dot chromosomes. *(A)* Enrichment of non-B DNA at the A compartments (euchromatic) and the B compartments (heterochromatic) for the 11 dot chromosomes in zebra finch (averaged across the compartments for each chromosome and haplotype separately) compared to the genome-wide average (red dashed line). Gray lines between violins connect the compartments from the same dot chromosome. \**P* <0.05; \*\*\**P* <0.001; Wilcoxon test adjusted for multiple testing using FDR. Note that some parts of the chromosomes were not assigned to A or B compartments; the telomeres and some satellites were excluded in the original annotations (Formenti et al. 2026). *(B)* Non-B DNA motif coverage and PacBio HiFi coverage on dot chromosomes in non-overlapping windows of length 1,024 bp, taken from (Formenti et al. 2026). Windows in A compartment are shown in pink, windows in B compartment are shown in blue, and windows not assigned to compartments are shown in gray. Red dashed lines show linear regression fits, and Pearson’s R^2^ (red) and Spearman’s ρ^2^ (blue) are shown for each correlation. Windows with sequence coverage greater than 100× are not displayed for visualization purposes. *(C)* Same as B but for ONT coverage data.

We next conducted a more detailed analysis of non-B DNA at A vs. B compartments on the dot chromosomes (Fig. 7A). We found that almost all non-B DNA motif types were more prevalent in A than B compartments, whereas Z-DNA showed the opposite pattern. A-phased repeats, direct repeats, and G4 motifs had particularly contrasting fold enrichment between A and B compartments. The high amount of G4s in the A compartment may be attributed to its high content of protein-coding genes, which in turn leads to an abundance of promoters and 5’UTRs, where G4s are enriched (Fig. 3A). Indeed, the A and B compartments had a gene coverage of 73% and 19%, respectively (exon coverage of 9.2% and 3.1%, respectively) (Formenti et al. 2026). Moreover, the A compartment had many minisatellites in the introns, in which A-phased repeats, direct repeats, and G4s were highly enriched (Fig. S25). A similar enrichment in non-B DNA motifs was also observed at intronic minisatellites in the B compartment, but these intronic minisatellites were much less prevalent in this compartment than in the A compartment (Fig. S25). In contrast, the B compartment contains many macrosatellites, including the Z-DNA-rich centromeric satellite Tgut716A (Formenti et al. 2026), which contributes to its overall Z-DNA enrichment. Note that there is no compartment annotation publicly available for chicken and other birds.

### Low sequencing depth on dot chromosomes is associated with high non-B DNA content

To test whether there is a link between non-B DNA and reduced sequencing depth on dot chromosomes, we evaluated the relationship between non-B DNA coverage and PacBio HiFi or Oxford Nanopore Technology (ONT) sequencing depth in 1,024-bp windows. This was performed separately for dot, micro-, and macrochromosomes. For PacBio HiFi, we found a negative relationship, which was particularly strong for dot chromosomes, with non-B DNA coverage explaining 53% of the variability in sequencing depth (Fig. 7B). When considered separately, the coverage of direct repeats and G4s explained 41% and 32% of the variability in sequencing depth, respectively. Similar, but much weaker, patterns were observed for micro- and macrochromosomes (Fig. S26). To investigate this further, we built a genome-wide multiple linear regression model with HiFi depth as response and non-B DNA coverage, repeat coverage, GC content, chromosome category, compartment, and GC content by non-B DNA coverage interaction (Table S16). In this model (R^2^=17.4%), chromosomal category and non-B DNA coverage were the strongest predictors, each explaining 8.1% and 3.9% of variance in HiFi sequencing depth. We also constructed separate models for each chromosomal category. Among them, the model for dot chromosomes explained the highest variance (R^2^=63.7%), followed by that for microchromosomes (R^2^=24.0%), with the model for macrochromosomes explaining the lowest amount of variance (R^2^=6.0%). In the dot chromosome model, non-B DNA coverage and chromosome compartment were the strongest predictors, each explaining 16.0% and 6.5% of variance. We also found a negative relationship between non-B DNA and ONT sequencing coverage (R^2^=16%; Figs. 7C and S27), but this was much less pronounced, especially in the multiple linear regression model (Table S17).

## Discussion

We conducted a comprehensive analysis of non-B DNA coverage in the zebra finch T2T genome (Formenti et al. 2026) and compared it to that of the second most complete bird genome, the chicken (Huang et al. 2023), as well as to those of seven other high-quality bird genomes (Chrysostomakis et al. 2025; Rhie et al. 2021; Luo et al. 2023; Li et al. 2025). The genomes of these nine species contained 6.1-10% of non-B DNA motifs despite more than 100 MY of divergence (Hackett et al. 2008). We note that these fractions may change when all these genomes (and not just zebra finch) are assembled to the T2T level.

Here, we used more stringent, conservative non-B annotations than previous studies (Smeds et al. 2025; Formenti et al. 2026) and selected motifs with the highest potential to fold. Notably, using more lenient definitions yields a non-B motif coverage of ∼11% in zebra finch and chicken, a level very similar to the non-B DNA motif coverage observed in ape T2T genomes (9.2-14.9%) (Smeds et al. 2025), which were also annotated with more lenient definitions. This is despite approximately 300 MY of divergence between mammals and birds (Hedges et al. 1996).

### Micro- and dot chromosomes

We found highly skewed non-B DNA motif coverage towards small (micro- and dot) chromosomes in all nine bird genomes analyzed here. Small chromosomes in bird genomes tend to have high GC-content, elevated recombination rates, high gene density in euchromatic A compartments, and high repeat content in heterochromatic B compartments (Huang et al. 2023). At the same time, small bird chromosomes appear to be dynamic, functionally important genomic units that require active regulation of transcription and recombination.

Dot chromosomes displayed particularly strong enrichment in non-B DNA, with the highest enrichment in their A compartment. The association between high recombination rates, GC-biased gene conversion, and open chromatin might provide a mechanistic explanation for the elevated non-B DNA content in A-compartments observed on dot chromosomes. Likely, non-B DNA helps maintain an ‘open’ state of these regions, allowing for efficient transcription of the many housekeeping genes that are located on dot chromosomes. Dot chromosomes also show unusually high interchromosomal interactions, as observed in Hi-C maps (e.g., (Waters et al. 2021; Formenti et al. 2026; Huang et al. 2023)), presumably because they are clustered in the interior of interphase nuclei (Habermann et al. 2001), and different 3D conformations of non-B motifs could facilitate these interactions (Tamon et al. 2025). Dot chromosomes experience a high recombination rate per unit length because each chromosome typically requires at least one crossover during meiosis. The fact that bird karyotypes are extremely conserved compared to, for example, mammals suggests that dot chromosome organization is under a selective constraint in avian evolution. This unique and ancient chromosome structure has been suggested to be key to bird and dinosaur survival (Griffin et al. 2026).

We have also observed the enrichment of A-rich motifs (i.e., A-phased repeats) on bird dot chromosomes. This pattern is largely explained by the abundance of minisatellite-like repeats that make up most of the dot chromosomes’ non-coding (i.e., intronic) euchromatic sequence. Minisatellites on dot chromosomes were recently observed in chicken, where euchromatic sequences were proposed to evolve via a local instability mechanism called sequence stuttering (Hron et al. 2026). Because these minisatellites contain multiple non-B motifs, they substantially increase non-B DNA motif enrichment for the euchromatic A compartment, whereas the heterochromatic B compartment typically has non-B DNA coverage close to genome-wide levels. Whether these minisatellites and the non-B DNA motifs within them have a function remains to be investigated, but, given the large phylogenetic distance between zebra finch and chicken, they appear to be important features of bird genomes, and our PDAL-Seq results suggest that they form non-B DNA structures.

### The role of non-B DNA in regulating bird genes

Our analysis indicates non-B DNA (particularly G4) enrichment at promoters and 5’UTRs in zebra finch and chicken (Fig. 3A), and G4 formation in the vicinity of TSSs in chicken (Fig. 5C). Moreover, our PDAL-Seq experiments in zebra finch found two elevated states enriched for several types of non-B DNA and for either genic functional elements or gene regulatory elements, such as promoters and 5’UTRs (Fig. 6B). Great ape promoters and 5’UTRs are enriched in non-B DNA motifs, particularly G4s and Z-DNA, as well (Smeds et al. 2025). This aligns with previous findings that G4s are important promoter elements (Esnault et al. 2025) and evolve under purifying selection at UTRs (Lee et al. 2020) in the human genome. G4s located at human promoter and UTR regions were also shown to be overrepresented, subject to purifying selection, and are predicted to form particularly stable structures (Guiblet et al. 2021b). This evidence points towards the functional role of non-B DNA at gene regulatory regions in both birds and primates, which is further strengthened by our *in vivo* analyses.

Building upon a previously noted negative correlation between G4 folding and methylation (Niu et al. 2025; Mao et al. 2018), we found low methylation, suggesting structure formation, at promoters and 5’UTRs in birds. This is consistent with observations for human and other great apes (Mohanty et al. 2025). Just as in great apes (Mohanty et al. 2025), bird protein-coding sequences, introns, and 3’UTRs display a bimodal distribution of methylated vs. unmethylated G4s. However, the zero methylation peak is taller with respect to the methylation peak in introns for birds than for great apes, which could suggest that a higher proportion of G4s fold in the former than in the latter.

Additionally, our results suggest that, compared to macro- and microchromosomes, dot chromosomes contain fewer CpGs at G4s in their regulatory regions. CpG sites can be methylated, preventing G4 formation. Thus, avoiding them ensures that these G4s always fold and the genes they regulate are consistently expressed. This agrees with the fact that dot chromosomes carry many housekeeping genes (Huang et al. 2023).

We observed that G4 motifs on most bird chromosomes are more prevalent on the template than the coding strand in UTRs and CDS, suggesting that G4 structures are avoided at the level of mRNA. This is consistent with previous observations in humans and other apes (Guiblet et al. 2021b; Mohanty et al. 2025; Huppert et al. 2008).

### Non-B DNA at repeats and centromeres

We observed a high enrichment for certain non-B motifs at specific bird TEs. The most striking enrichment in zebra finch was of Z-DNA motifs at Ngaro elements (Goodwin and Poulter 2004), which also have the strongest signal in our PDAL-Seq data, suggesting structure formation in cells. Non-B DNA has been suggested to play a role in the life cycle of some mammalian LINE-1s (e.g., (Sahakyan et al. 2017b)). Experimental validation *in vitro* of two highly abundant G4 sequences at zebra finch LINE/CR1, along with our PDAL-seq analysis, suggests that non-B DNA can form and play functional roles at repeats in bird genomes.

Non-B DNA was found to be enriched at bird short tandem repeats (i.e., micro- and minisatellites), which fold into slipped-strand structures, resulting in copy-number changes (Eckert and Hile 2009). This pattern likely drives high repetitive element content on dot chromosomes, which have specific minisatellite sequences located in introns and intergenic regions of euchromatin. The PDAL-Seq experiment suggested non-B structure formation in tandem repeats of all lengths, but we acknowledge that multi-mapping makes these regions challenging to study with short-read data, and thus requires further investigation. Intronic non-B DNA might be involved in alternative splicing (Georgakopoulos-Soares et al. 2022). Consistent with this, G4s were overrepresented on the coding strand of bird introns (Fig. 3B). Alternatively or additionally, intronic non-B DNA might facilitate recombination (Kshirsagar et al. 2017) and might represent recombination hotspots on compact dot chromosomes (Formenti et al. 2026).

While satellites (i.e., macrosatellites) as a class had low non-B DNA motif content, some particular satellites were enriched in non-B DNA. One abundant macrosatellite, Tgut716A, which is presently the strongest candidate for centromeric function in zebra finch, showed only some Z-DNA motif enrichment on certain chromosomes with the annotation method used in this study (see Methods), but significant enrichment on almost all chromosomes when annotated with gfa (Fig. S15), the software used for the non-B DB (Cer et al. 2011). The two methods differ in the purine-pyrimidine repeat length threshold used—12 bp and 10 bp. Our experimental results from PDAL-Seq suggest that non-B DNA structures do form at zebra finch centromeres.

We found chicken centromeres enriched in A-phased repeats, not in Z-DNA, as observed in zebra finch. This is not surprising, as no sequence similarity was found between chicken and zebra finch centromeres (Formenti et al. 2026), and centromeres are known to evolve rapidly (Henikoff and Furuyama 2010). Most chicken micro- and dot chromosome centromeres harbor a 41-bp tandem repeat called CNM, which often forms higher-order repeats (HORs), whereas macrochromosomes harbor different, chromosome-specific tandem repeats (Huang et al. 2023). The more precise centromere-defining CENP-A binding regions in chicken were not available for our study; however, they were shown to overlap with a short tandem repeat, sometimes identical to the telomeric sequence (TTAGGG)_n_ (Huang et al. 2023), likely explaining the G4 enrichment for some chicken centromeres.

Non-B DNA structures have been suggested to contribute to defining centromeres (Kasinathan and Henikoff 2018; Chittoor and Giunta 2024), to help maintain centromere position and integrity (Chardon et al. 2022), and to generate satellite copy number variation important for centromere drive (Talbert and Henikoff 2025). Z-DNA motifs have previously been shown to be enriched at centromeres in plants (Liu et al. 2023) and at specific chromosomes in human, chimpanzee, and bonobo (Smeds et al. 2025), but depleted at Drosophila centromeres (Patchigolla and Mellone 2022). Drawing general conclusions about the role of non-B DNA in bird centromere function will require more complete bird genomes, finer-scale analyses of centromeric components and the motifs they contain, and experimental tests of whether these motifs actually form non-B structures.

### Non-B DNA on the W chromosome

The fact that the highly repetitive zebra finch W (with repeat content of 86%, Table S18) did not show enrichment in non-B DNA can be explained by the fact that most of its repeats consist of TEs (Table S18), and TEs are in general not enriched in non-B DNA in the zebra finch. The notable exception—a peak enriched in all non-B DNA motifs except triplexes in the PAR region—had features similar to those of the peaks on the dot chromosomes, with minisatellites in the introns. PAR behaves similarly to dot chromosomes also in terms of its high recombination rate (Singhal et al. 2015). Satellite Tgut191A was highly abundant in the PAR on sex chromosomes and in the region between the telomere and euchromatin of microchromosomes. It showed no overall enrichment in non-B DNA motifs (Fig. 4A), but PDAL-Seq results suggest it forms ssDNA, indicating large slipped-strand or yet-undiscovered non-B DNA structures.

Chicken W shows higher non-B DNA motif enrichment than zebra finch W. This chromosome has not been assembled to the T2T level, so its non-B DNA motif coverage might still change. Among the other bird species considered, five have incompletely assembled W chromosomes and two lack W altogether, but the non-B motif coverage of the assembled sequence is more similar to that of zebra finch than to that of chicken, except for the golden pheasant. The golden pheasant belongs to the same order as chicken (Galliformes) and has largely conserved chromosome synteny with the latter, but its genome is also incomplete (Li et al. 2025).

### High non-B DNA content on dot chromosomes as a potential reason for their underrepresentation in older assemblies

The smallest chromosomes of bird genomes have been notoriously challenging to sequence and assemble (Formenti et al. 2026; Mathers et al. 2026). Dot chromosomes have a high GC content—a known problem for short-read Illumina sequencing, but a smaller problem for long-read PacBio and even less so for ONT sequencing (Ross et al. 2013; Browne et al. 2020). Therefore, the fact that dot chromosomes have been largely missing from bird assemblies until very recently (Huang et al. 2023; Formenti et al. 2026) may be partly explained by a shift from Illumina to long-read technologies. Additionally, non-B DNA structures, which we hypothesize form during sequencing, increase polymerase stalling for PacBio technology (Guiblet et al. 2018) and the speed of going through pores for ONT (Hosseini et al. 2023). Moreover, non-B DNA motifs increase sequencing errors, particularly for Illumina sequencing technology, but to a lesser extent for ONT and PacBio technologies (Weissensteiner et al. 2023). Even in the recent zebra finch genome sequencing effort (Formenti et al. 2026), we observed lower sequencing depths in euchromatic parts of the dot chromosomes, particularly with PacBio.

Examining this phenomenon genome-wide and for a complete T2T genome, we found that non-B DNA content explains a large proportion of dropout in PacBio HiFi sequencing depth, and a smaller proportion in ONT sequencing depth, particularly on the dot chromosomes. PacBio HiFi targets double-stranded fragments for sequencing, which become single-stranded (circular) after SMRT bell adapters attach, and are read repeatedly in a circular manner to produce high-confidence reads. The extreme density of non-B DNA motifs (especially G4s) in the euchromatic parts of dot chromosomes may result in many partial short single-stranded reads that cannot be captured by this technique. However, the high long-read sequencing depth used to generate T2T assemblies, and the combination of different long-read sequencing technologies—as was suggested (Weissensteiner et al. 2023)—alleviates some of the difficulties of sequencing through non-B DNA, and results in more accurate non-B DNA motif sequences and gapless genome assemblies, rescuing dot chromosomes.

These results are consistent with a small-scale study examining sequencing depth at five bird genes using Illumina and PacBio technologies (Beauclair et al. 2019) as well as a study investigating the cause of assembly gaps in birds-of-paradise (Peona et al. 2021), both of which suggested that non-canonical DNA structures may explain dropout in sequencing depth. Another recent study presents a tool developed for the Darwin Tree of Life assemblies to recover dot chromosome sequences from the unassigned scaffolds using conserved gene synteny (Mathers et al. 2026). The authors acknowledge that ONT works better than PacBio for birds, and since 2025, the Tree of Life project has used ONT to sequence bird and fish genomes (Oxford Nanopore Technologies, 2026). We recommend combining PacBio HiFi and ONT technologies to ensure a good representation of dot chromosome sequences in bird reference genomes.

## Materials and Methods

### Annotation of non-B DNA in the zebra finch and other bird genomes

The maternal and paternal haplotypes of the zebra finch T2T genome (GCA_048771995.1 and GCA_048772025.1 (Formenti et al. 2026)) were annotated for the following non-B DNA motifs: A-phased repeats (APR), direct repeats (DR), G-quadruplexes (G4), inverted repeats (IR), short tandem repeats (STR), mirror triplex motifs (TRI), and Z-DNA (Z). G4 motifs were annotated with the `G4DISCOVERY` pipeline (Mohanty et al. 2025), a hybrid approach that utilizes both `PQSFINDER` (Hon et al. 2017) and `G4HUNTER` (Bedrat et al. 2016) and keeps only motifs that are likely to form, based on experimental data. Z-DNA motifs were annotated using the `Z-DNA HUNTER` webserver (Petrovič et al. 2025) with Model=2, downloaded in a bedgraph format and converted to chromosome positions using a python script (https://github.com/makovalab-psu/T2T_bird_nonB/blob/main/python/remap_zdna.py). All other motifs were annotated with `GFA` (https://github.com/abcsFrederick/non-B_gfa (Cer et al. 2011)), with default motif parameter settings, which allow for spacer lengths up to 100 bp for mirror repeats and IRs, and 10 bp for DRs. It is debated whether motifs with long spacers actually form non-B DNA. After inspecting the spacer and repeat length distributions, which showed that most spacers in fact were shorter than 10 bp (Fig. S28), we filtered out motifs with spacers longer than 10 bp. For inverted repeats, we allowed the repeat arm length to be from 6-30 bp (threshold taken from `PALINDROME ANALYZER` (Brázda et al. 2016)), and for triplex motifs we only considered mirror repeats with 100% purine or pyrimidine content, as these are most likely to form triplex structures (Belotserkovskii et al. 1990). The output was converted to `BED` format using custom scripts (available on github). As an alternative approach, we assessed G4 with `QUADRON` (Sahakyan et al. 2017a) (using a dockerized version available at docker://kxk302/quadron:1.0.0) and Z-DNA with `GFA` and `ZSEEKER` (Wang et al. 2025). `QUADRON` and `GFA` are more lenient and find more motifs than `G4DISCOVERY` and `Z-DNA HUNTER`, but they overall showed a very similar motif distribution across the genome (Fig. S29). `ZSEEKER` found more motifs than `Z-DNA HUNTER`, but fewer than `GFA`. Conservatively, we decided to proceed with only the strict set of motifs. For some analyses all motifs were merged into a single track (named All) using `BEDTOOLS MERGE v2.31.0` (Quinlan and Hall 2010). The latest version of the near complete chicken genome (Huang et al. 2023) (downloaded from https://www.dropbox.com/scl/fo/plq2tm2w9lzlk0ua1rzph/h?rlkey=l6z3rgmjs7ec9azun8nundnzl&e=1&dl=0, including gene annotations), and genome assemblies for Ural owl (Chrysostomakis et al. 2025), band-tailed pigeon, Anna’s hummingbird (Rhie et al. 2021), great bustard (Luo et al. 2023), golden pheasant (Li et al. 2025), peking duck and emu (accession numbers GCF_047716275.1, GCF_037038585.1, GCF_003957555.1, GCA_026413225.1, GCA_037305975.1 (chr W from GCA_037216735.1), GCF_047663525.1 and GCF_036370855.1, respectively, downloaded from NCBI) were processed in the same manner.

### Chromosome synteny and assignment

Protein-coding gene sequences for eight species (there was no RefSeq annotation for golden pheasant) were extracted from the genome assemblies using `GFFREAD v0.12.7` (Pertea and Pertea 2020) and aligned to each other with `ORTHOFINDER v2.5.5` (Emms and Kelly 2019). For each species pair, homologous genes were summarized based on chromosome location, and chromosomes were assigned to macro-, micro-, or dot chromosome categories based on their gene homology to the zebra finch genome (maternal + Z), for which chromosome group assignments were taken from (Formenti et al. 2026). This grouping in zebra finch was based on structure rather than just size, so sometimes dot chromosomes were larger than the smallest microchromosomes. For chicken and golden pheasant, chromosome group assignments were given in the respective publication, and we decided to keep these assignments, which largely agreed with zebra finch (Huang et al. 2023; Li et al. 2025). The chromosome synteny across species is shown in Table S19.

### Non-B DNA coverage and enrichment at genes, repeats, and centromeres

For coverage plots, chromosomes were divided into non-overlapping 100-kb windows using `BEDTOOLS MAKEWINDOWS`, and overlap to each motif type was calculated using `BEDTOOLS INTERSECT` and custom scripts (https://github.com/makovalab-psu/T2T_bird_nonB/tree/main/python). Positions that were annotated for more than one motif (where different motifs overlapped each other) were only counted once per motif category. Hence, coverage is defined as the fraction of each window that is annotated as non-B DNA motifs. Circos plots were generated using `CIRCOS v0.69` (Krzywinski et al. 2009).

Per-chromosome coverage for all motifs was calculated as the sum of positions annotated as each motif type divided by the chromosome length. Coverage was also aggregated into three categories of chromosomes: macro-, micro-, and dot chromosomes, as well as genome-wide (diploid for zebra finch and haploid for the other species). These chromosome-, category-, and genome-wide coverages (as fractions of the regions) were used as baseline levels for different downstream enrichment calculations. For zebra finch, euchromatic (A compartment) and heterochromatic (B compartment) regions (defined in (Formenti et al. 2026)) in both haplotypes were downloaded as `BED` format for 200-kb windows from GenomeArk (https://genomeark.s3.amazonaws.com/species/Taeniopygia_guttata/bTaeGut7/manuscript/annotations/3D/bTaeGut7v0.4_MT_rDNA.Cooltools.E1.200kbp.flipped.dip.collated.v0.1.bed) and intersected with the non-B DNA motif annotations. Fold enrichment for A and B compartments compared to genome-wide coverage was calculated separately for each chromosome and haplotype, and the compartments were compared to each other using a Wilcoxon test, corrected for multiple testing with FDR.

To investigate the amount of expected non-B DNA motifs for different nucleotide compositions, the nucleotides of each zebra finch chromosome were randomly shuffled ten times (using the script https://github.com/makovalab-psu/T2T_bird_nonB/blob/main/python/shuffle_genome.py), keeping the overall chromosome GC-content unchanged. The ten shuffled genomes were annotated for non-B DNA motifs as described above, and per-chromosome coverage was calculated accordingly.

Different functional regions were extracted from the diploid zebra finch gene annotations (created with `EGAPX` (https://github.com/ncbi/egapx) and available at GenomeArk https://genomeark.s3.amazonaws.com/species/Taeniopygia_guttata/bTaeGut7/manuscript/annotations/genes/bTaeGut7v0.4_MT_rDNA.EGAPx.v0.1.gtf.gz) and haploid gene annotations for the other species (downloaded from NCBI for all species except chicken, that was downloaded from https://www.dropbox.com/scl/fo/plq2tm2w9lzlk0ua1rzph/h?rlkey=l6z3rgmjs7ec9azun8nundnzl&e=1&dl=0 and golden pheasant that lacks annotation). We used only the longest available isoform for each gene. UTRs were not included in the annotations and we extracted them by subtracting CDSs from exons using `BEDTOOLS SUBTRACT`, and defined them as 5’ or 3’UTRs depending on their position relative to the CDS and gene direction using a custom script (https://github.com/makovalab-psu/T2T_bird_nonB/blob/main/bash/05_functional_enrichment.sh). Promoters were not annotated, so we defined them as 1 kb upstream of each TSS (using only the longest isoforms of protein-coding genes). Introns were defined as the region between adjacent exons within the same gene (using the longest isoform), and overlaps with CDS, UTRs, and promoters (in overlapping genes) were removed using `BEDTOOLS INTERSECT`. Intergenic regions were defined as the regions between adjacent genes (including promoters), excluding the ends before the first and after the last gene on each chromosome. All regions were converted into `BED` format before downstream analysis. Each functional region was overlapped with each motif `BED` file using `BEDTOOLS INTERSECT`, and the sum of the unique overlapping positions was divided by the length of the region to obtain the non-B DNA motif coverage. Enrichment was calculated as region motif coverage divided by the genome-wide motif coverage. To test the robustness of the data, we randomly subsampled half of the region length for each functional class and chromosome category and calculated the enrichment for each subsample. This was repeated 100 times. The range of obtained enrichment values, excluding the two lowest and two highest values, was used to obtain error bars for each class and motif type. The error bars were compared to the chromosome category average enrichment, not to the genome-wide, to account for the different base-levels between the chromosomes and to reflect significant deviation from the rest of the chromosome. For G4s, enrichment was also calculated per gene based on whether they occurred on the coding or template strand, in relation to gene annotations. The content of coding and template G4s were compared in a paired Wilcoxon test, adjusted for multiple testing with FDR.

We used three sets of repeat annotations for zebra finch described in (Formenti et al. 2026)—which contain transposable elements, tandem repeats, and satellites, respectively. All annotations were downloaded from GenomeArk (https://genomeark.s3.amazonaws.com/index.html?prefix=species/Taeniopygia_guttata/bTaeGut7/manuscript/annotations/repeats/), converted into `BED` format, and labelled based on their classification using custom bash scripts (https://github.com/makovalab-psu/T2T_bird_nonB/blob/main/bash/04_repeat_enrichment.sh). Tandem repeats were grouped into the following length categories: 1-4-mers, 5-10-mers, 11-50-mers, 51-100-mers, and >100-mers, based on the length of a single repeat unit. Note that the three different repeat annotation sets overlapped significantly, as the same sequence frequently belonged to more than one annotated repeat type. Coverage and enrichment were calculated for each repeat category in the same manner as for genes, using the genome-wide coverage as the denominator. No repeat annotations were publicly available for the other bird genomes.

Centromeric annotations for zebra finch (defined in (Formenti et al. 2026)) was downloaded from GenomeArk (https://genomeark.s3.amazonaws.com/species/Taeniopygia_guttata/bTaeGut7/manuscript/ann otations/centromeres/bTaeGut7v0.4_MT_rDNA.centromere_detector.v0.1.gff) and converted to `BED` format. Centromere locations for most chicken chromosomes were provided by the authors (Huang et al. 2023). Enrichment in centromeres was calculated both in relation to the genome-wide coverage as well as to chromosome-wide coverage, the latter to account for inter-chromosomal differences and to test whether each centromere was enriched or depleted for non-B DNA motifs as compared to the rest of the chromosome. To test for significance, we divided the non-centromeric parts of each chromosome into 100 windows of the same size as the corresponding centromere. If there were more than 100 non-overlapping windows, 100 were selected randomly from the total set; if there were fewer, the windows were allowed to overlap.

For each window, the coverage and genome- and chromosome-wide enrichment were calculated, and these were used as a background distribution per chromosome to which the centromere enrichment could be compared in a two-sided randomization test. If the centromere enrichment was in the top or bottom 2.5% of the background distribution, it was considered significantly different from the chromosome average. No centromere information was available for the other species.

### Methylation analysis

Methylation data for zebra finch downloaded from GenomeArk was converted from bigwig to bedgraph with the UCSC software `BIGWIGTOWIG` (Kent et al. 2010). This file contains the percentage of 5-methyl-cytosine (5mC) methylated PacBio HiFi reads for each CpG site in the zebra finch genome. We extracted relevant sites in genes from each chromosome category using `BEDTOOLS INTERSECT` and separated them based on whether they occurred within G4 motifs or not. For each gene region, we calculated the median methylation level within or outside of G4s. Distributions of all medians were compared with a Mann-Whitney U test. We also investigated what fraction of genes had CpG sites (that could be methylated), for each chromosome category and gene class (within and outside of G4 motifs), and compared fractions from micro- and dot chromosomes to those of macrochromosomes using a Z-test, correcting for multiple testing with FDR.

### Experimental *in vitro* validation of zebra finch G4 sequences

We obtained single-strand DNA oligos from four common zebra finch G4 motifs from Integrated DNA Technologies (IDT) with high pressure liquid chromatography purification. We investigated structure using circular dichroism (CD) spectra, UV-absorbance thermal melting spectra ,and native gel electrophoresis, as described in (Kejnovská et al. 2017). Briefly, oligos were resuspended in nuclease-free water, and buffer exchanged into 140 mM LiCl, 20 mM LiMOPS pH 7.2 buffer, to remove trace small-molecule impurities. Concentrations were determined with Beer-Lambert law (Kejnovská et al. 2019) using the UV-absorbance at 260 nm and extinction coefficients provided by IDT. Samples were further diluted to a final 10 µM concentration in either 140 mM LiCl, 20 mM LiMOPS pH 7.2 or the same buffer with 100 mM KCl. Samples were denatured at 95°C for 1.5 min and slowly annealed to 4°C over 90 min. Samples were stored at 4°C for subsequent analysis. CD spectra were collected at 20°C in 0.1 cm cuvettes using a Jasco J-1500 spectrometer. Background CD from the buffer was subtracted from every wavelength for each sample, and molar ellipticity was calculated using the formula E = CD/(33000 × molarity × pathlength in cm). UV-absorbance thermal melting spectra were collected on an OLIS HP 8425 diode array spectrophotometer from 5 to 95℃ at a ramp rate of 0.5℃/min, collecting the absorbance spectra every 0.5℃. Background absorbance from the buffer was subtracted from each sample and absorptivity was calculated using Beer’s law. Conformational heterogeneity was assessed by native gel electrophoresis, where samples were fractionated on a 15% polyacrylamide gel containing 10 mM KCl and 1× Tris Borate EDTA (TBE) buffer for about 3 hours at 150 V and 4℃. Bands were visualized using SYBR gold stain on a Bio-Rad Gel Doc Go imager. Molecular weights were compared to a New England Biolabs Low Molecular Weight dsDNA ladder (catalog number N3233).

### G4 ChIP-seq validation of chicken G4 sequences

Previously published ChIP-seq data from the chicken embryonic fibroblast cell line DF1 that targeted G4 structures using an artificial G4 probe protein (Zheng et al. 2020), was downloaded from SRA (accessions SRR9603969 and SRR9603970), and processed following (Qian et al. 2024). In brief, the data was trimmed with `TRIMGALORE v2.2.0` (Krueger 2026), mapped to the chicken reference with `BOWTIE2 v2.5.2` (Langmead and Salzberg 2012), and sorted and deduplicated with `SAMTOOLS v1.19.2` (Danecek et al. 2021). The enrichment compared to the control sample was assessed with `BAMCOVERAGE` from `DEEPTOOLS v3.5.6` (Ramírez et al. 2016), using RPKM normalization, and the TSS profiles were generated with `COMPUTEMATRIX` from the same package. G4 peaks were called using `MACS3 CALLPEAK v3.0.4` (Zhang et al. 2008). Peak enrichment was calculated by dividing the peak density of a region to the genome-wide or chromosome-category density.

### Permanganate/S1 footprinting with direct adapter ligation (PDAL-Seq) data collection and analysis

Zebra finch CFS414 fibroblast cells were obtained from the Antibody and Bioresource Core at Memorial Sloan Kettering Cancer center. Cell culture reagents were ordered from Gibco. Cells were grown in Dulbecco’s Modified Eagle Medium (high glucose) supplemented with 10% non-heat inactivated fetal bovine serum, penicillin, streptomycin, glutamax, non-essential amino acids, and sodium pyruvate in T25 or T75 flasks. Fixed cells were rinsed with PBS, suspended with 0.25% trypsin-EDTA for 7 min, rinsed, and seeded into fresh media between passages during expansion. Cells were grown at 37℃ with a 5% carbon dioxide atmosphere until five 40% to 60% confluent in T75 flasks (50-70 million cells). Duplicate 20 and 40 mM permanganate-treated PDAL-Seq libraries were prepared as described previously (Lahnsteiner et al. 2024). A PCR-free, whole-genome-sequencing library was also prepared from untreated DNA with a NEBNext® Ultra™ II FS DNA PCR-Free E7430 Library Prep Kit for Illumina (New England Biolabs) according to manufacturer’s specifications. Libraries were sequenced on a NextSeq 2000 using a P2 kit with paired-end 2 x 150 bp sequencing kit. Because PDAL-Seq localizes a single-stranded-DNA–derived ligation point at the 5′ end of the forward read, only R1 was analyzed.

PDAL-Seq data were preprocessed, validated, associated with fit to HMMs, and analyzed in repetitive regions as described previously (Sieg et al. 2025). Briefly, a single haplotype reference was constructed by retaining maternal chromosomes and chromosome Z. Reads were trimmed with `FASTP v0.24.0` (-p -P 50 -y -x -3) (Chen et al. 2018), mapped with `BWA MEM v0.7.18` (Li 2013), sorted with `SAMTOOLS SORT v1.21` (Li 2013; Danecek et al. 2021), de-duplicated with `SAMTOOLS RMDUP` and -s. Two 20 mM and two 40 mM replicates had high, genome-wide correlation and were merged. For analysis of PDAL-Seq signals in uniquely mappable regions, sex chromosomes, mitochondrial DNA, rDNA morphs, and 10 kbp windows that lost more than 10% of their signal to multimappers were blacklisted. We quantified hidden Markov model (HMM) segmentation of PDAL-Seq signal in 1 kbp windows, normalized it to the summed genome-wide coverage total, and converted it to a z-score using the mean and standard deviation across windows. A variational Gaussian HMM (hmmlearn vhmm VariationalGaussianHMM, covariance_type=“full”, n_iter=2000, fixed random_state=42) was fit to the windowed z-scores. A k = 5 state HMM was chosen because it robustly reflects functional associations between PDAL-Seq signals and annotations using the minimum number of states (Figure S23B). Enrichment was computed by a subsample-and-shuffle procedure implemented with `BEDTOOLS v2.31.1` (Quinlan and Hall 2010) and defined as mean(observed) divided by mean(null) over the 100 iterations. Significance was assigned with a Wilcoxon rank-sum test for annotations with more than 10,000 loci. For smaller annotations, an empirical p-value was computed as the fraction of null iterations deviating from the null mean by at least as much as the observed mean. Publicly available RNA-Seq reads (SRR17849680 and SRR17849681, (Biegler et al. 2022)) were trimmed with the same `FASTP` settings and mapped with `STAR v2.7.11b` (Dobin et al. 2013). Per-state 5-methy-cytosine rates were computed with Pacific Biosciences long reads (Formenti et al. 2026).

All genomic regions and reads were retained when determining the enrichment of PDAL-Seq signals in repeats. Repeat annotations were intersected with a 1 kbp window grid, yielding “fragments”. For each library × repeat class, and for each of 100 iterations, 1,000 fragments were subsampled (all fragments were retained for classes with fewer) and enrichment was calculated. The observed statistic was the summed per-base read coverage over the subsample. The null statistic was the same summed per-base read coverage after the subsample was shuffled with `BEDTOOLS SHUFFLE` (-chrom flag included). Enrichment was mean(observed) divided by mean(null). The final statistic was reported for each satellite class as the ratio of Enrichment(PDAL-Seq library) / Enrichment(PCR-free control), because a repeat class can accumulate or deplete reads due to sequencing or mapping artifacts.

### Coverage analysis in relation to non-B DNA motifs

PacBio HiFi and ONT coverage for zebra finch (in wig format, averaged for 1024-bp windows) were downloaded from GenomeArk (https://genomeark.s3.amazonaws.com/index.html?prefix=species/Taeniopygia_guttata/bTaeGut7/assembly_verkko_0.1/manual_curation/bTaeGut7v0.4/mapping/) and converted to `BED` format using `WIG2BED` (Kent et al. 2010). For each window, the non-B DNA motif content was calculated for each motif type, and the correlation between sequence coverage and non-B content was assessed for each chromosome category using a linear model in `R` (R Core Team 2024). Pearson’s R^2^ and Spearman’s ρ^2^ were calculated in `R` (R Core Team 2024) using the package `GGPUBR` (Kassambara 2026). The linear mixed models for impact on sequencing coverage were set up as SeqCov ∼ NonBFrac + RepFrac + GCFrac + Compartment + [Category] + (1 | Chr) using the R package `LMERTEST` (Kuznetsova et al. 2017). We ran both a full model with chromosome category as a categorical predictor, and three separate models for each chromosome category (macro, micro and dot). Windows without compartment information were excluded from all analyses. To assess the impact of predictors we performed a dropout analysis by removing one predictor at the time from the model and recorded the change in marginal R^2^. A separate model was run to test for the interaction between Non-B DNA and GC (nonBFrac × GCFrac). The values were first normalized around their means (value_norm = value - mean(value)) to make them comparable, and the change in marginal R^2^ was calculated as the difference to the original model without interaction. All figures were prepared in `R` unless otherwise indicated, using the libraries `TIDYVERSE` (Wickham et al. 2019), `GGPLOT2` (Wickham 2016), `GGUPSET` (Ahlmann-Eltze 2026), `COWPLOT` (Wilke 2025), `VIRIDIS` (Garnier et al. 2024), and `GGH4X` (van den Brand 2025).

## Data access

The non-B DNA motifs in bed format are available for download from Zenodo, doi 10.5281/zenodo.19225047, and all the code for this manuscript is available on GitHub: https://github.com/makovalab-psu/T2T_bird_nonB, and archived on Zenodo https://doi.org/10.5281/zenodo.19934862. PDAL-Seq sequencing reads were deposited on the National Institutes of Health National Center for Biotechnology Information Sequence Read Archive accession number PRJNA1530990. The full PDAL-Seq data analysis pipeline is available at https://github.com/makovalab-psu/ZebraFinch_PDAL-Seq.

## Acknowledgements

We thank Kaivan Kamali for the dockerized version of Quadron and Luohao Xu for centromere coordinates in the chicken genome assembly. We are grateful to Matthias Weissensteiner for providing thoughtful comments on an earlier version of this manuscript. This research was supported by the grant R35GM151945 and by the Willaman Chair Endowment Fund from the Eberly College of Science to KDM. Computations were performed at the Penn State Institute of Computational Data Sciences (RRID:SCR_025154), which provided access to computational research infrastructure within the Roar Core Facility (RRID:SCR_026424). Sequencing was performed by Ashley Price at the Penn State Huck Institutes’ Genomics Core Facility (RRID SCR_02645).

## AI statement

JPS used Claude Code Opus (Anthropic) to adapt the pipeline from Sieg et al. (2025) to the zebra finch PDAL-Seq samples. LS used Claude Sonnet 5 (Anthropic) to replicate the pipeline from Qian et al. (2024) for chicken G4P GhIP-seq data, and for improving R code.

