## Supplementary Figures S1-S29 for "Non-canonical DNA and sequencing challenges in bird genomes"

**Figure S1.** Non-B DNA coverage on the zebra finch alternative haplotype

Circos plot of zebra finch alternative haplotype (paternal excluding Z chromosome) with non-B DNA motif coverage in 100-kb windows. Each motif type is scaled from 0 to the maximum coverage for that type (meaning absolute bar heights are not comparable between the motif types). Centromeres are marked with red bars in the karyotype. Abbreviations: APR: A-phased repeats; DR: direct repeats; STR: short tandem repeats; IR: inverted repeats; TRI: triplex motifs; G4: G-quadruplexes; Z: Z-DNA. Bird silhouette is from https://www.phylopic.org.

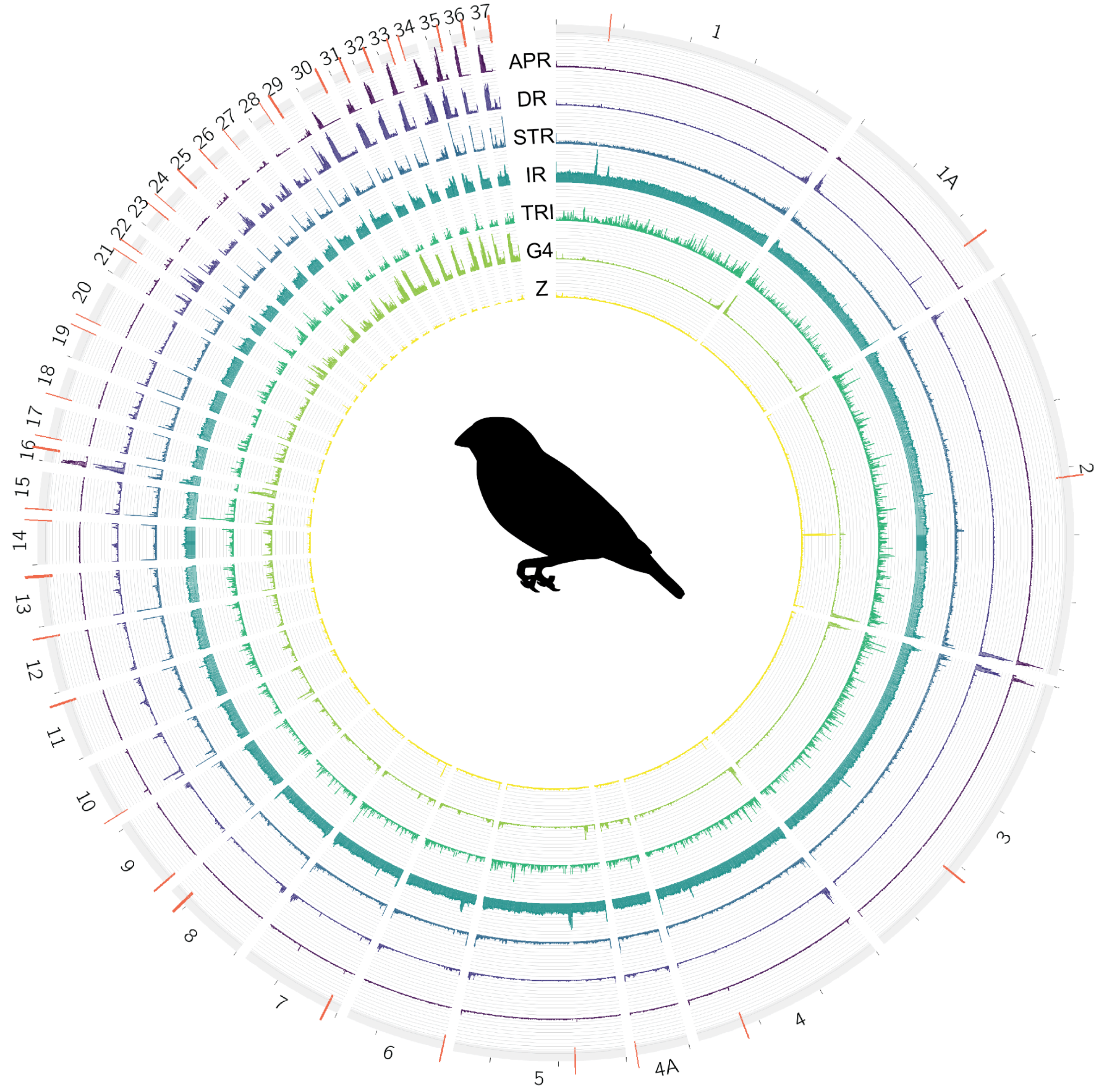

**Figure S2.** Non-B DNA motifs and GC content

Non-B DNA motif content in relation to GC content for *(A)* zebra finch and *(B)* chicken. Both maternal and paternal chromosomes are included for zebra finch. “All” means all motifs taken together. The outlier dot chromosome visible for chicken APR, STR and TRI is the W chromosome. Abbreviations are as in Fig. S1. Bird silhouettes are from https://www.phylopic.org.

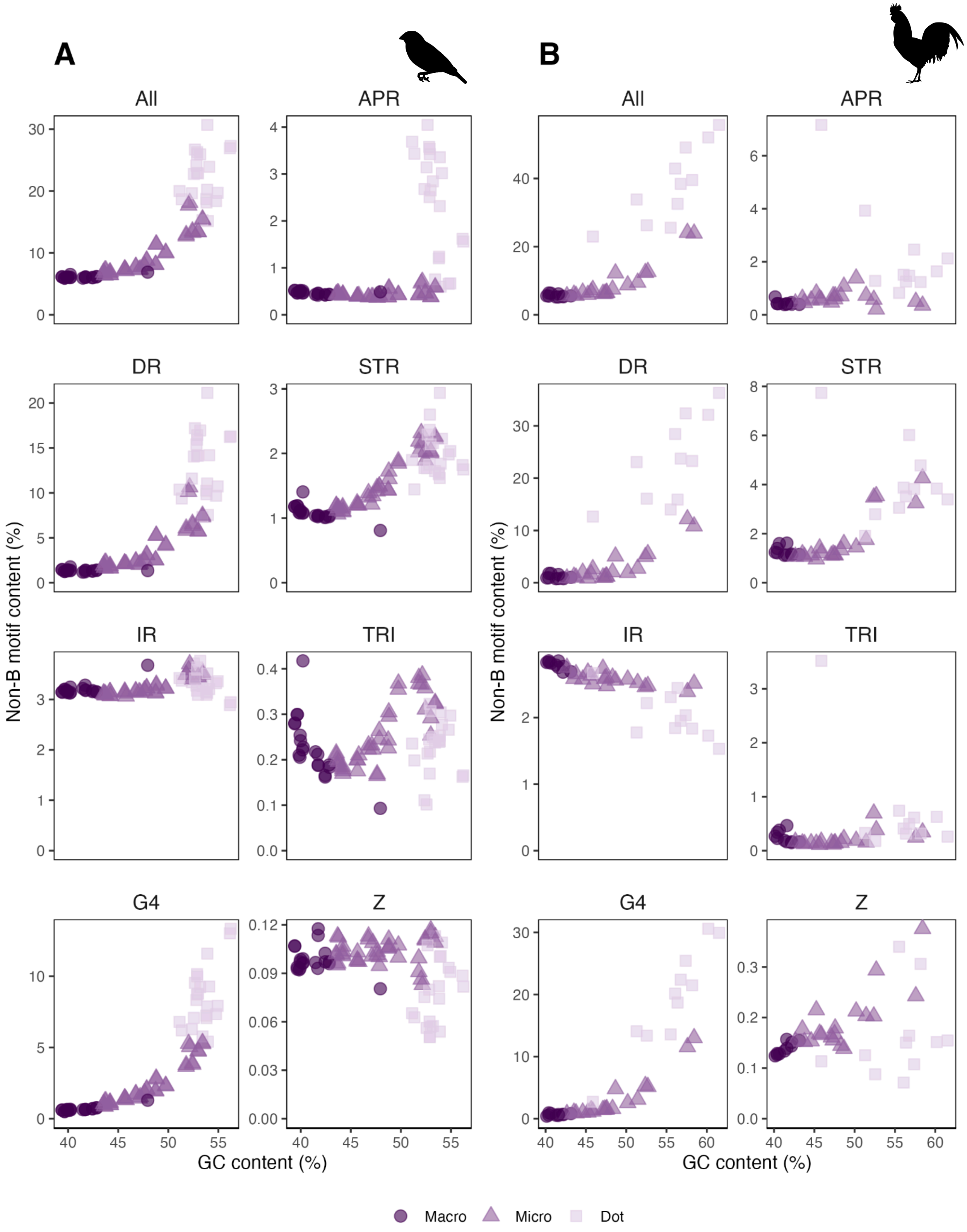

**Figure S3**. Non-B DNA motif coverage along zebra finch chromosomes

Same as Fig. 2B, but for all chromosomes. A and B compartments are shown below in blue and pink, respectively; and centromeres are marked in red, with an arrowhead above.

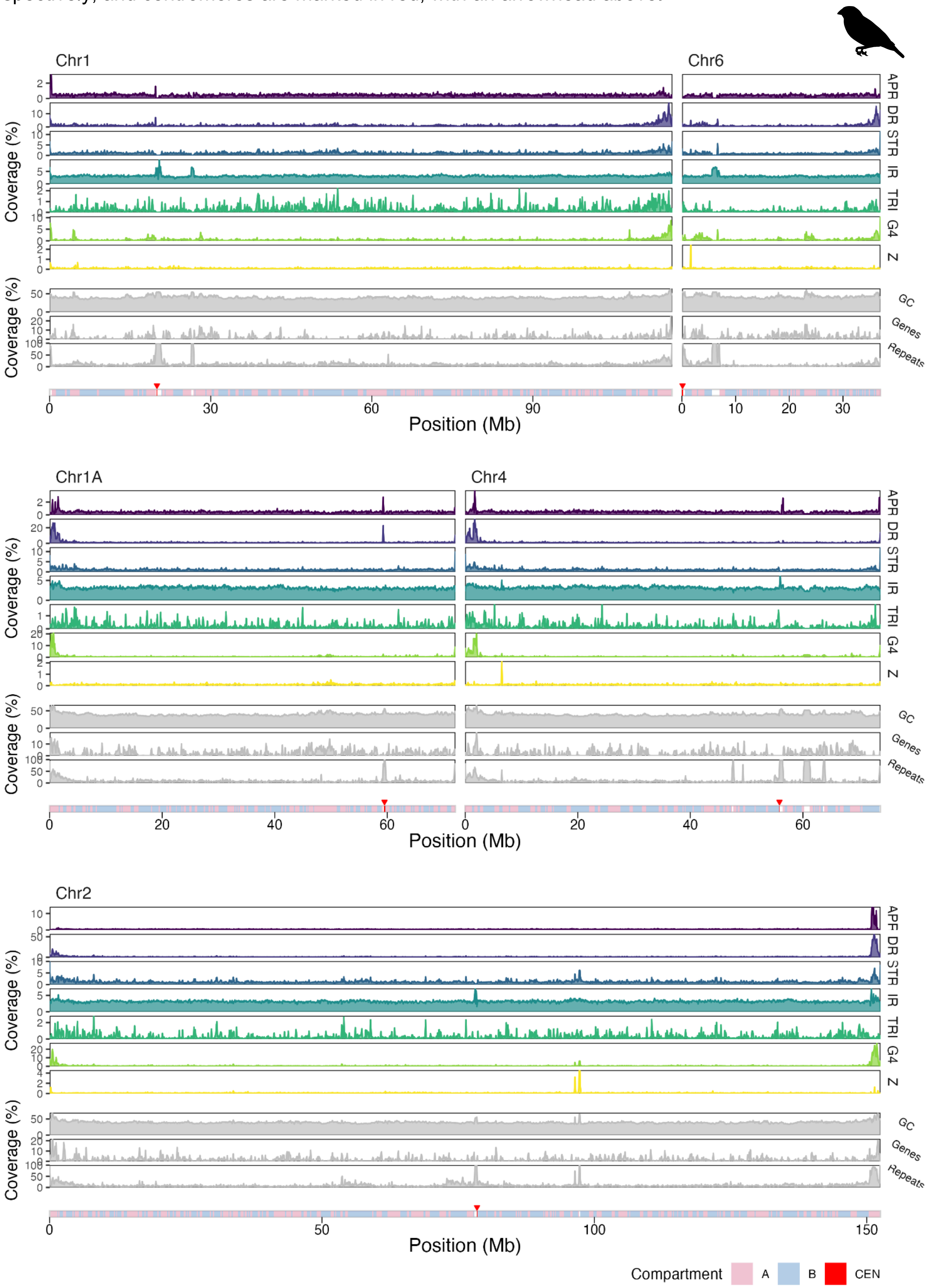

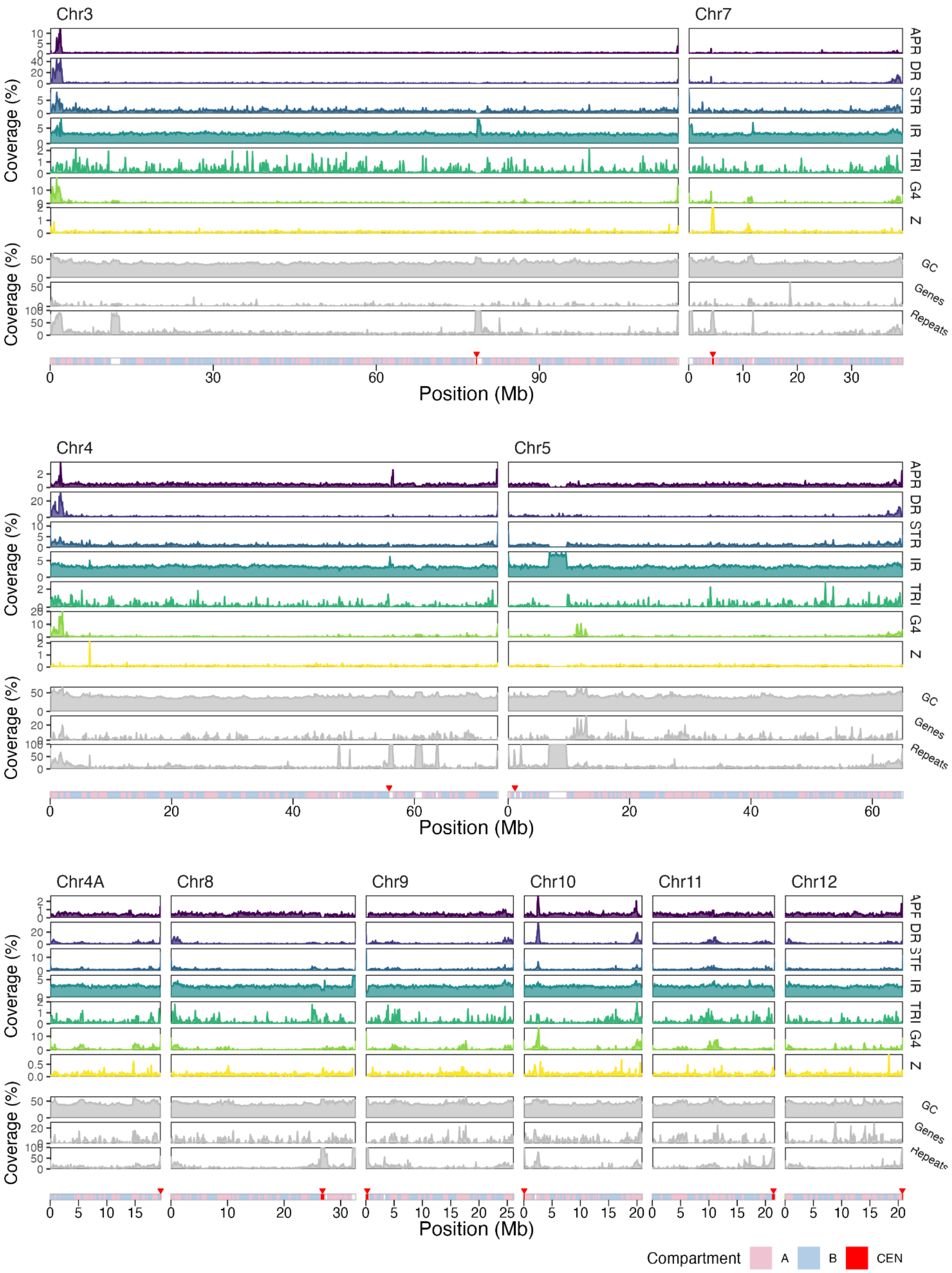

Chr3
Chr7
Chr4
Chr5
Chr4A
Chr8
Chr9
Chr10
Chr11
Chr12
APR
DR
STR
IR
TRI
G4
Z
GC
Genes
Repeats
Coverage (%)
Position (Mb)
Compartment
A
B
CEN

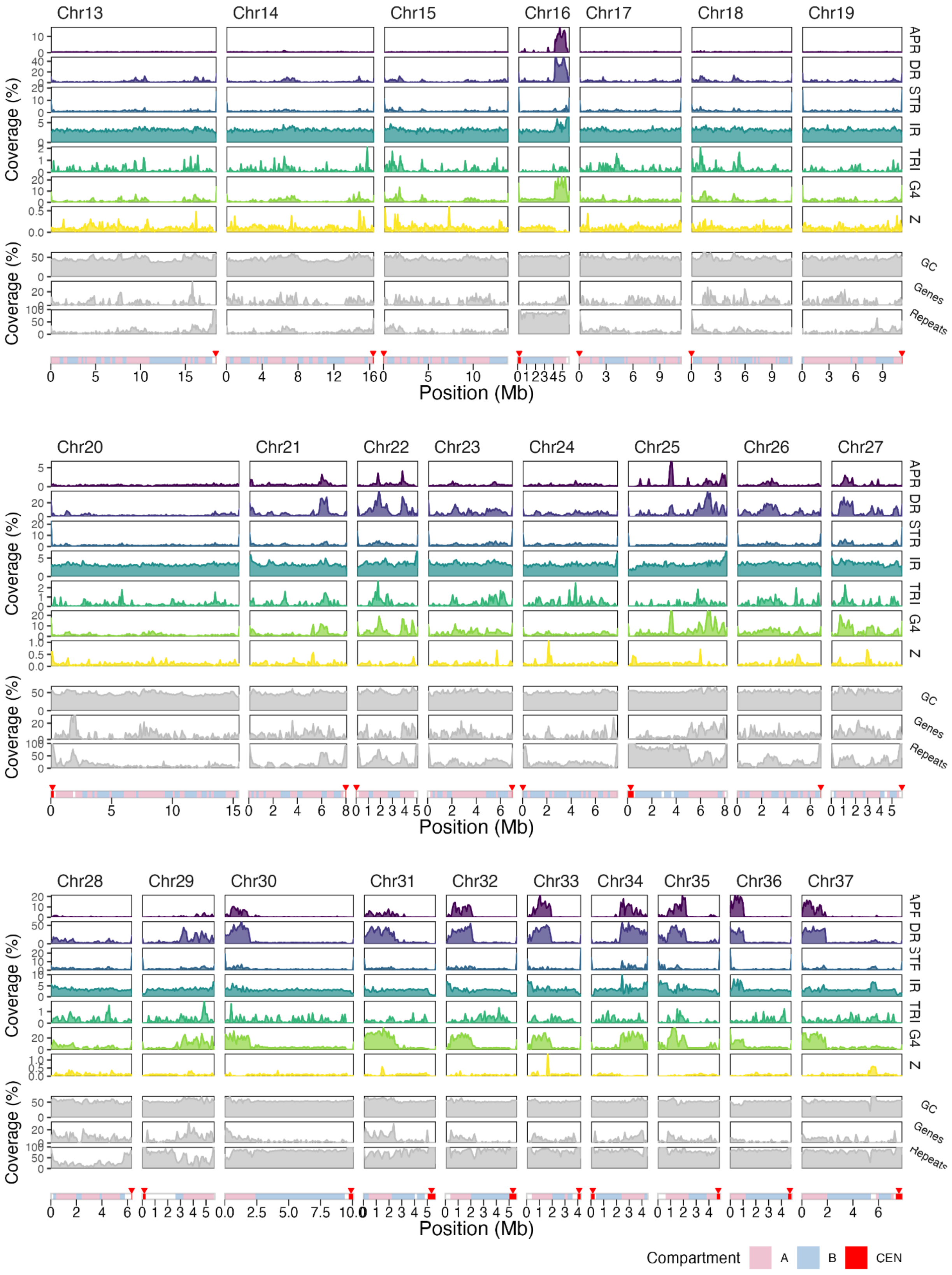

Chr13
Chr14
Chr15
Chr16
Chr17
Chr18
Chr19
Chr20
Chr21
Chr22
Chr23
Chr24
Chr25
Chr26
Chr27
Chr28
Chr29
Chr30
Chr31
Chr32
Chr33
Chr34
Chr35
Chr36
Chr37
Coverage (%)
APR
DR
STR
IR
TRI
G4
Z
GC
Genes
Repeats
Position (Mb)
Compartment
A
B
CEN

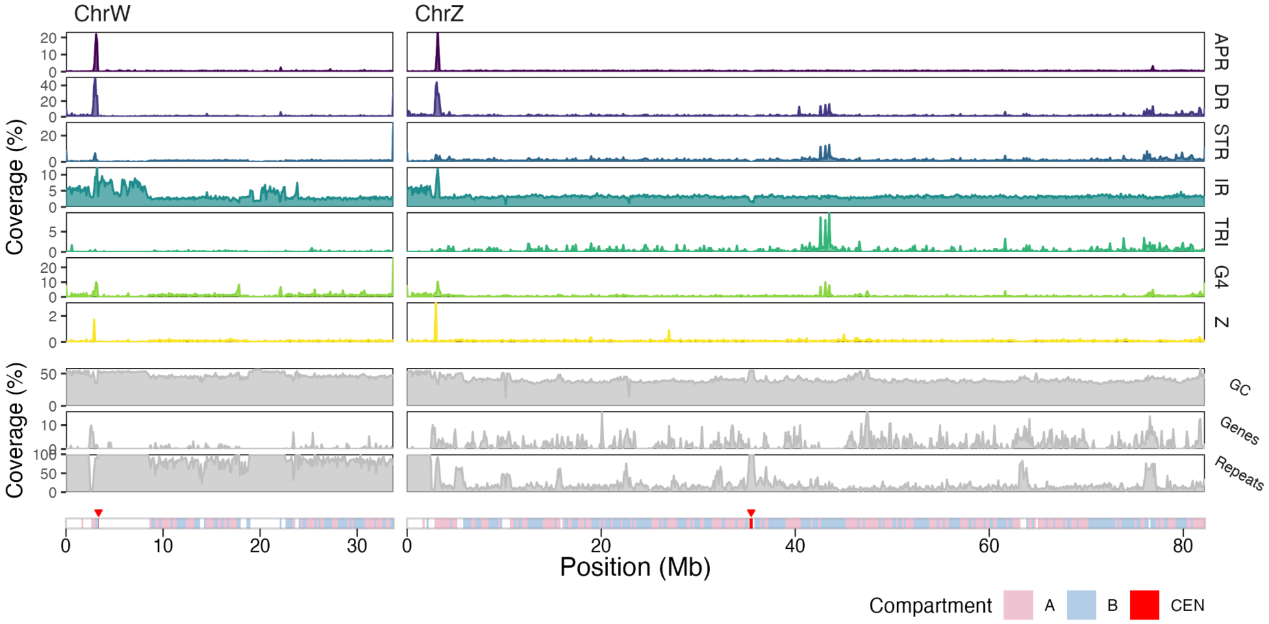

ChrW
ChrZ
APR
DR
STR
IR
TRI
G4
Z
GC
Genes
Repeats
Coverage (%)
Position (Mb)
Compartment
A
B
CEN

**Figure S4.** A schematic evolutionary tree for the nine species
Generated with timetree.org. The great bustard was replaced with another bustard (*Chlamydotis undulata*) due to the lack of representation of the genus *Otis* on timetree.

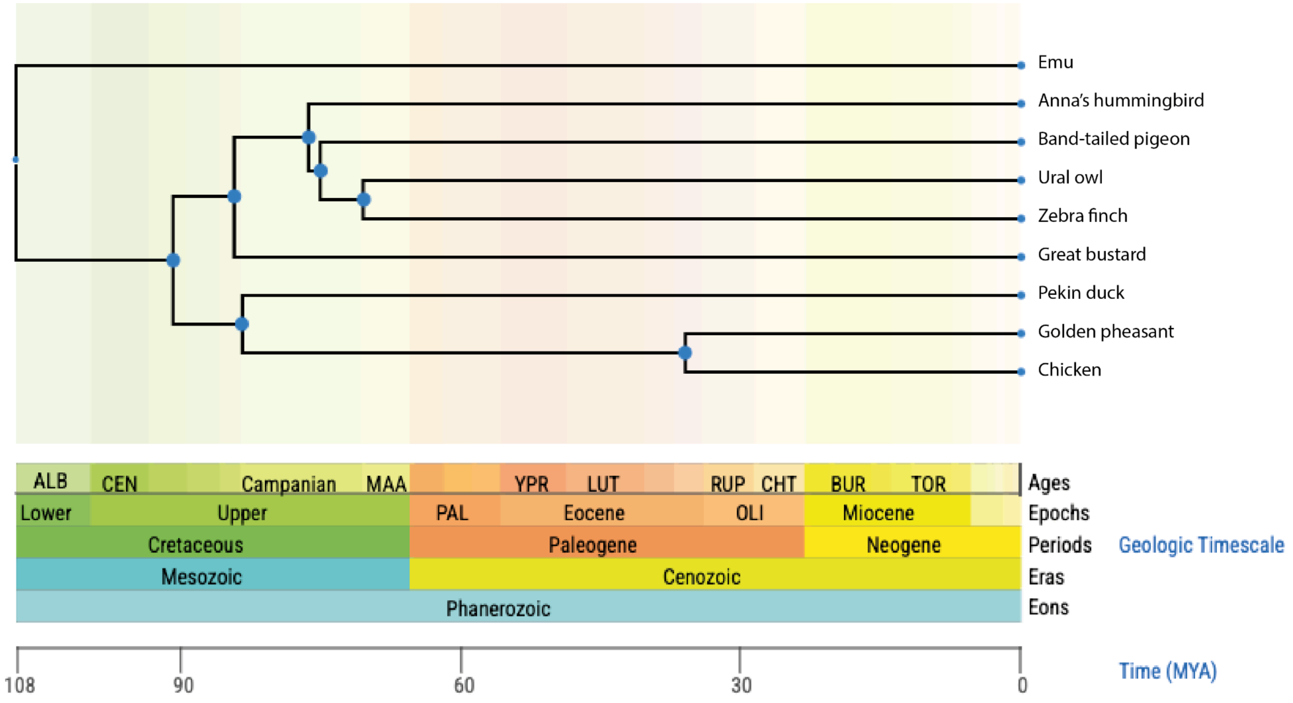

**Figure S5.** Non-B DNA motif coverage vs. chromosome size

Correlation between non-B DNA motif coverage and chromosome size (log-scaled) in nine bird species: *(A)* zebra finch, *(B)* chicken, *(C)* Ural owl, *(D)* band-tailed pigeon, *(E)* Anna's hummingbird, *(F)* great bustard, *(G)* golden pheasant, *(H)* Pekin duck, and *(I)* emu. Note that only the zebra finch assembly is completely T2T, and some of the other species are missing sequences, especially from the dot chromosomes.

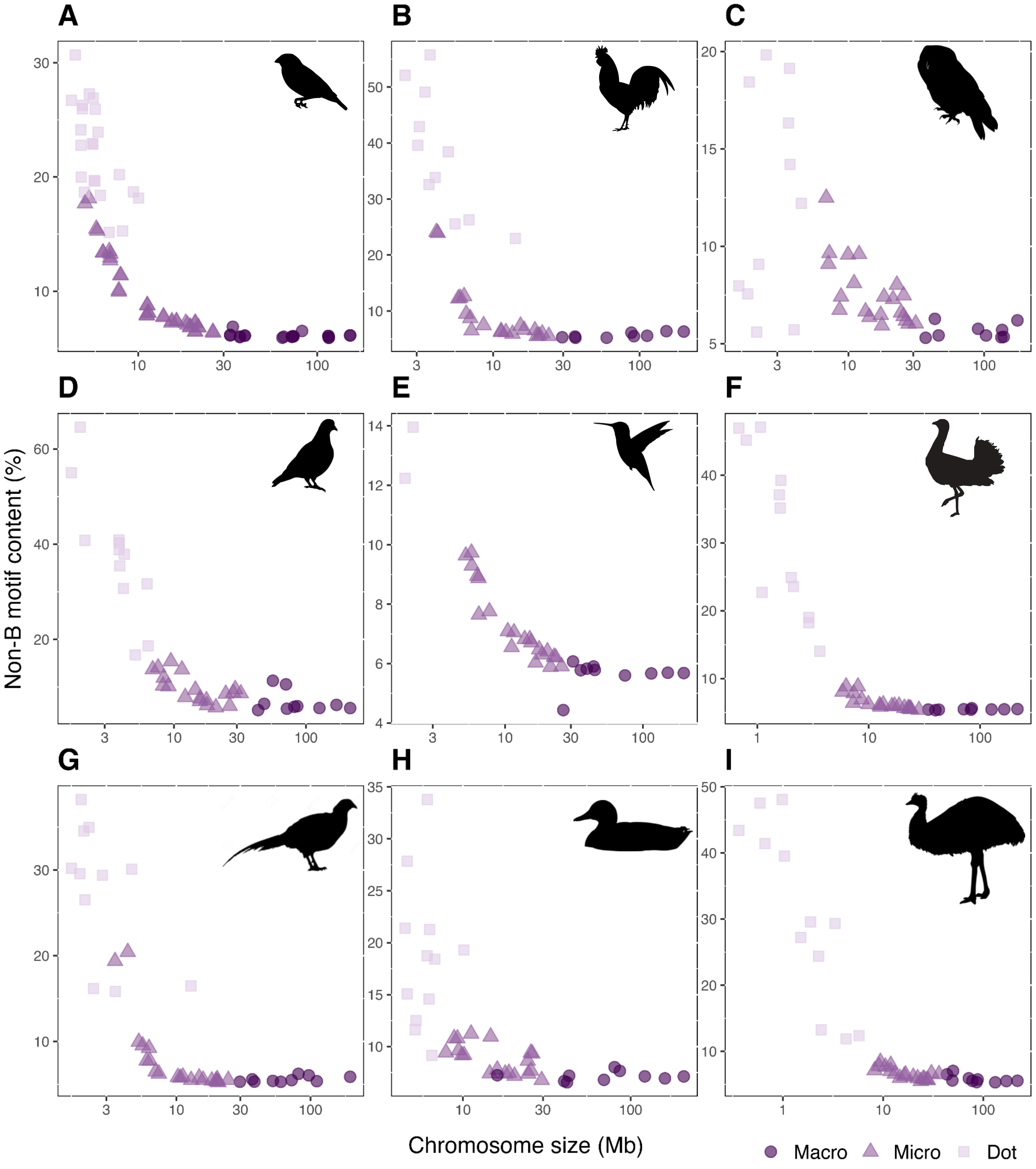

**Figure S6.** Non-B DNA motif coverage vs GC content in bird species
Correlation between non-B DNA motif coverage and GC content in nine bird species: *(A)* zebra finch, *(B)* chicken, *(C)* Ural owl, *(D)* band-tailed pigeon, *(E)* Anna's hummingbird, *(F)* great bustard, *(G)* golden pheasant, *(H)* Pekin duck, and *(I)* emu. Note that only the zebra finch assembly is completely T2T, and missing sequences could influence both non-B DNA motif coverage and GC content in the other species.

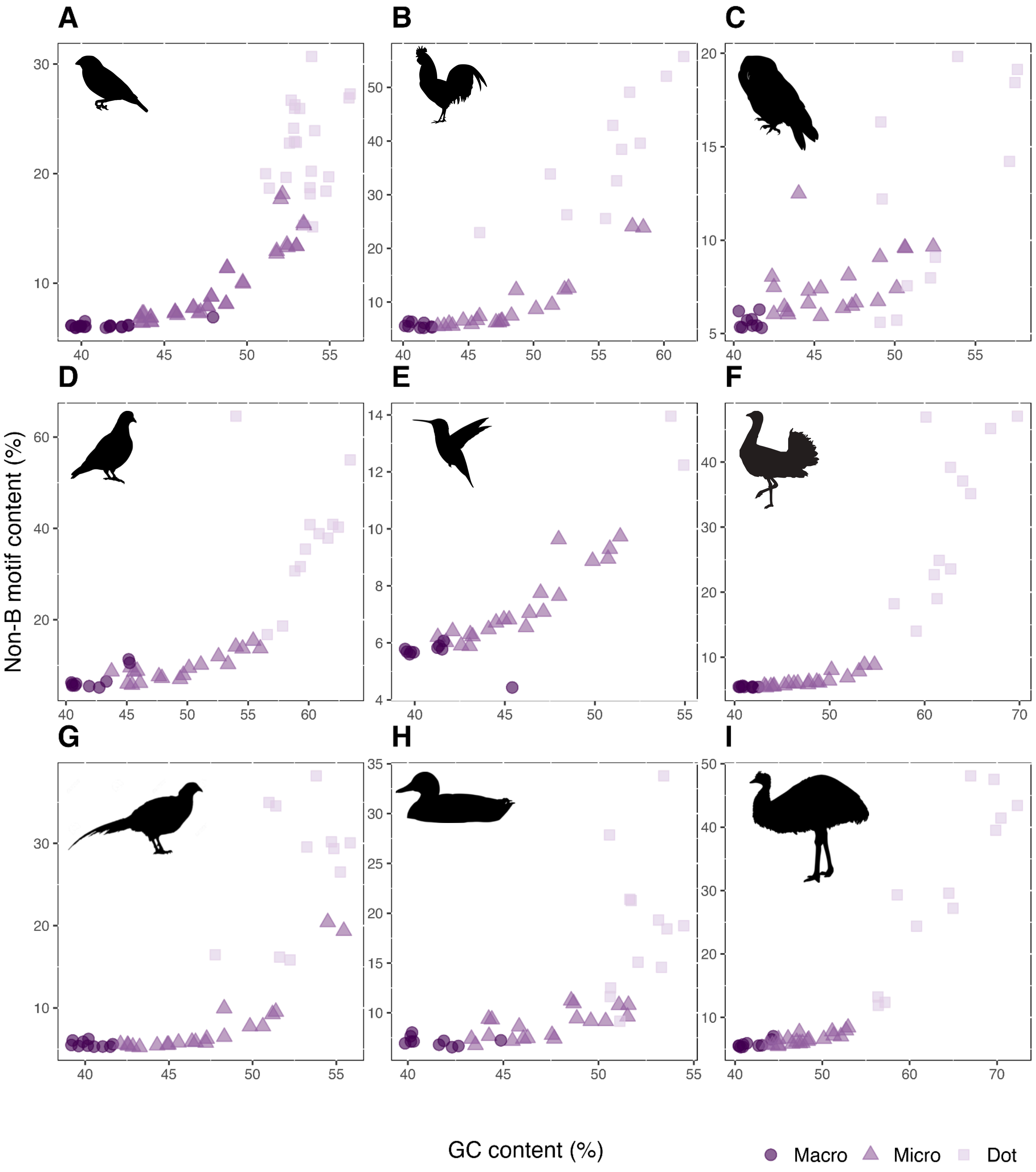

**Figure S7.** Non-B DNA coverage of other bird genomes

Circos plots for *(A)* Ural owl, *(B)* band-tailed pigeon, *(C)* Anna's hummingbird, *(D)* great bustard, *(E)* golden pheasant, *(F)* Pekin duck, *(G)* emu. Abbreviations: APR: A-phased repeats; DR: direct repeats; STR: short tandem repeats; IR: inverted repeats; TRI: triplex motifs; G4: G-quadruplexes; Z: Z-DNA. Bird silhouettes are taken from [https://www.phylopic.org](https://www.phylopic.org) (except bustard, drawn by the authors).

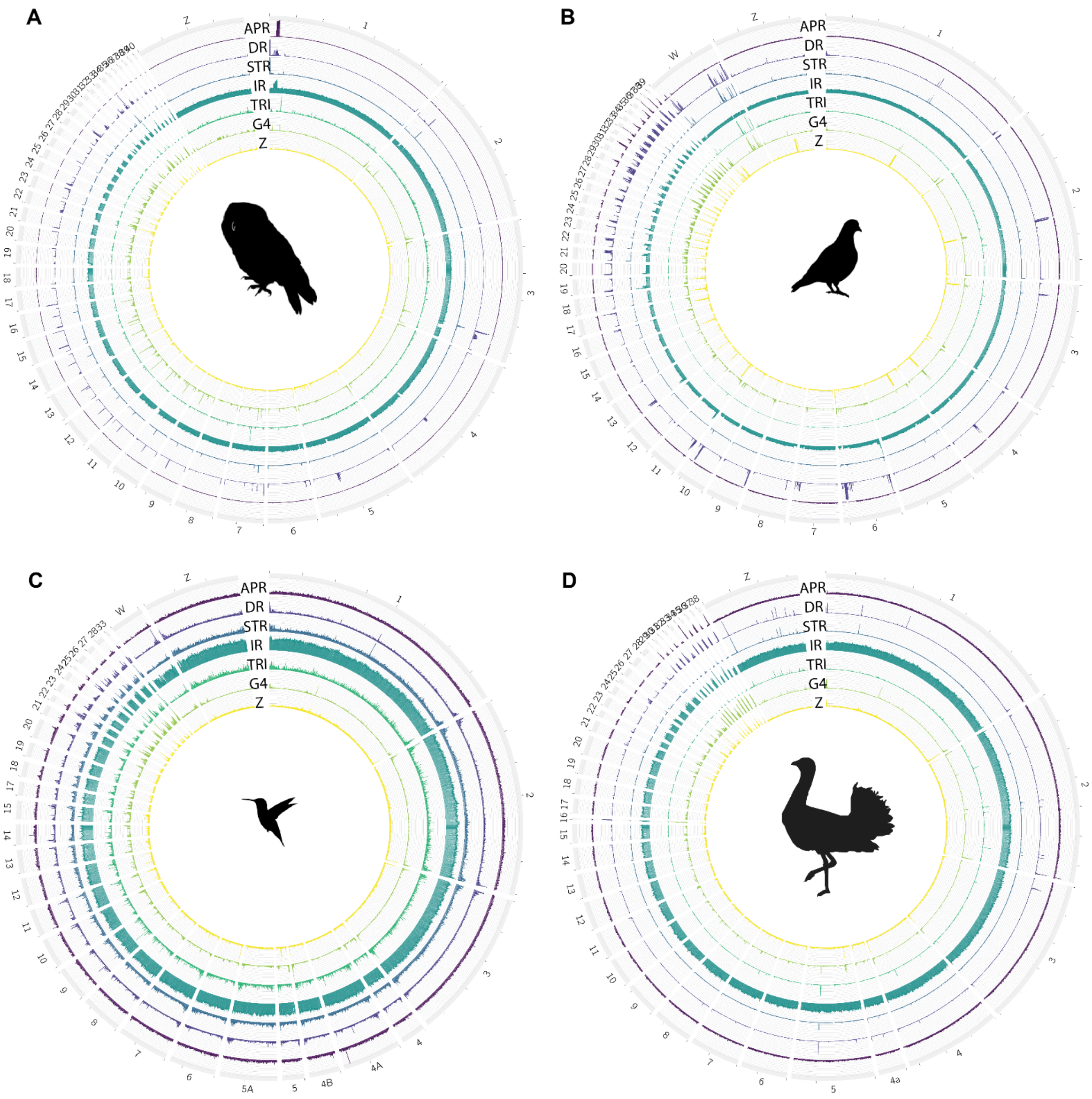

**E**

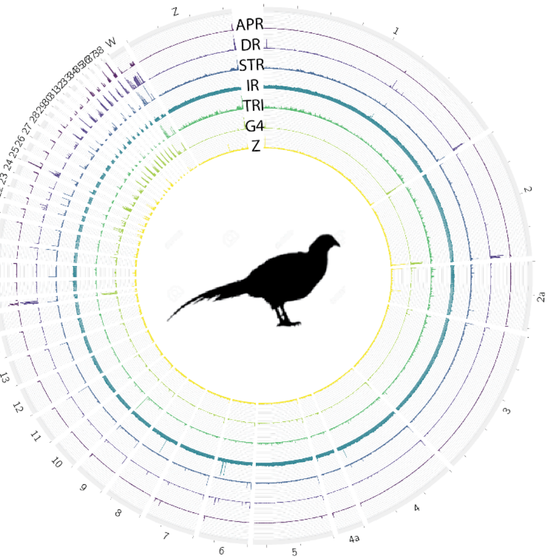
APR
DR
STR
IR
TRI
G4
Z
Z
W
1
2
2a
3
4
4a

**F**

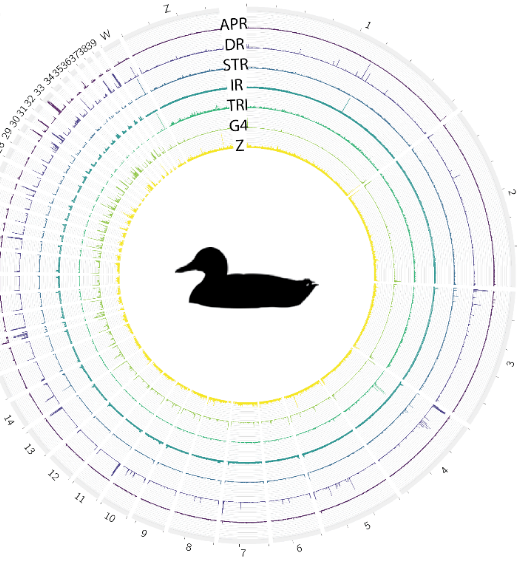
APR
DR
STR
IR
TRI
G4
Z
Z
W

**G**

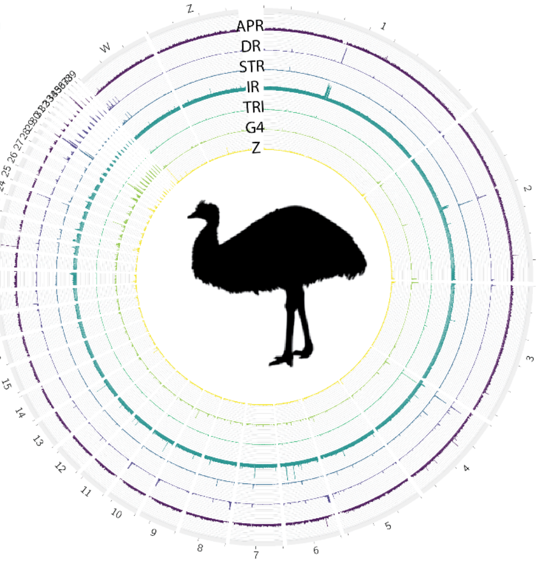
APR
DR
STR
IR
TRI
G4
Z
Z
W

**Figure S8.** Motif overlaps for macro-, micro-, dot chromosomes and sex chromosomes
Overlaps between non-B DNA motif types in different chromosome categories for *(A)* zebra finch (diploid genome), *(B)* chicken, *(C)* Ural owl, *(D)* band-tailed pigeon, *(E)* Anna's hummingbird, *(F)* great bustard, *(G)* Pekin duck, and *(H)* emu. Left side, all panels: pairwise overlap between all non-B motifs, for macro-, micro-, and dot chromosomes, respectively. The number in each box denotes the percent of the motif to the left that overlaps with the motif at the bottom. Right side, all panels: Upset plot with the Mb overlap for all non-B DNA motif combinations (only showing combinations with more than 10 kb overlap for visual purposes); bars are colored based on the number of overlapping motifs. Abbreviations as in Figure S1. Note that Ural owl and great bustard assemblies lack the W chromosome. Bird silhouettes are from phylopic.org.

A

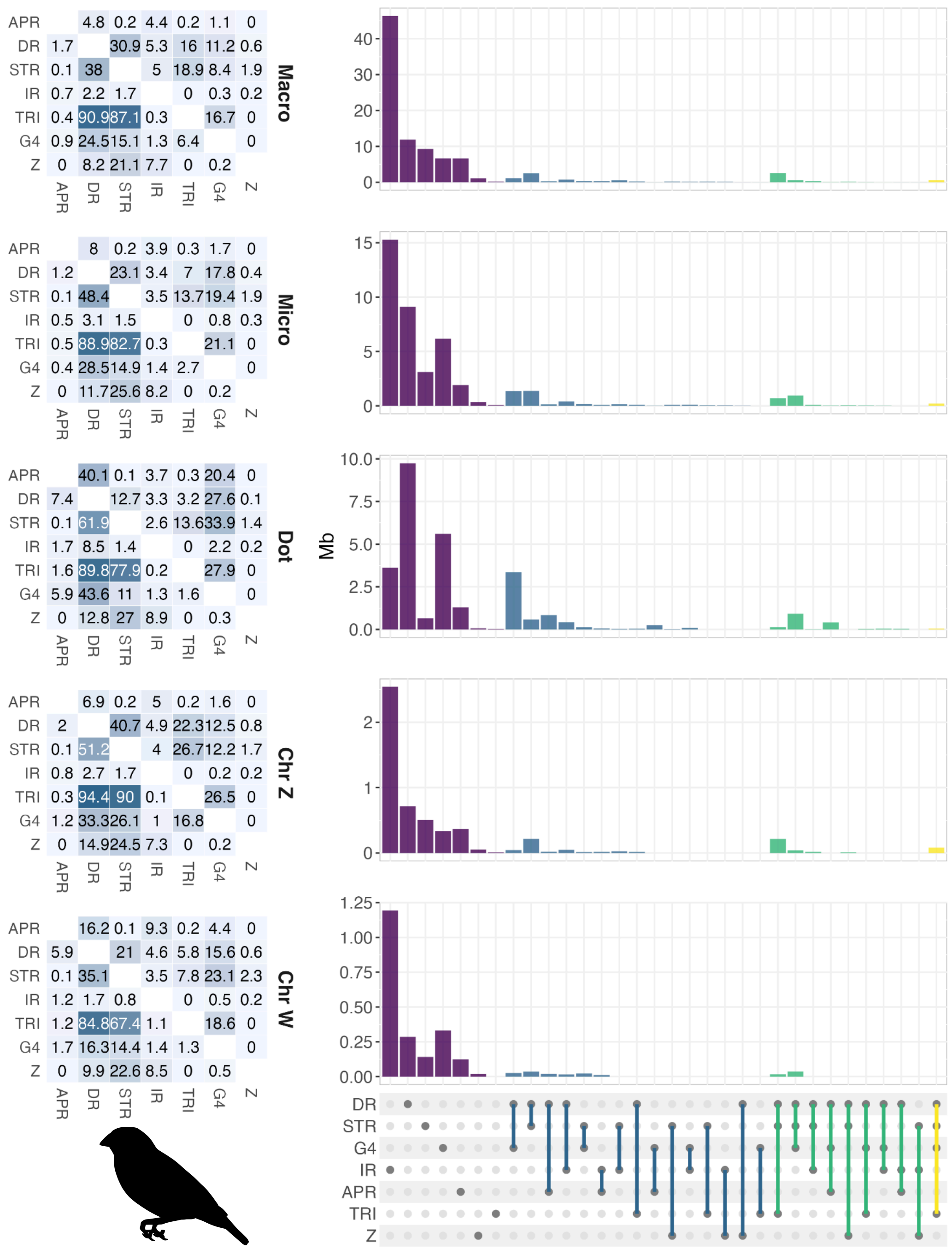

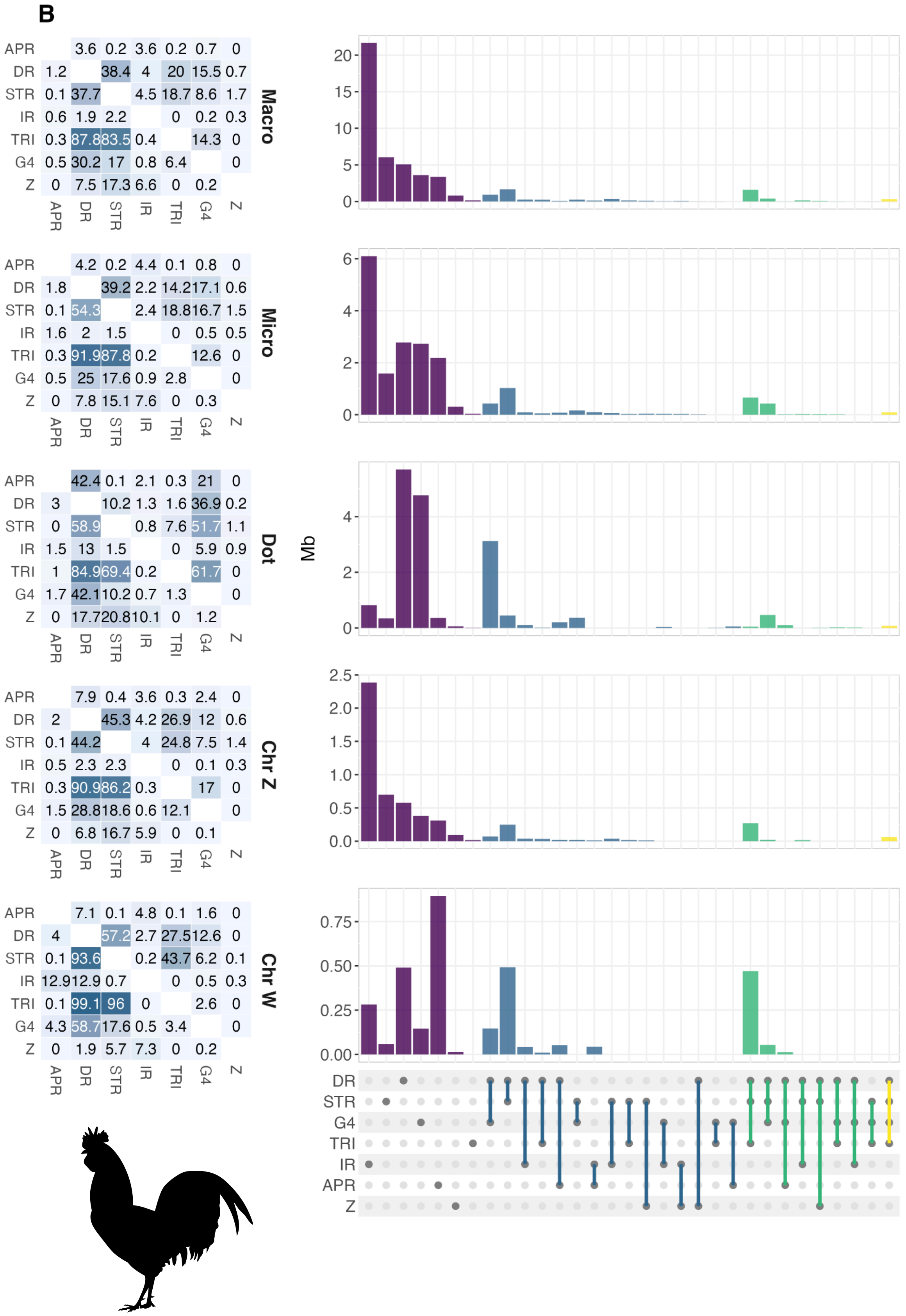
B
APR
DR
STR
IR
TRI
G4
Z
Macro
Micro
Dot
Chr Z
Chr W
Mb

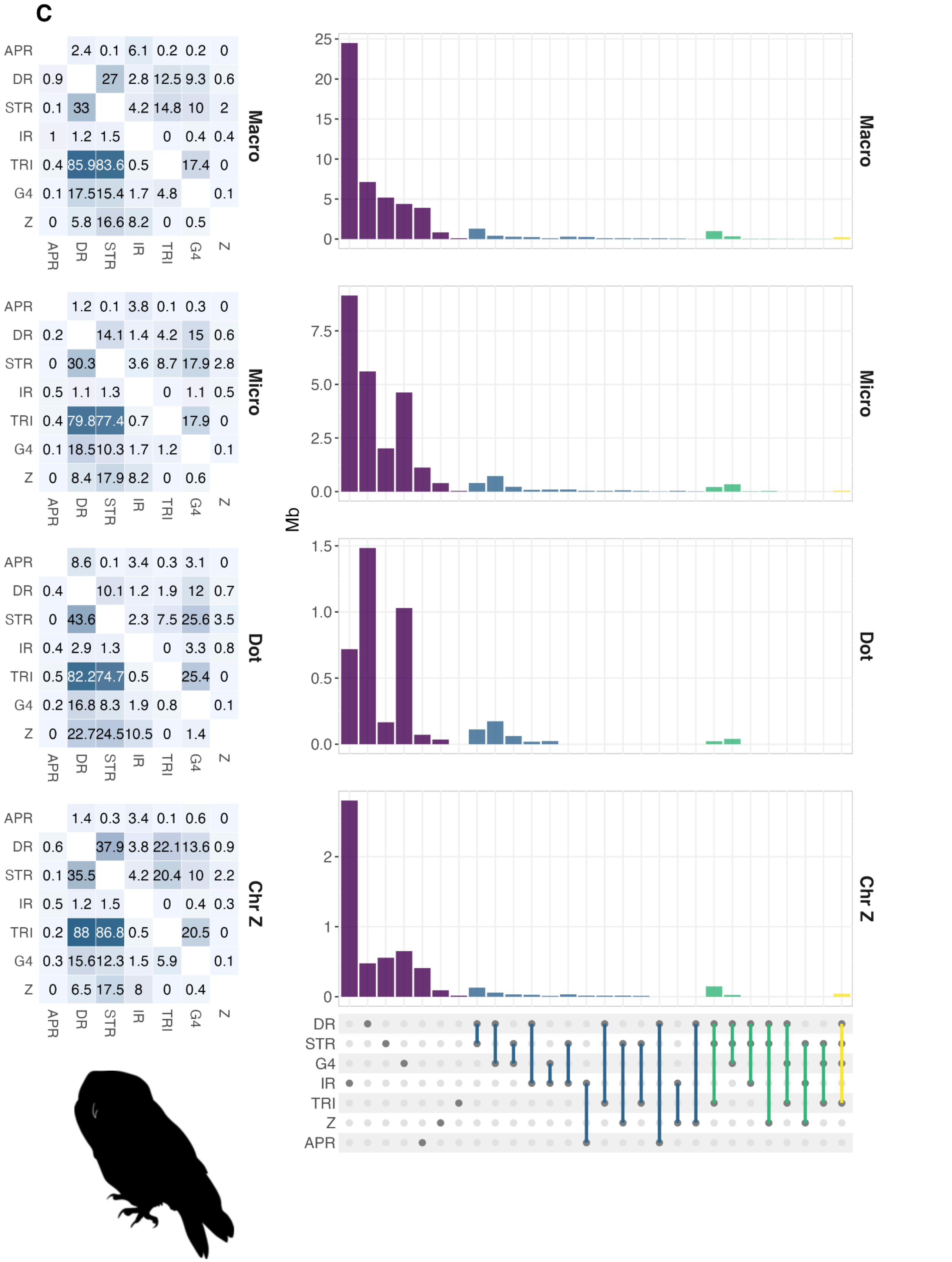
C
APR
DR
STR
IR
TRI
G4
Z
Macro
Micro
Dot
Chr Z
Mb
2.4 0.1 6.1 0.2 0.2 0
0.9 27 2.8 12.5 9.3 0.6
0.1 33 4.2 14.8 10 2
1 1.2 1.5 0 0.4 0.4
0.4 85.9 83.6 0.5 17.4 0
0.1 17.5 15.4 1.7 4.8 0.1
0 5.8 16.6 8.2 0 0.5
1.2 0.1 3.8 0.1 0.3 0
0.2 14.1 1.4 4.2 15 0.6
0 30.3 3.6 8.7 17.9 2.8
0.5 1.1 1.3 0 1.1 0.5
0.4 79.8 77.4 0.7 17.9 0
0.1 18.5 10.3 1.7 1.2 0.1
0 8.4 17.9 8.2 0 0.6
8.6 0.1 3.4 0.3 3.1 0
0.4 10.1 1.2 1.9 12 0.7
0 43.6 2.3 7.5 25.6 3.5
0.4 2.9 1.3 0 3.3 0.8
0.5 82.2 74.7 0.5 25.4 0
0.2 16.8 8.3 1.9 0.8 0.1
0 22.7 24.5 10.5 0 1.4
1.4 0.3 3.4 0.1 0.6 0
0.6 37.9 3.8 22.1 13.6 0.9
0.1 35.5 4.2 20.4 10 2.2
0.5 1.2 1.5 0 0.4 0.3
0.2 88 86.8 0.5 20.5 0
0.3 15.6 12.3 1.5 5.9 0.1
0 6.5 17.5 8 0 0.4

**D**

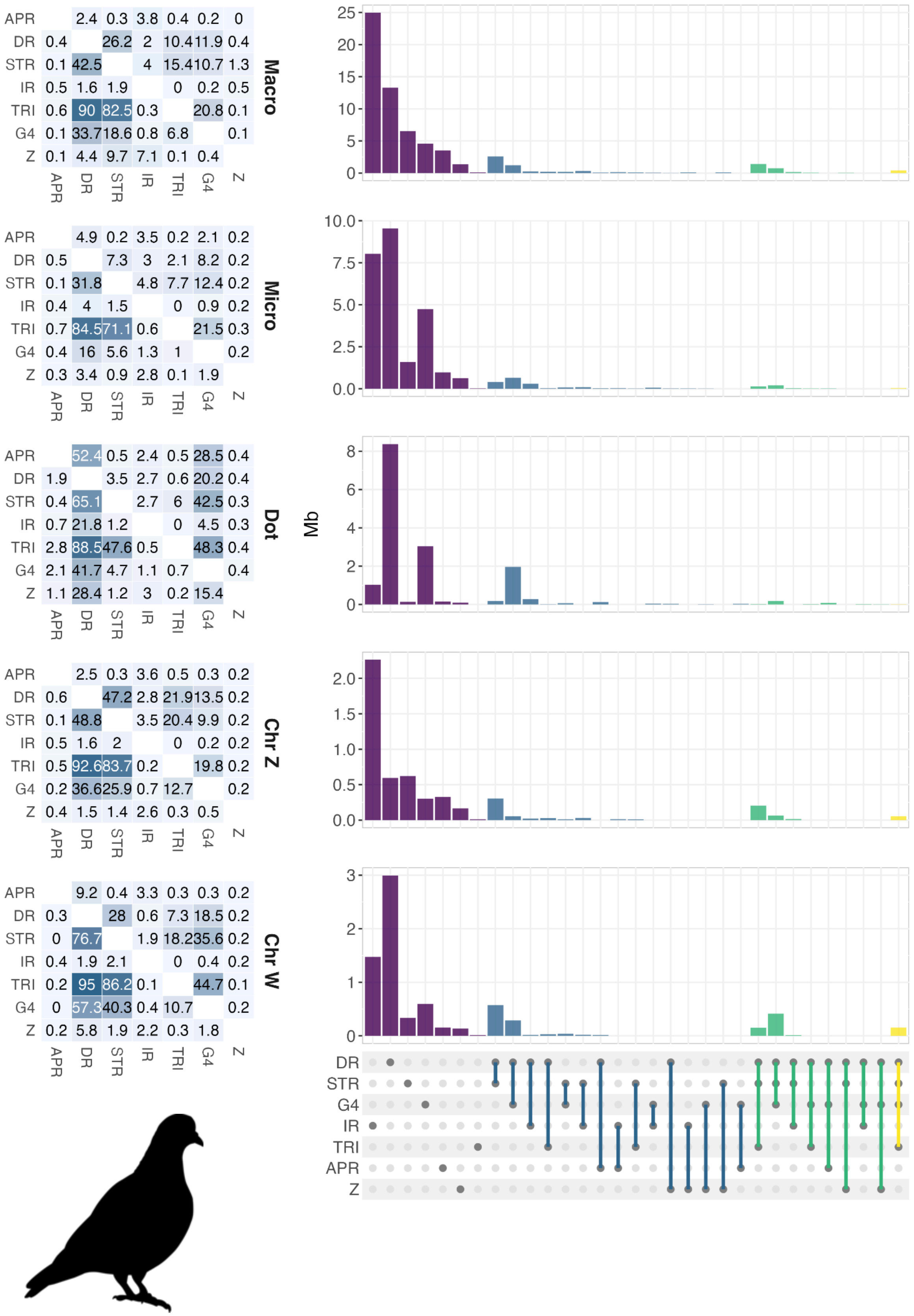

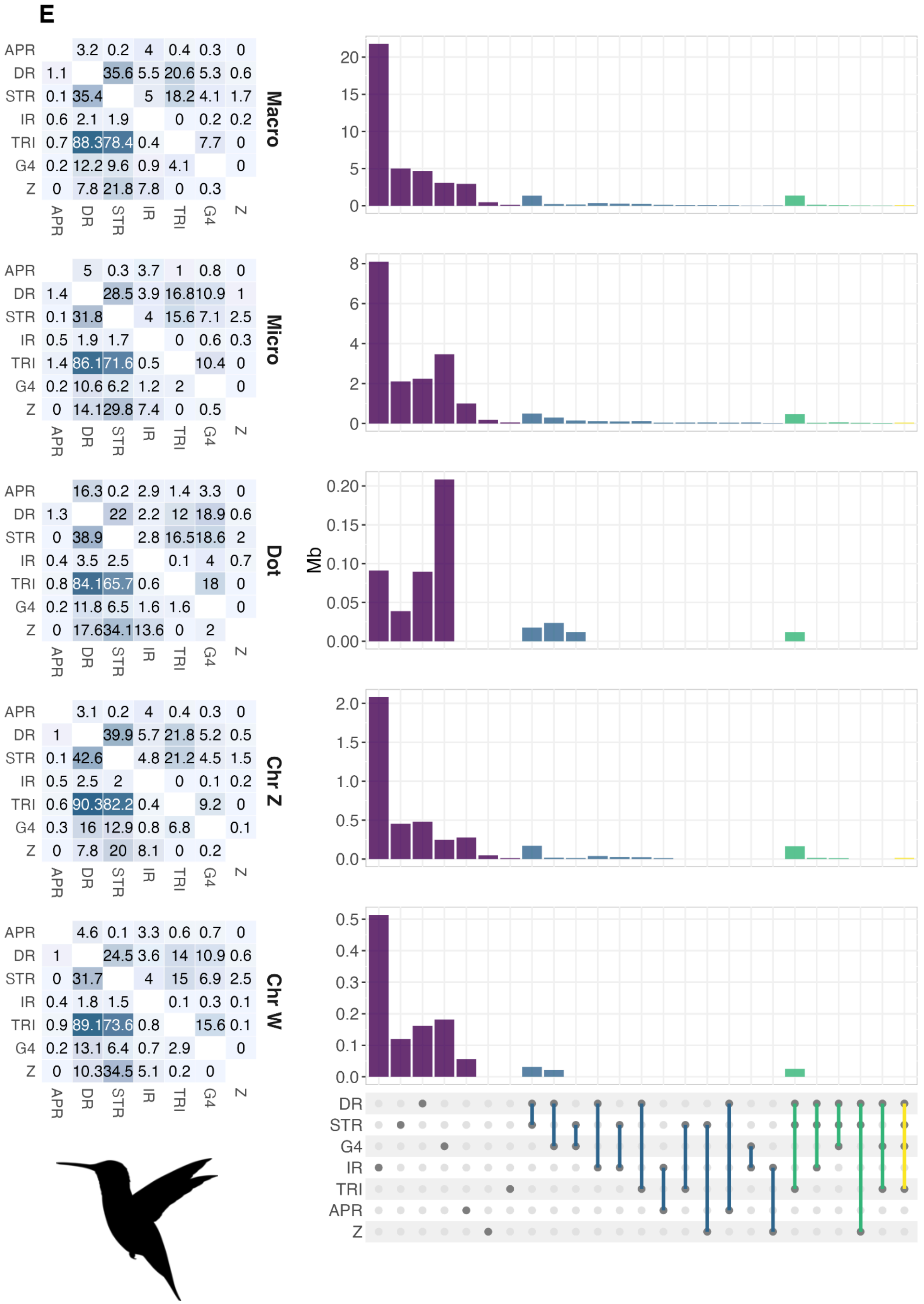
E
Macro
APR 3.2 0.2 4 0.4 0.3 0
DR 1.1 35.6 5.5 20.6 5.3 0.6
STR 0.1 35.4 5 18.2 4.1 1.7
IR 0.6 2.1 1.9 0 0.2 0.2
TRI 0.7 88.3 78.4 0.4 7.7 0
G4 0.2 12.2 9.6 0.9 4.1 0
Z 0 7.8 21.8 7.8 0 0.3
APR DR STR IR TRI G4 Z
Micro
APR 5 0.3 3.7 1 0.8 0
DR 1.4 28.5 3.9 16.8 10.9 1
STR 0.1 31.8 4 15.6 7.1 2.5
IR 0.5 1.9 1.7 0 0.6 0.3
TRI 1.4 86.1 71.6 0.5 10.4 0
G4 0.2 10.6 6.2 1.2 2 0
Z 0 14.1 29.8 7.4 0 0.5
APR DR STR IR TRI G4 Z
Dot
APR 16.3 0.2 2.9 1.4 3.3 0
DR 1.3 22 2.2 12 18.9 0.6
STR 0 38.9 2.8 16.5 18.6 2
IR 0.4 3.5 2.5 0.1 4 0.7
TRI 0.8 84.1 65.7 0.6 18 0
G4 0.2 11.8 6.5 1.6 1.6 0
Z 0 17.6 34.1 13.6 0 2
APR DR STR IR TRI G4 Z
Chr Z
APR 3.1 0.2 4 0.4 0.3 0
DR 1 39.9 5.7 21.8 5.2 0.5
STR 0.1 42.6 4.8 21.2 4.5 1.5
IR 0.5 2.5 2 0 0.1 0.2
TRI 0.6 90.3 82.2 0.4 9.2 0
G4 0.3 16 12.9 0.8 6.8 0.1
Z 0 7.8 20 8.1 0 0.2
APR DR STR IR TRI G4 Z
Chr W
APR 4.6 0.1 3.3 0.6 0.7 0
DR 1 24.5 3.6 14 10.9 0.6
STR 0 31.7 4 15 6.9 2.5
IR 0.4 1.8 1.5 0.1 0.3 0.1
TRI 0.9 89.1 73.6 0.8 15.6 0.1
G4 0.2 13.1 6.4 0.7 2.9 0
Z 0 10.3 34.5 5.1 0.2 0
APR DR STR IR TRI G4 Z
Mb
DR STR G4 IR TRI APR Z

F

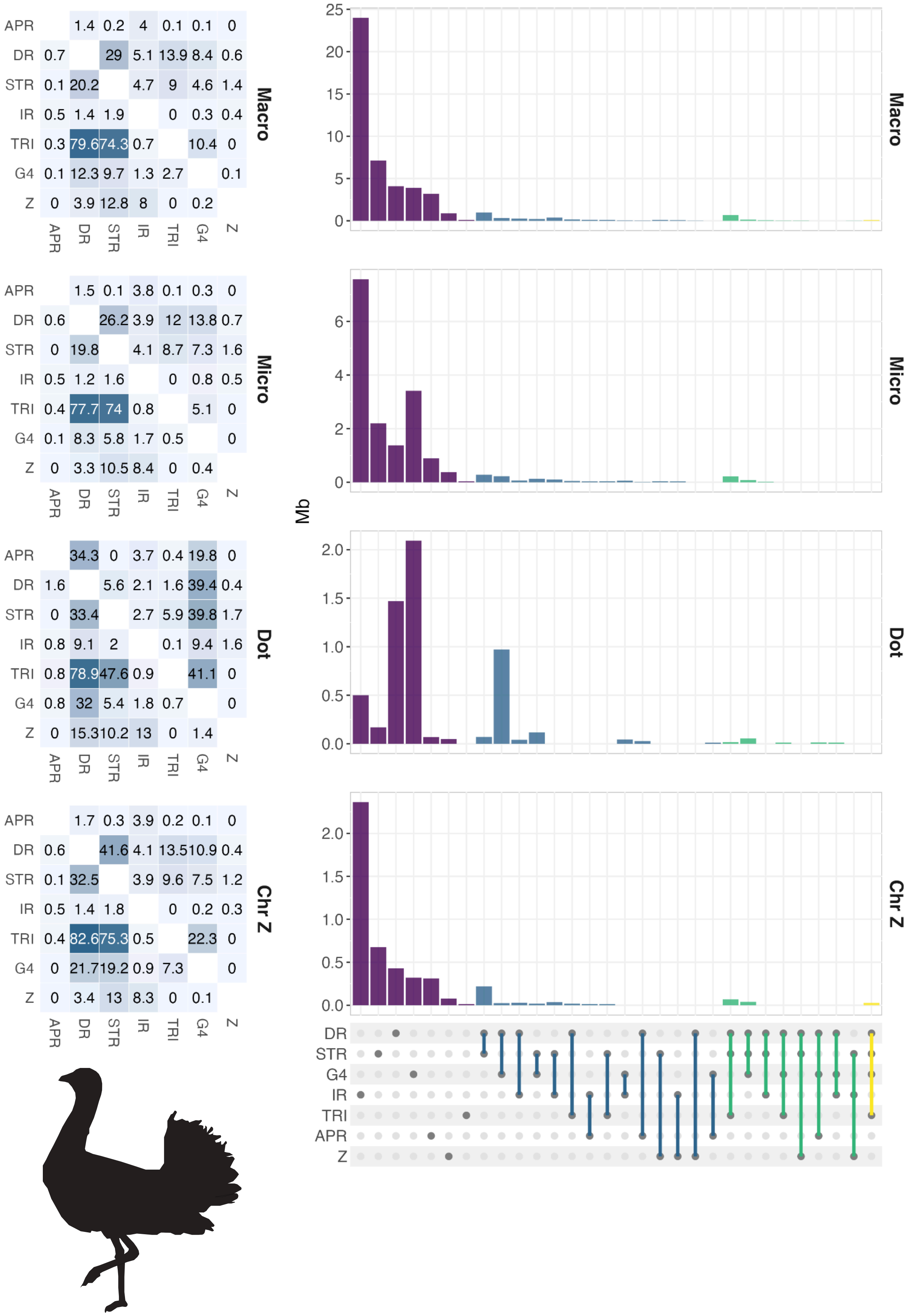
Macro
Micro
Dot
Chr Z
APR
DR
STR
IR
TRI
G4
Z
Mb
25
20
15
10
5
0
6
4
2
2.0
1.5
1.0
0.5
0.0

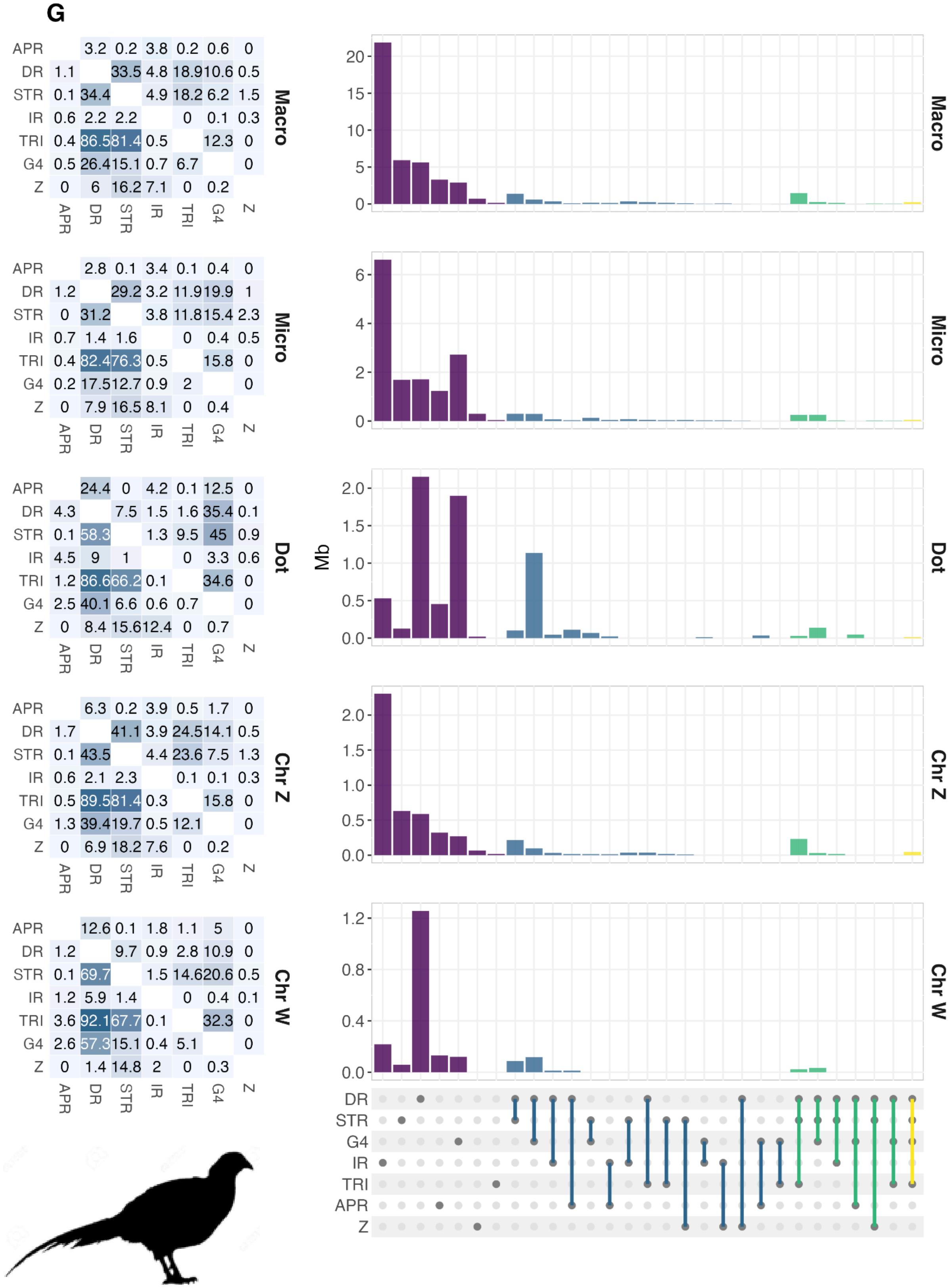
G
APR 3.2 0.2 3.8 0.2 0.6 0
DR 1.1 33.5 4.8 18.9 10.6 0.5
STR 0.1 34.4 4.9 18.2 6.2 1.5
IR 0.6 2.2 2.2 0 0.1 0.3
TRI 0.4 86.5 81.4 0.5 12.3 0
G4 0.5 26.4 15.1 0.7 6.7 0
Z 0 6 16.2 7.1 0 0.2
APR DR STR IR TRI G4 Z
Macro
APR 2.8 0.1 3.4 0.1 0.4 0
DR 1.2 29.2 3.2 11.9 19.9 1
STR 0 31.2 3.8 11.8 15.4 2.3
IR 0.7 1.4 1.6 0 0.4 0.5
TRI 0.4 82.4 76.3 0.5 15.8 0
G4 0.2 17.5 12.7 0.9 2 0
Z 0 7.9 16.5 8.1 0 0.4
APR DR STR IR TRI G4 Z
Micro
APR 24.4 0 4.2 0.1 12.5 0
DR 4.3 7.5 1.5 1.6 35.4 0.1
STR 0.1 58.3 1.3 9.5 45 0.9
IR 4.5 9 1 0 3.3 0.6
TRI 1.2 86.6 66.2 0.1 34.6 0
G4 2.5 40.1 6.6 0.6 0.7 0
Z 0 8.4 15.6 12.4 0 0.7
APR DR STR IR TRI G4 Z
Dot
APR 6.3 0.2 3.9 0.5 1.7 0
DR 1.7 41.1 3.9 24.5 14.1 0.5
STR 0.1 43.5 4.4 23.6 7.5 1.3
IR 0.6 2.1 2.3 0.1 0.1 0.3
TRI 0.5 89.5 81.4 0.3 15.8 0
G4 1.3 39.4 19.7 0.5 12.1 0
Z 0 6.9 18.2 7.6 0 0.2
APR DR STR IR TRI G4 Z
Chr Z
APR 12.6 0.1 1.8 1.1 5 0
DR 1.2 9.7 0.9 2.8 10.9 0
STR 0.1 69.7 1.5 14.6 20.6 0.5
IR 1.2 5.9 1.4 0 0.4 0.1
TRI 3.6 92.1 67.7 0.1 32.3 0
G4 2.6 57.3 15.1 0.4 5.1 0
Z 0 1.4 14.8 2 0 0.3
APR DR STR IR TRI G4 Z
Chr W
Mb
Macro
Micro
Dot
Chr Z
Chr W
DR
STR
G4
IR
TRI
APR
Z

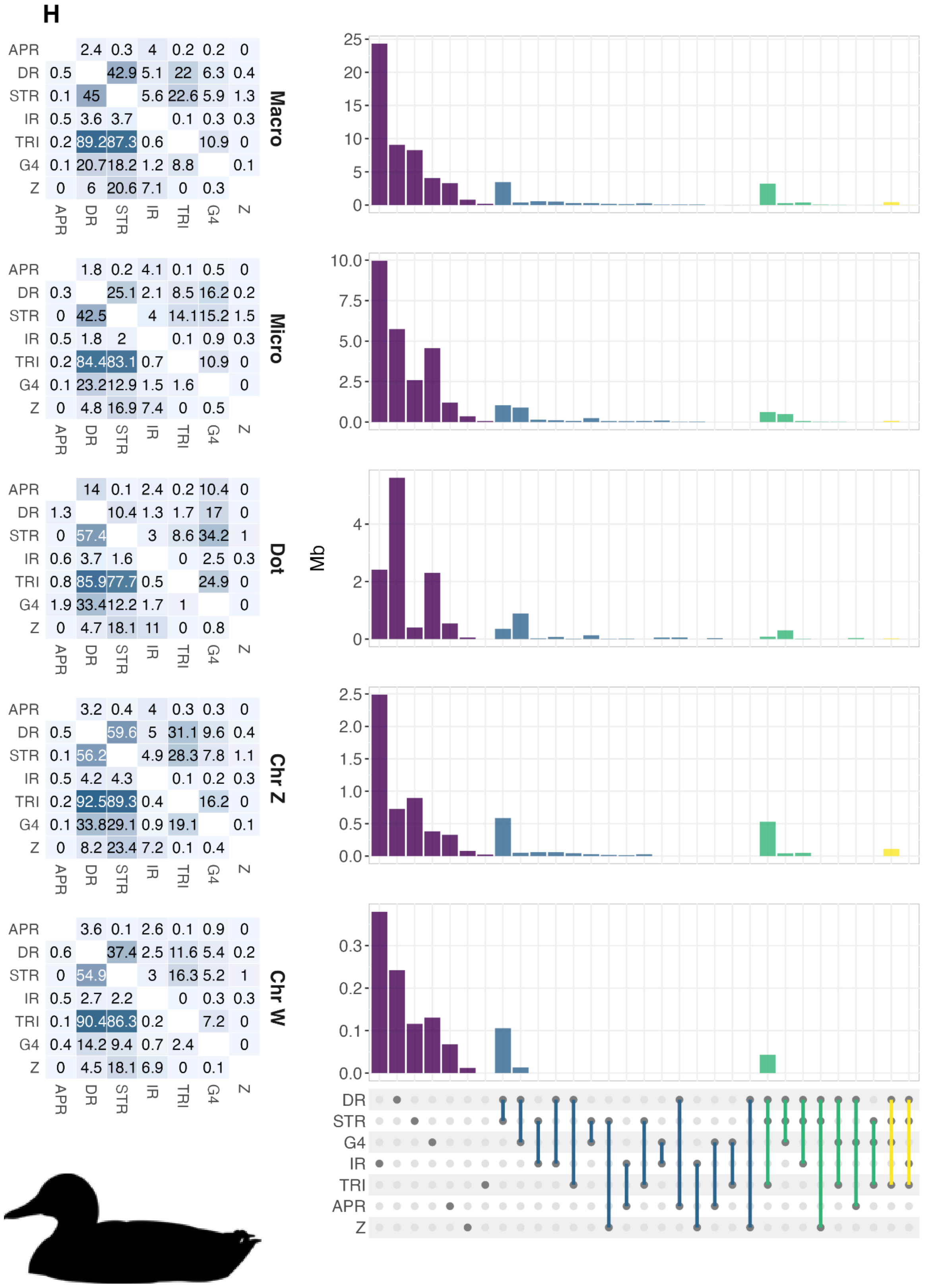
H
APR 2.4 0.3 4 0.2 0.2 0
DR 0.5 42.9 5.1 22 6.3 0.4
STR 0.1 45 5.6 22.6 5.9 1.3
IR 0.5 3.6 3.7 0.1 0.3 0.3
TRI 0.2 89.2 87.3 0.6 10.9 0
G4 0.1 20.7 18.2 1.2 8.8 0.1
Z 0 6 20.6 7.1 0 0.3
APR DR STR IR TRI G4 Z
Macro
APR 1.8 0.2 4.1 0.1 0.5 0
DR 0.3 25.1 2.1 8.5 16.2 0.2
STR 0 42.5 4 14.1 15.2 1.5
IR 0.5 1.8 2 0.1 0.9 0.3
TRI 0.2 84.4 83.1 0.7 10.9 0
G4 0.1 23.2 12.9 1.5 1.6 0
Z 0 4.8 16.9 7.4 0 0.5
APR DR STR IR TRI G4 Z
Micro
APR 14 0.1 2.4 0.2 10.4 0
DR 1.3 10.4 1.3 1.7 17 0
STR 0 57.4 3 8.6 34.2 1
IR 0.6 3.7 1.6 0 2.5 0.3
TRI 0.8 85.9 77.7 0.5 24.9 0
G4 1.9 33.4 12.2 1.7 1 0
Z 0 4.7 18.1 11 0 0.8
APR DR STR IR TRI G4 Z
Dot
APR 3.2 0.4 4 0.3 0.3 0
DR 0.5 59.6 5 31.1 9.6 0.4
STR 0.1 56.2 4.9 28.3 7.8 1.1
IR 0.5 4.2 4.3 0.1 0.2 0.3
TRI 0.2 92.5 89.3 0.4 16.2 0
G4 0.1 33.8 29.1 0.9 19.1 0.1
Z 0 8.2 23.4 7.2 0.1 0.4
APR DR STR IR TRI G4 Z
Chr Z
APR 3.6 0.1 2.6 0.1 0.9 0
DR 0.6 37.4 2.5 11.6 5.4 0.2
STR 0 54.9 3 16.3 5.2 1
IR 0.5 2.7 2.2 0 0.3 0.3
TRI 0.1 90.4 86.3 0.2 7.2 0
G4 0.4 14.2 9.4 0.7 2.4 0
Z 0 4.5 18.1 6.9 0 0.1
APR DR STR IR TRI G4 Z
Chr W
Mb
25 20 15 10 5 0
Macro
10.0 7.5 5.0 2.5 0.0
Micro
4 2 0
Dot
2.5 2.0 1.5 1.0 0.5 0.0
Chr Z
0.3 0.2 0.1 0.0
Chr W
DR
STR
G4
IR
TRI
APR
Z

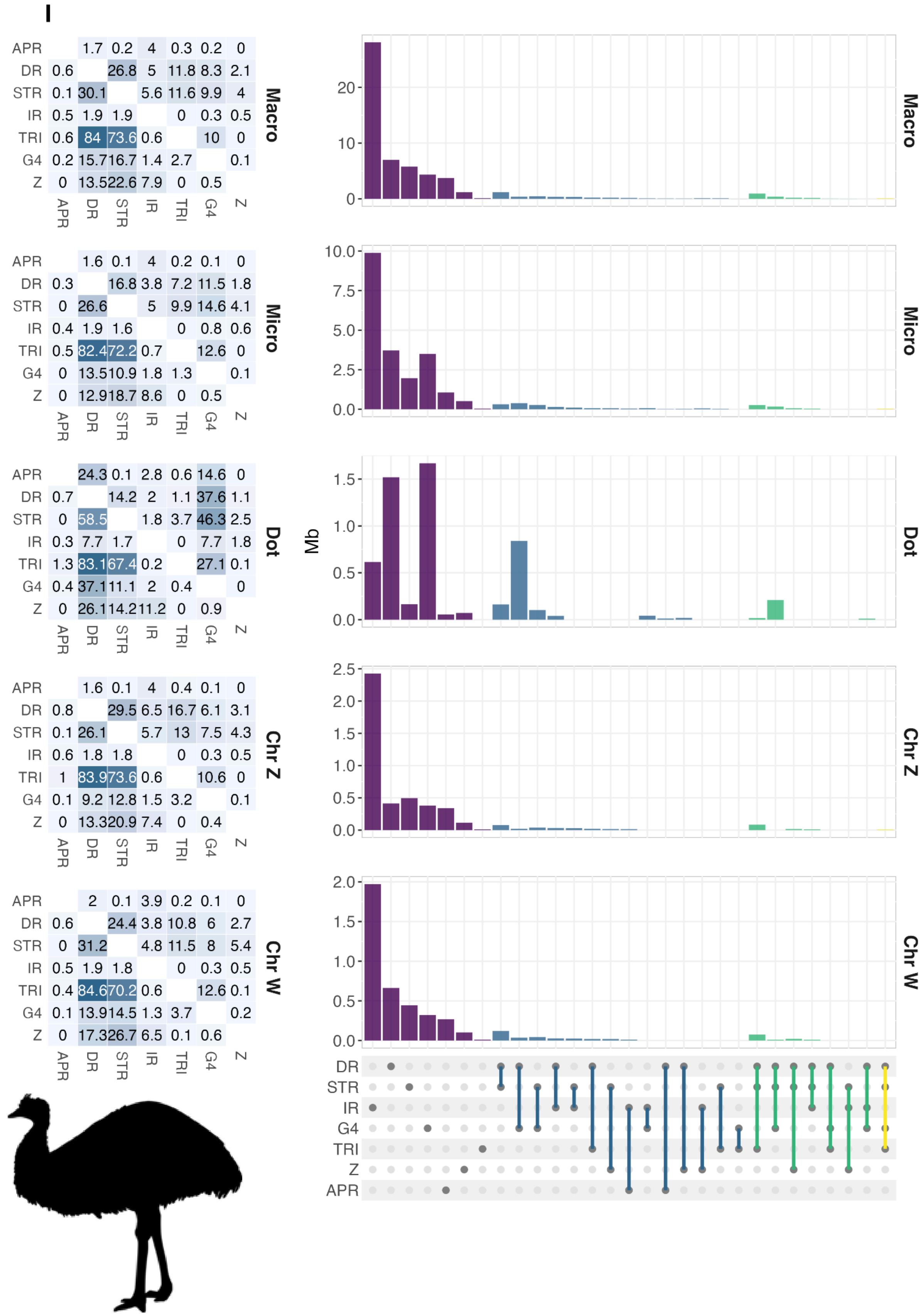
I
APR 1.7 0.2 4 0.3 0.2 0
DR 0.6 26.8 5 11.8 8.3 2.1
STR 0.1 30.1 5.6 11.6 9.9 4
IR 0.5 1.9 1.9 0 0.3 0.5
TRI 0.6 84 73.6 0.6 10 0
G4 0.2 15.7 16.7 1.4 2.7 0.1
Z 0 13.5 22.6 7.9 0 0.5
APR DR STR IR TRI G4 Z
Macro
APR 1.6 0.1 4 0.2 0.1 0
DR 0.3 16.8 3.8 7.2 11.5 1.8
STR 0 26.6 5 9.9 14.6 4.1
IR 0.4 1.9 1.6 0 0.8 0.6
TRI 0.5 82.4 72.2 0.7 12.6 0
G4 0 13.5 10.9 1.8 1.3 0.1
Z 0 12.9 18.7 8.6 0 0.5
APR DR STR IR TRI G4 Z
Micro
APR 24.3 0.1 2.8 0.6 14.6 0
DR 0.7 14.2 2 1.1 37.6 1.1
STR 0 58.5 1.8 3.7 46.3 2.5
IR 0.3 7.7 1.7 0 7.7 1.8
TRI 1.3 83.1 67.4 0.2 27.1 0.1
G4 0.4 37.1 11.1 2 0.4 0
Z 0 26.1 14.2 11.2 0 0.9
APR DR STR IR TRI G4 Z
Dot
APR 1.6 0.1 4 0.4 0.1 0
DR 0.8 29.5 6.5 16.7 6.1 3.1
STR 0.1 26.1 5.7 13 7.5 4.3
IR 0.6 1.8 1.8 0 0.3 0.5
TRI 1 83.9 73.6 0.6 10.6 0
G4 0.1 9.2 12.8 1.5 3.2 0.1
Z 0 13.3 20.9 7.4 0 0.4
APR DR STR IR TRI G4 Z
Chr Z
APR 2 0.1 3.9 0.2 0.1 0
DR 0.6 24.4 3.8 10.8 6 2.7
STR 0 31.2 4.8 11.5 8 5.4
IR 0.5 1.9 1.8 0 0.3 0.5
TRI 0.4 84.6 70.2 0.6 12.6 0.1
G4 0.1 13.9 14.5 1.3 3.7 0.2
Z 0 17.3 26.7 6.5 0.1 0.6
APR DR STR IR TRI G4 Z
Chr W
Mb
0
10
20
0.0
2.5
5.0
7.5
10.0
0.5
1.0
1.5
2.0
DR
STR
IR
G4
TRI
Z
APR

**Figure S9.** Non-B DNA motif enrichment in functional regions in bird genomes
*(A)*,*(C)*,*(E)*,*(G)*,*(I)*,*(K)* Non-B DNA enrichment at functional regions on macro-, micro-, and dot chromosomes, as compared to genome-wide non-B DNA motif content. Average non-B DNA motif content per chromosome category are indicated to the right of the bars. For each bar, an interval was constructed by downsampling the data to 50% 100 times, and excluding the two highest and the two lowest enrichment values obtained; a bar is marked with "*" if the entire interval is above the chromosome category baseline. *(B)*,*(D)*,*(F)*,*(H)*,*(J)*,*(L)* Coverage of G4 motifs on the coding and template strand. Bars are compared using a paired Wilcoxon test on the coverage per gene for each group, corrected with FDR, *** $P<0.001$, ** $P<0.01$, * $P<0.05$, and 'ns' denotes non-significant. Note that intergenic regions are strand-ignorant and therefore excluded here. A,B show Ural owl, C,D band-tailed pigeon, E,F Anna's hummingbird, G,H great bustard, I,J Pekin duck, and K,L emu. Note that golden pheasant lacks a RefSeq gene annotation and is therefore excluded.

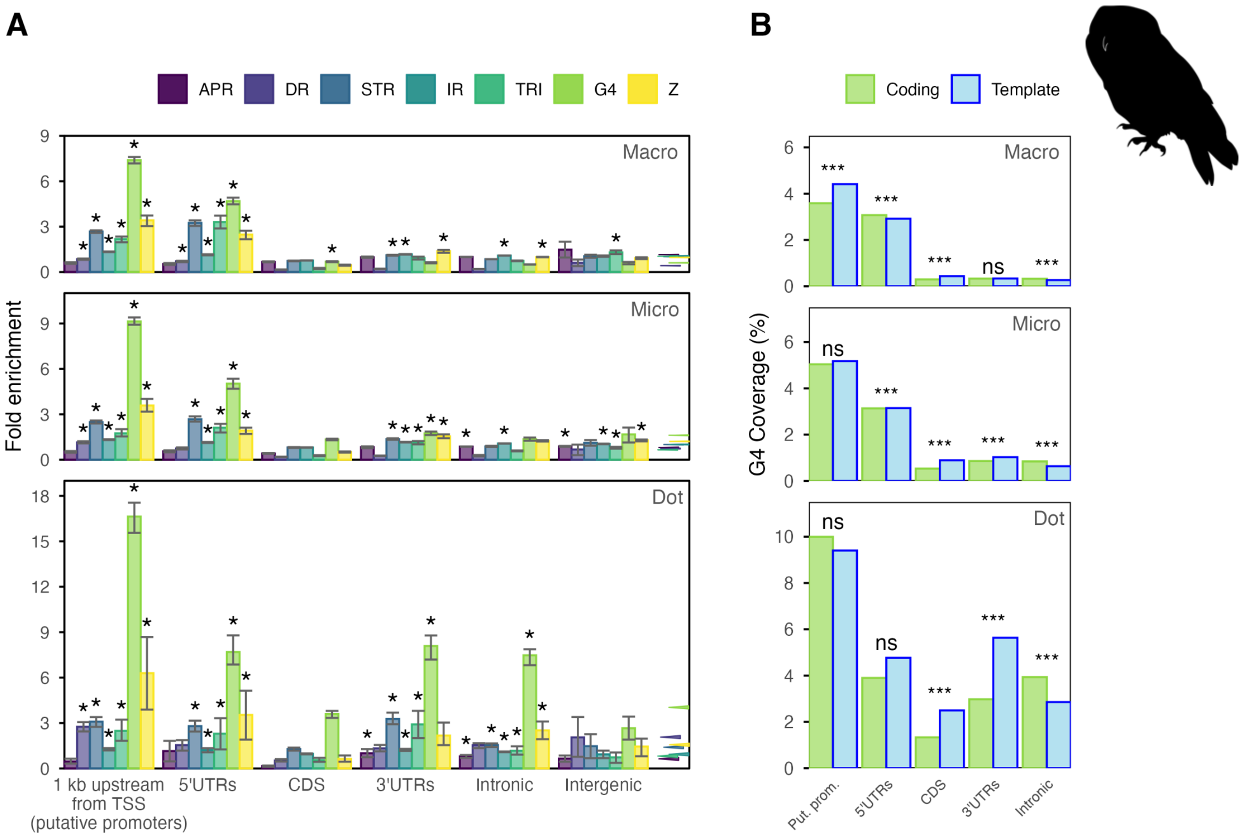

**C**

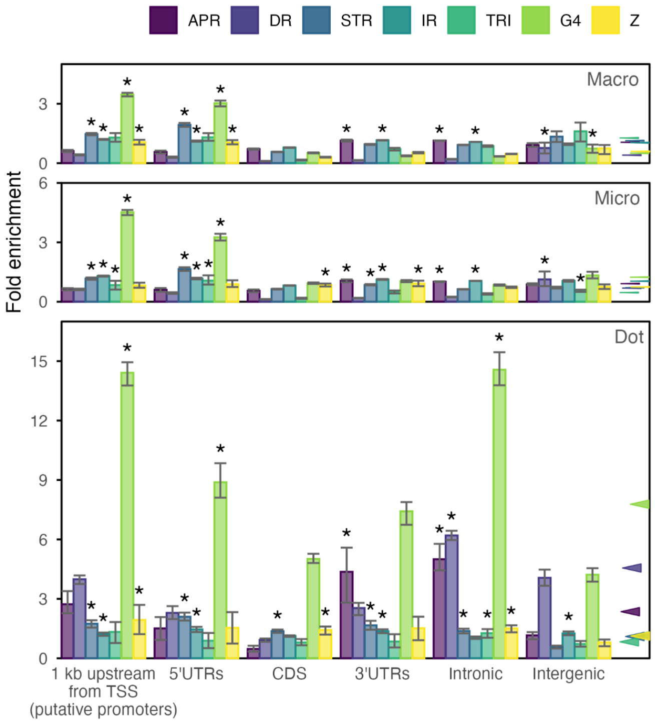
APR
DR
STR
IR
TRI
G4
Z
Macro
Micro
Dot
Fold enrichment
1 kb upstream from TSS (putative promoters)
5'UTRs
CDS
3'UTRs
Intronic
Intergenic

**D**

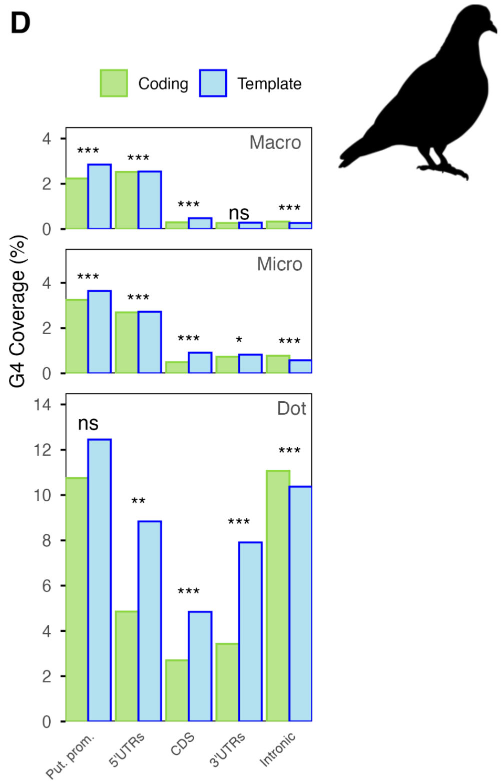
Coding
Template
Macro
Micro
Dot
G4 Coverage (%)
Put. prom.
5'UTRs
CDS
3'UTRs
Intronic

**E**

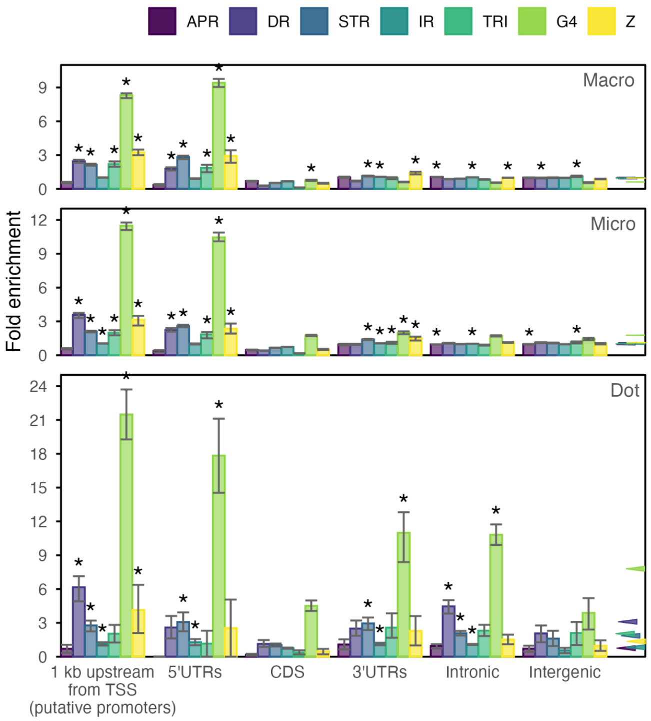
APR
DR
STR
IR
TRI
G4
Z
Macro
Micro
Dot
Fold enrichment
1 kb upstream from TSS (putative promoters)
5'UTRs
CDS
3'UTRs
Intronic
Intergenic

**F**

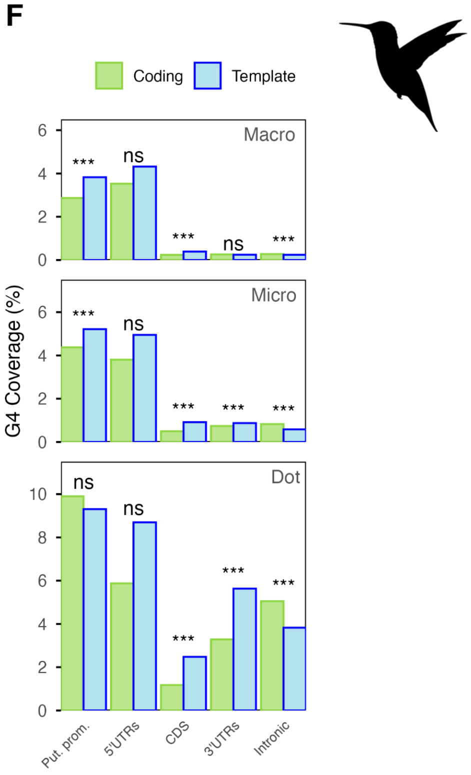
Coding
Template
Macro
Micro
Dot
G4 Coverage (%)
Put. prom.
5'UTRs
CDS
3'UTRs
Intronic

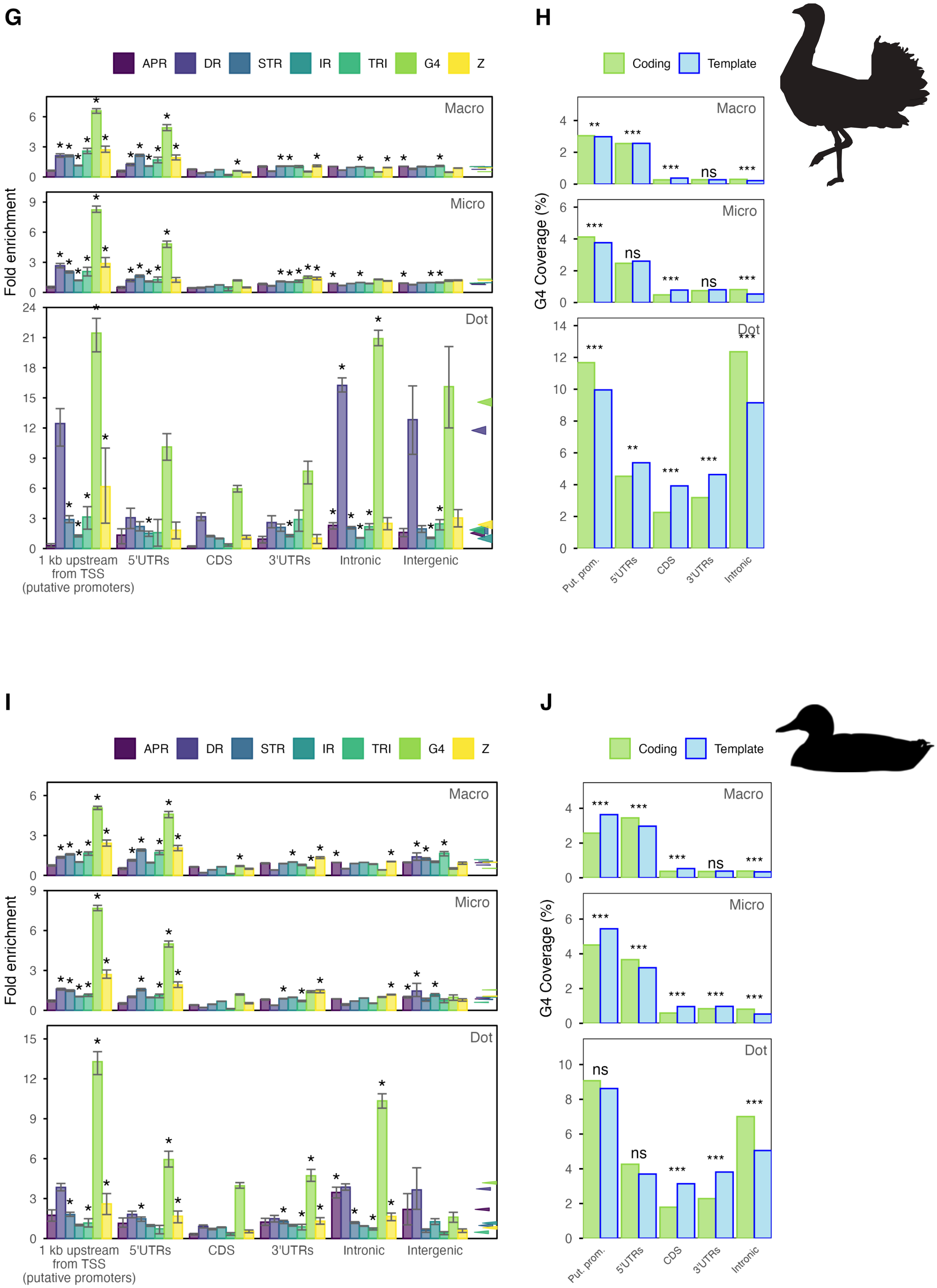
G
APR DR STR IR TRI G4 Z
Fold enrichment
Macro
Micro
Dot
1 kb upstream from TSS (putative promoters)
5'UTRs
CDS
3'UTRs
Intronic
Intergenic
H
Coding Template
G4 Coverage (%)
Macro
Micro
Dot
Put. prom.
5'UTRs
CDS
3'UTRs
Intronic
I
APR DR STR IR TRI G4 Z
Fold enrichment
Macro
Micro
Dot
1 kb upstream from TSS (putative promoters)
5'UTRs
CDS
3'UTRs
Intronic
Intergenic
J
Coding Template
G4 Coverage (%)
Macro
Micro
Dot
Put. prom.
5'UTRs
CDS
3'UTRs
Intronic

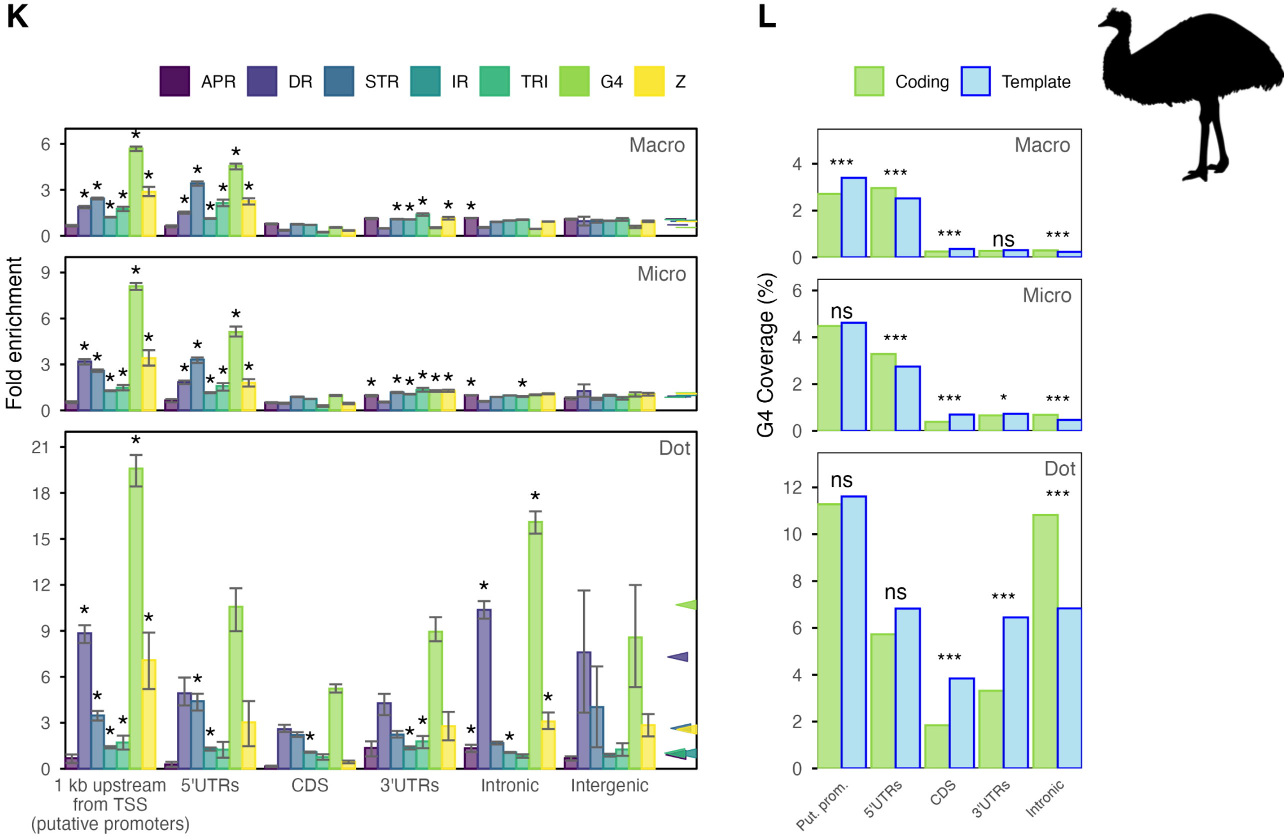

K
L
APR
DR
STR
IR
TRI
G4
Z
Coding
Template
Fold enrichment
Macro
Micro
Dot
1 kb upstream from TSS (putative promoters)
5'UTRs
CDS
3'UTRs
Intronic
Intergenic
G4 Coverage (%)
Put. prom.
ns
***

**Figure S10.** Enrichment of non-B DNA motif coverage at zebra finch tandem repeats
Fold enrichment of non-B DNA motif coverage at zebra finch tandem repeats with different repeat unit lengths.

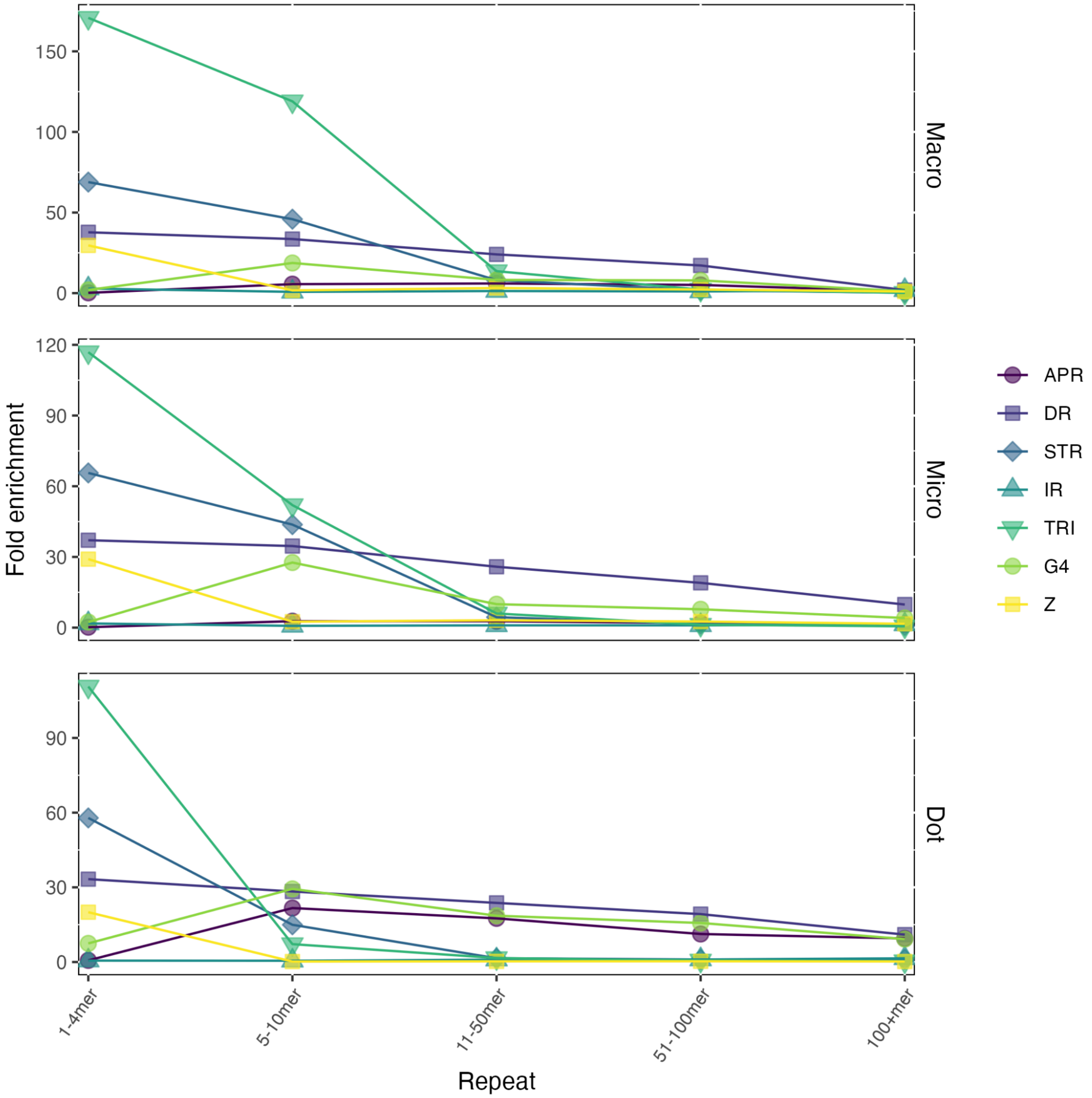

**Figure S11.** Non-B DNA motif enrichment peak region in zebra finch PAR
The PAR peak region (read rectangle) as seen on the zebra finch chromosome W in IGV. The tandem repeats are seen in blue in the third track from the bottom.

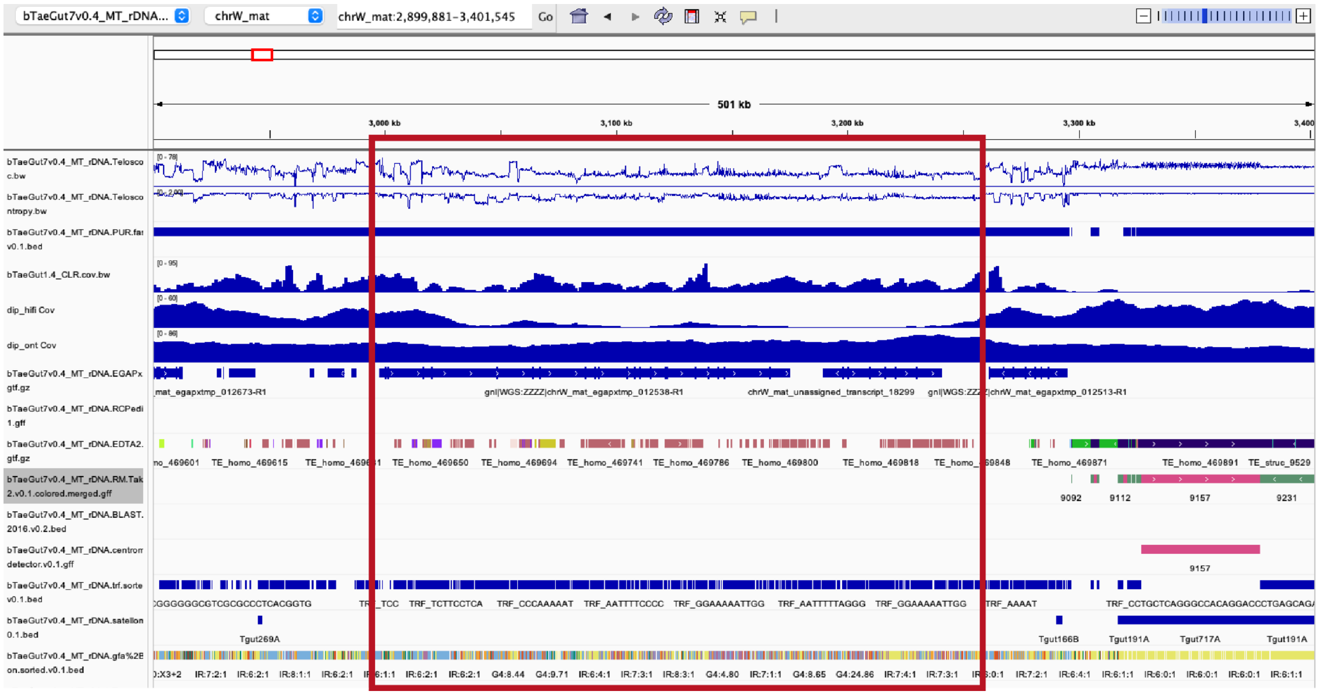

**Figure S12.** Enrichment of non-B DNA motifs at satellite repeats
Detailed enrichment pattern at all zebra finch satellites (including very short ones). Red denotes enrichment compared to the genome-wide motif content, while blue denotes depletion. Gray means the repeat class is missing from the respective chromosome category.

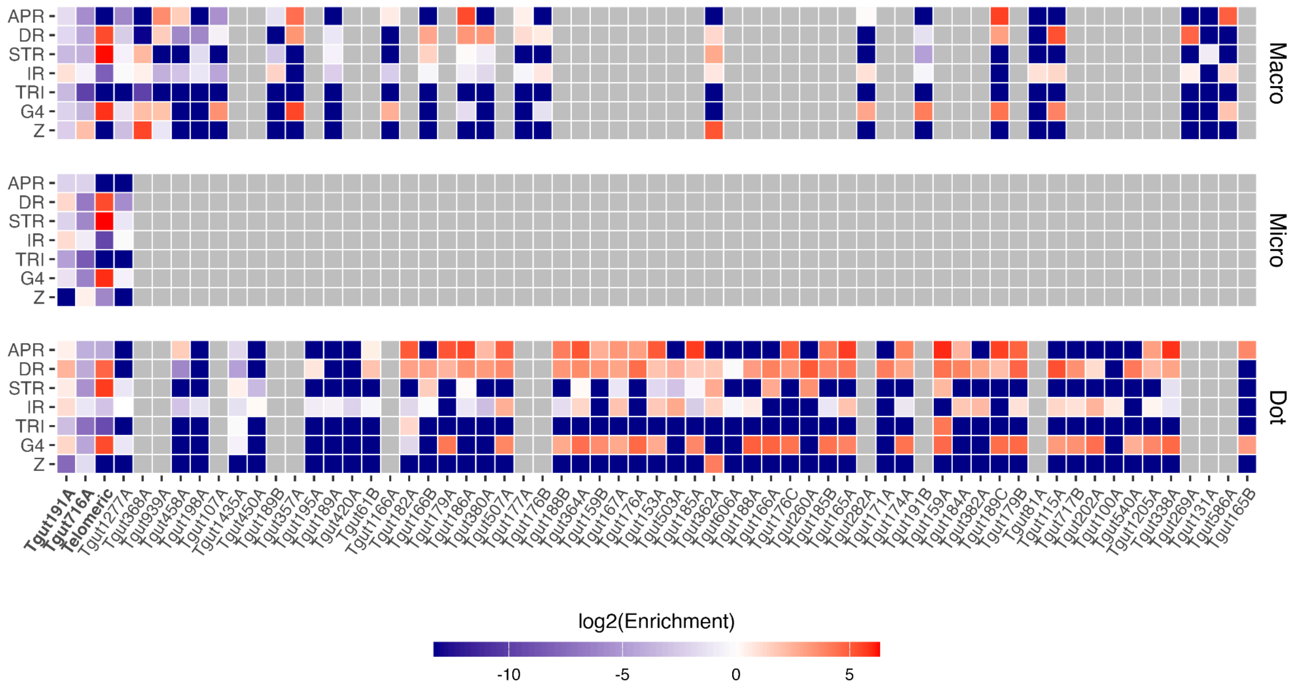

**Figure S13.** Non-B DNA motif enrichment at zebra finch centromeres.

Enrichment of non-B DNA motifs at centromeres in the zebra finch genome , compared to the genome-wide average. Red and blue denote, respectively, enrichment and depletion compared to the genome-wide non-B DNA content. Values marked with "*" are significantly enriched or depleted as compared to the genome-wide average (two-sided randomization test compared to background of 100 windows, $P < 0.05$, see Methods). Abbreviations are as in Fig. 1. See Fig. S14 for fold comparisons to the chromosome average instead of the genome-wide average.

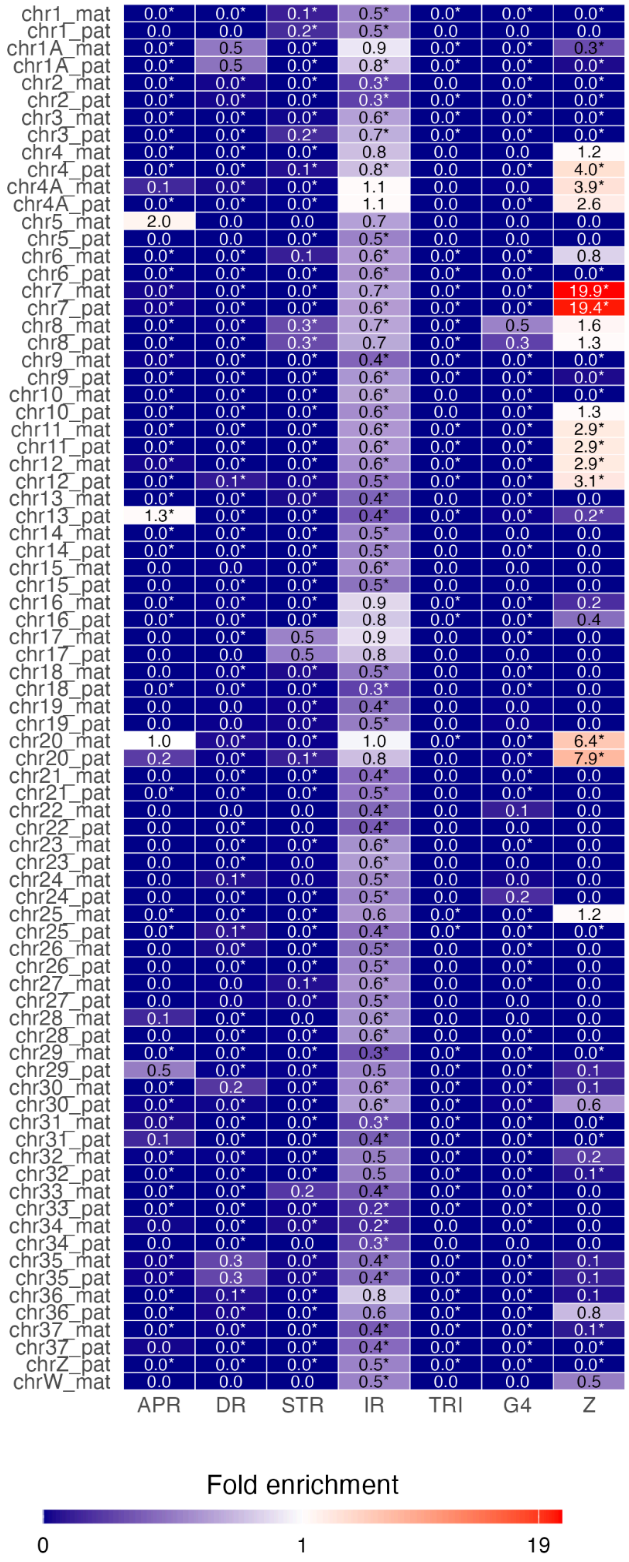

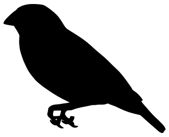

**Figure S14.** Enrichment of non-B DNA motifs at zebra finch centromeres, based on chromosome coverage

Enrichment at centromeres on all chromosomes in the diploid zebra finch genome, compared to the average coverage for each chromosome (instead of genome-wide coverage). Red colors denote enrichment, while blue denote depletion.

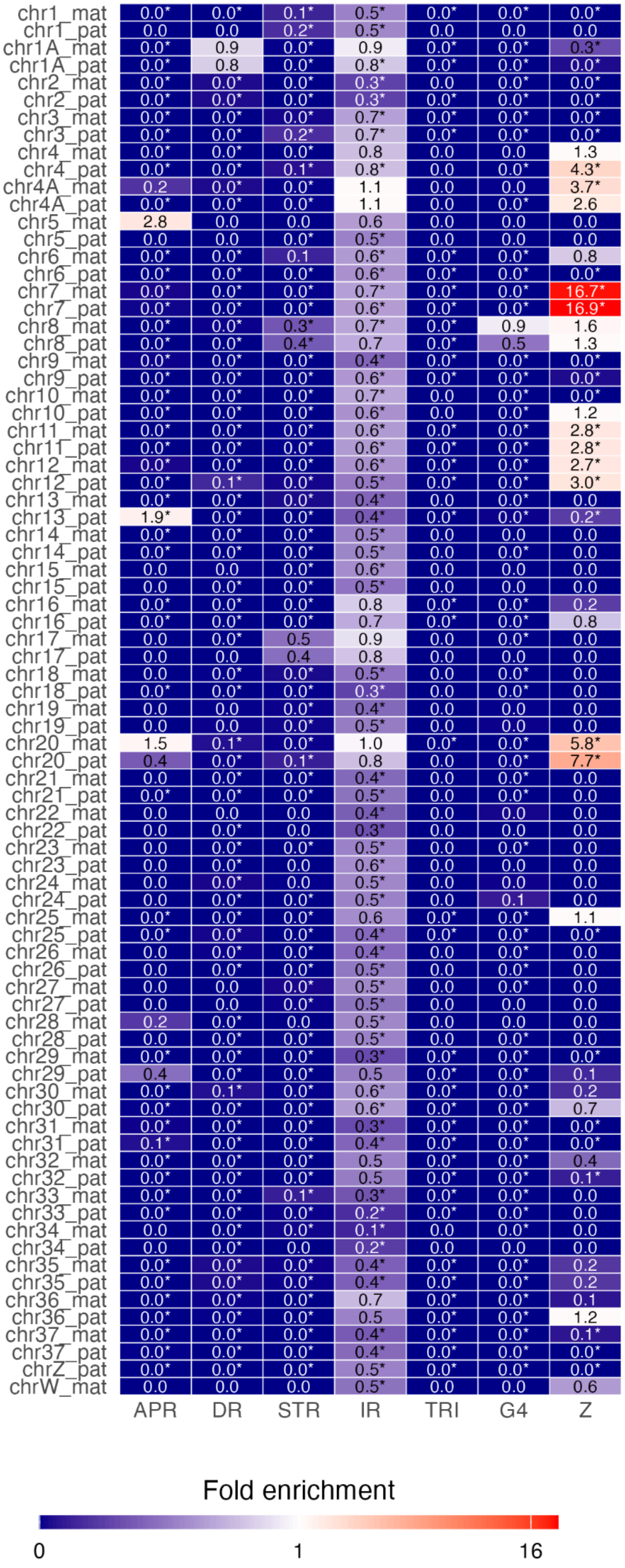

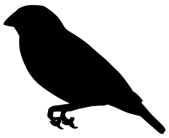

**Figure S15.** Z-DNA motif enrichment at centromeres for different algorithms

Z-DNA motif enrichment at centromeres on all chromosomes in the diploid zebra finch genome, for Z DNA motifs annotated with Z-DNA Hunter model 1 and model 2 (the latter one is used for Fig. 5), GFA and ZSeeker. Red colors denote enrichment, while blue denote depletion.

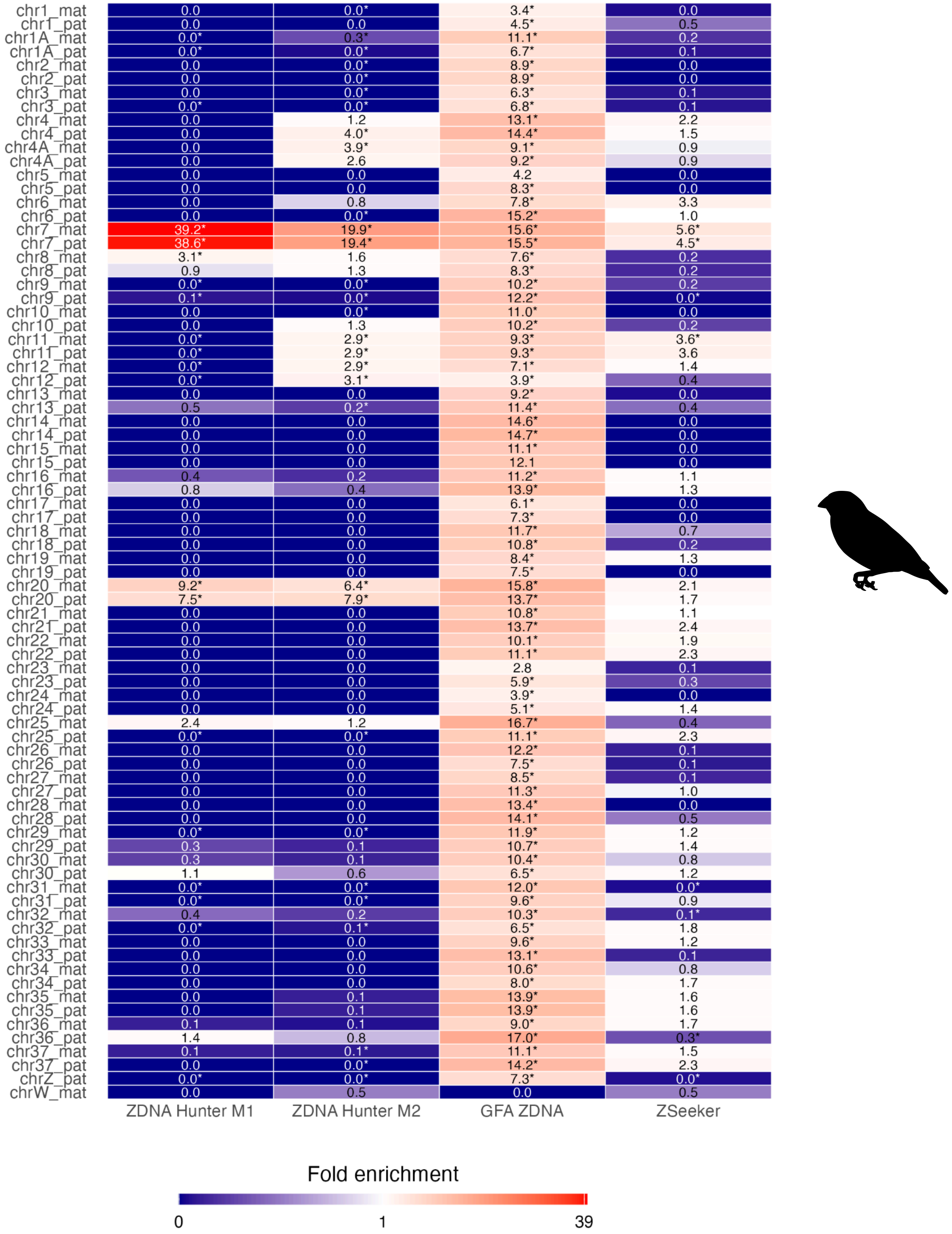

**Figure S16.** Enrichment of non-DNA motifs at chicken centromeres
Enrichment at centromeres on all chromosomes in the chicken genome. Blue denotes depletion and red enrichment; significance test as in Fig. S15. Note that centromeric positions were not available for all chicken chromosomes.

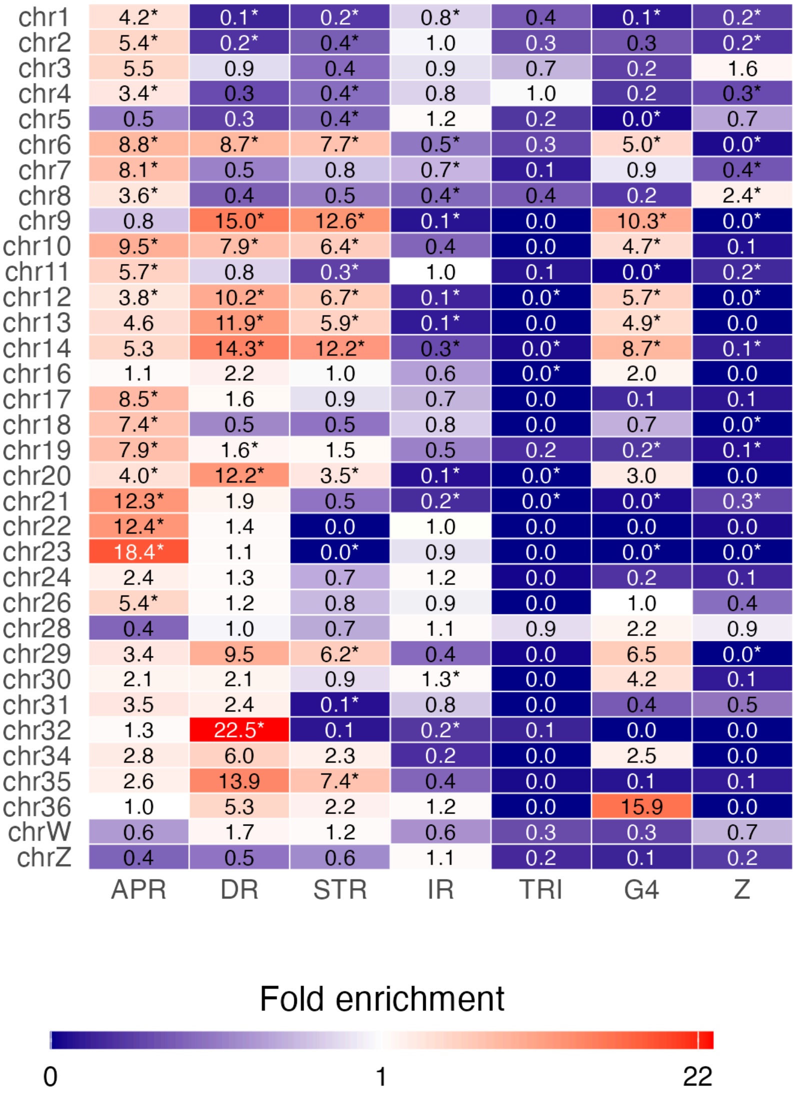

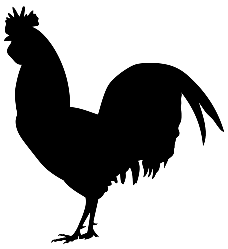

**Figure S17.** Methylation levels at G4s, strand-dependent in zebra finch

Distribution of median methylation levels per gene in zebra finch, calculated from CpG sites in G4s on the template and the coding strand, respectively.

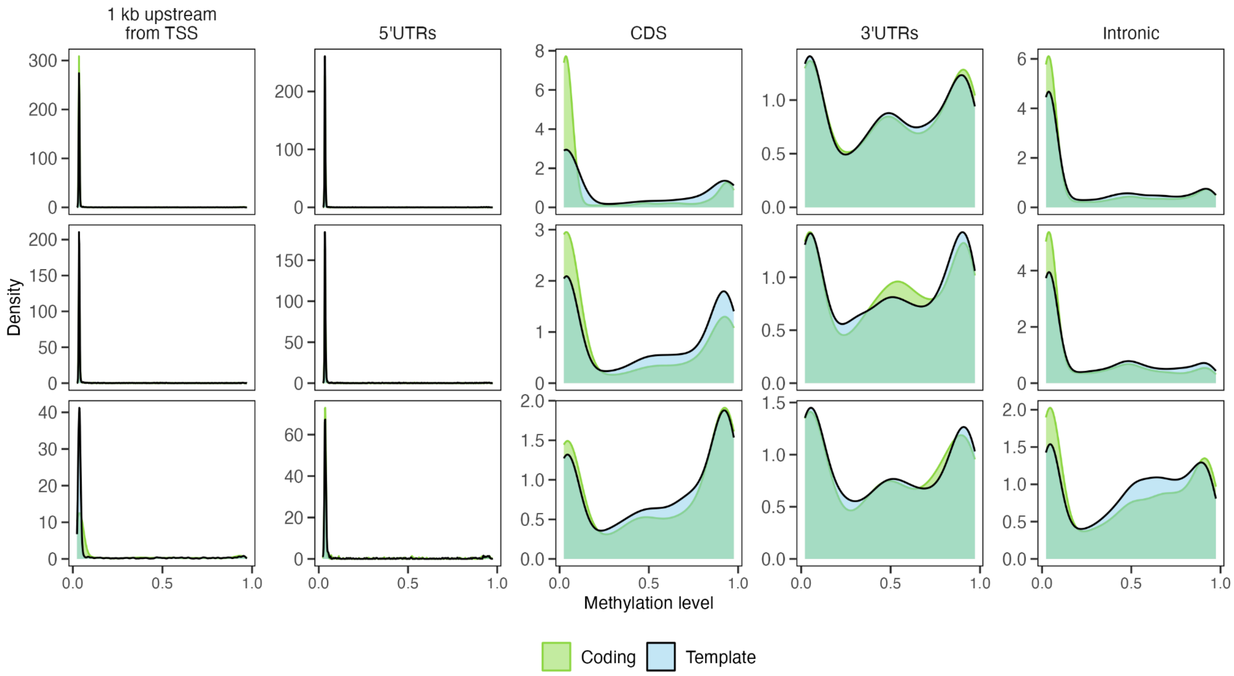

**Figure S18.** Proportion of genes with CpG sites in zebra finch

Pie charts showing the proportion of zebra finch gene regions with CpG sites in the diploid genome. The colored part of the pie represents genes with CpG sites, while the white portions represent genes that lack CpG sites. Background considers the gene region with G4s removed, while Template and Coding consider only the G4s on either the opposite strand as the gene annotation or on the same strand. The number above each circle represents the total number of gene regions in each category. The proportions of Micro and Dot categories are tested for differences with Macro using a pairwise z-test with FDR correction; significance is indicated in each circle (*** *P*<0.001, ** *P*<0.01, * *P*<0.05, n.s. - not significantly different from Macro).

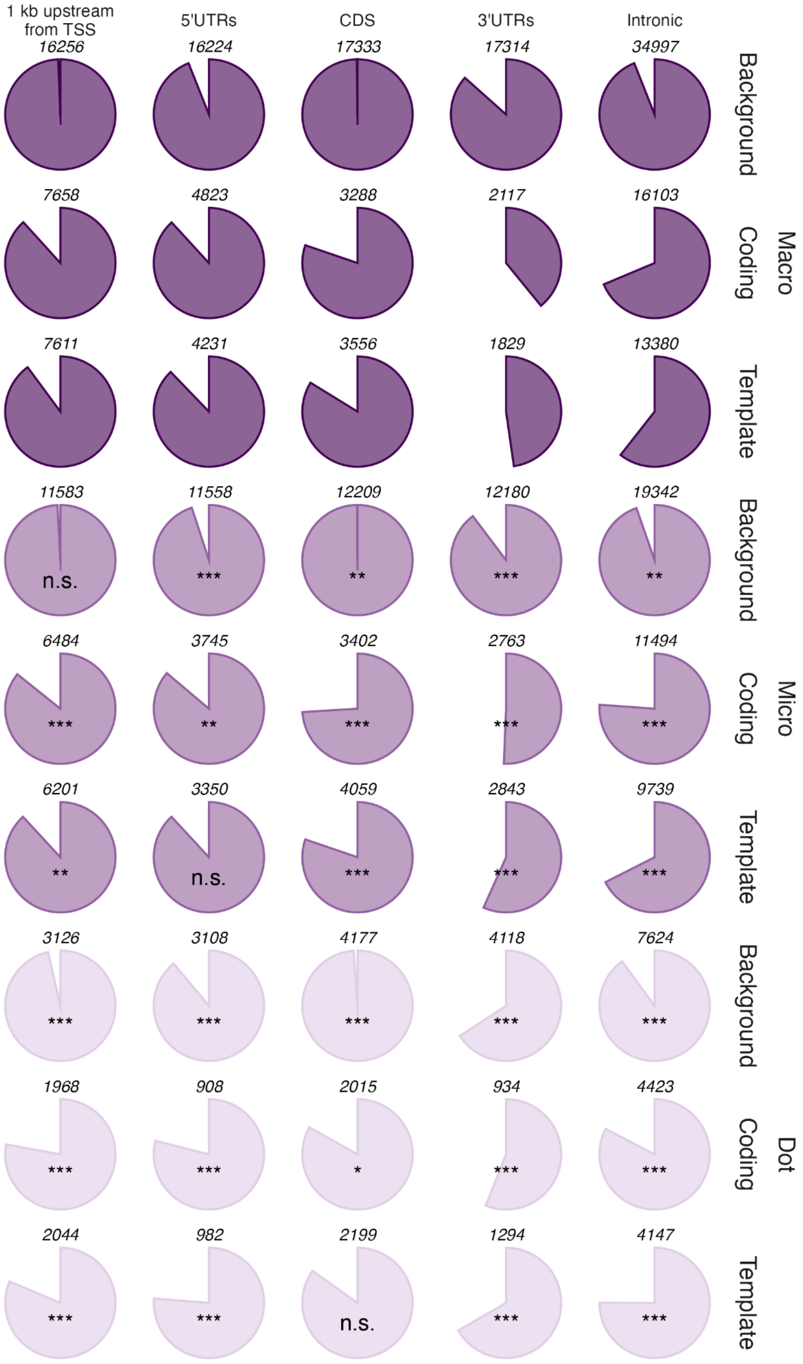

**Figure S19.** Spectroscopic characterization of G4 structures

Spectroscopic characterization of four zebra finch G4 structures: **A-D** GGTGGGGGGCAGGAGGGAAGAGG from satellite Tgu368 (5S rDNA), **E-H** GGGACTGGGGACAAGGGATGGAGGG from transposable element LINE/CR1, **I-L** GGGCTGGGCTGGGCTGGG from tandem repeat and promoters, and **M-P** GGGCAGGGATGGATGGGATATTGGG from LINE/CR1. Panels **A,E,I,M** show Circular dichroism (CD) spectra (same as Fig. 4D). Panels **B,F,J,N** show temperature-dependent difference absorbance spectra (delta absorbance) across the wavelength range 240–320 nm. The left spectrum shows data in 100 mM KCl with 140 mM LiCl buffer; the right spectrum shows data in 140 mM LiCl buffer alone. Panels **C,G,K,O** show thermal melting curves monitoring molar absorptivity at 260 nm as a function of temperature (5–95°C), and panels **D,H,L,P** show the same at 294 nm.

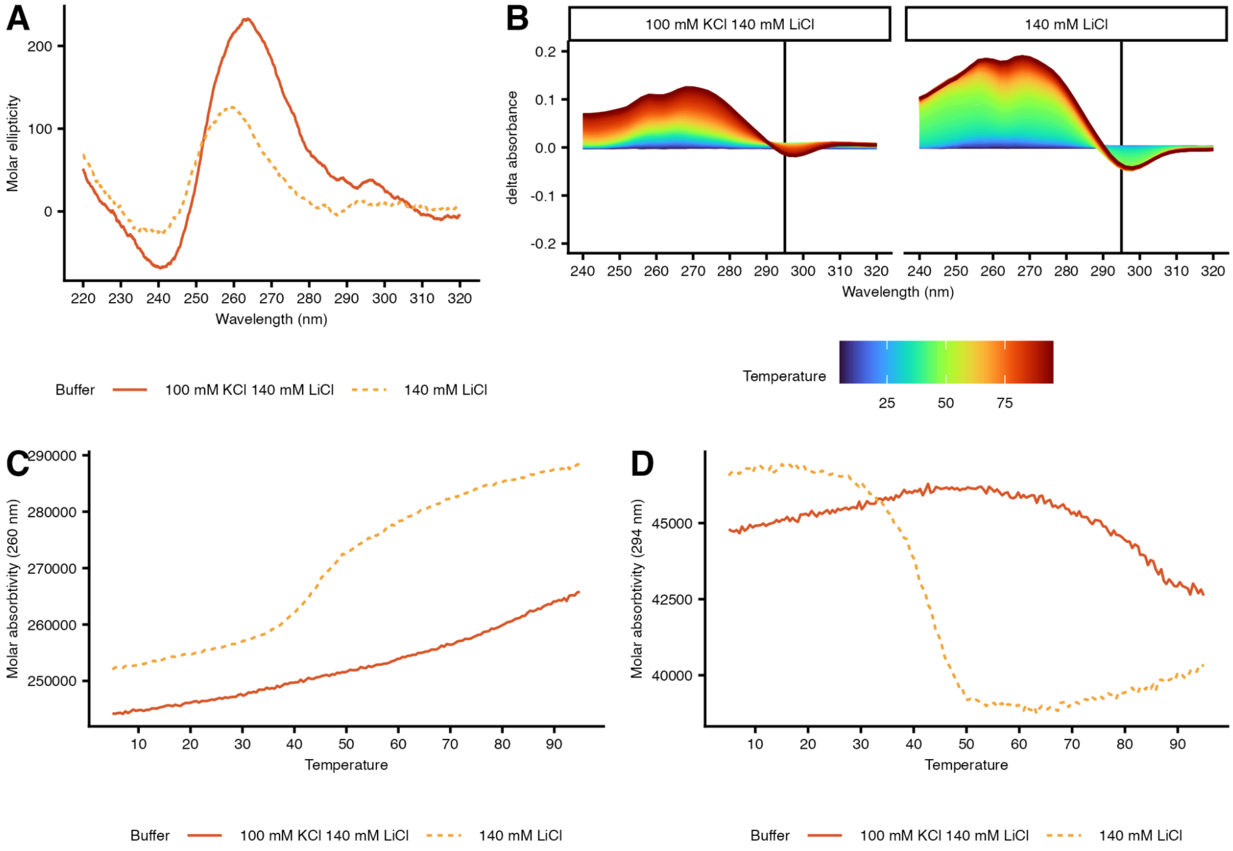

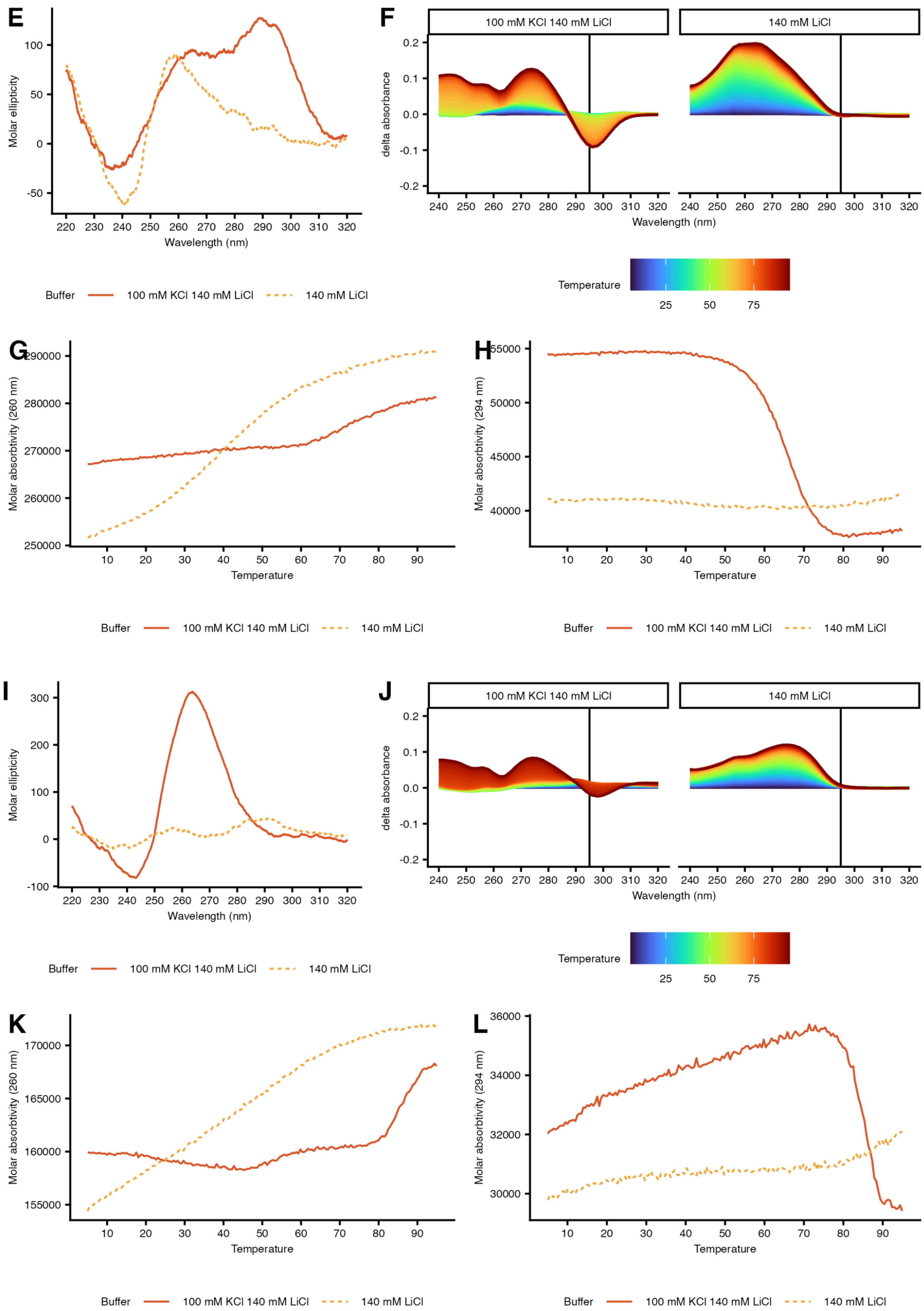
E
Molar ellipticity
Wavelength (nm)
Buffer
100 mM KCl 140 mM LiCl
140 mM LiCl
F
100 mM KCl 140 mM LiCl
140 mM LiCl
delta absorbance
Wavelength (nm)
Temperature
25
50
75
G
Molar absorbtivity (260 nm)
Temperature
Buffer
100 mM KCl 140 mM LiCl
140 mM LiCl
H
Molar absorbtivity (294 nm)
Temperature
Buffer
100 mM KCl 140 mM LiCl
140 mM LiCl
I
Molar ellipticity
Wavelength (nm)
Buffer
100 mM KCl 140 mM LiCl
140 mM LiCl
J
100 mM KCl 140 mM LiCl
140 mM LiCl
delta absorbance
Wavelength (nm)
Temperature
25
50
75
K
Molar absorbtivity (260 nm)
Temperature
Buffer
100 mM KCl 140 mM LiCl
140 mM LiCl
L
Molar absorbtivity (294 nm)
Temperature
Buffer
100 mM KCl 140 mM LiCl
140 mM LiCl

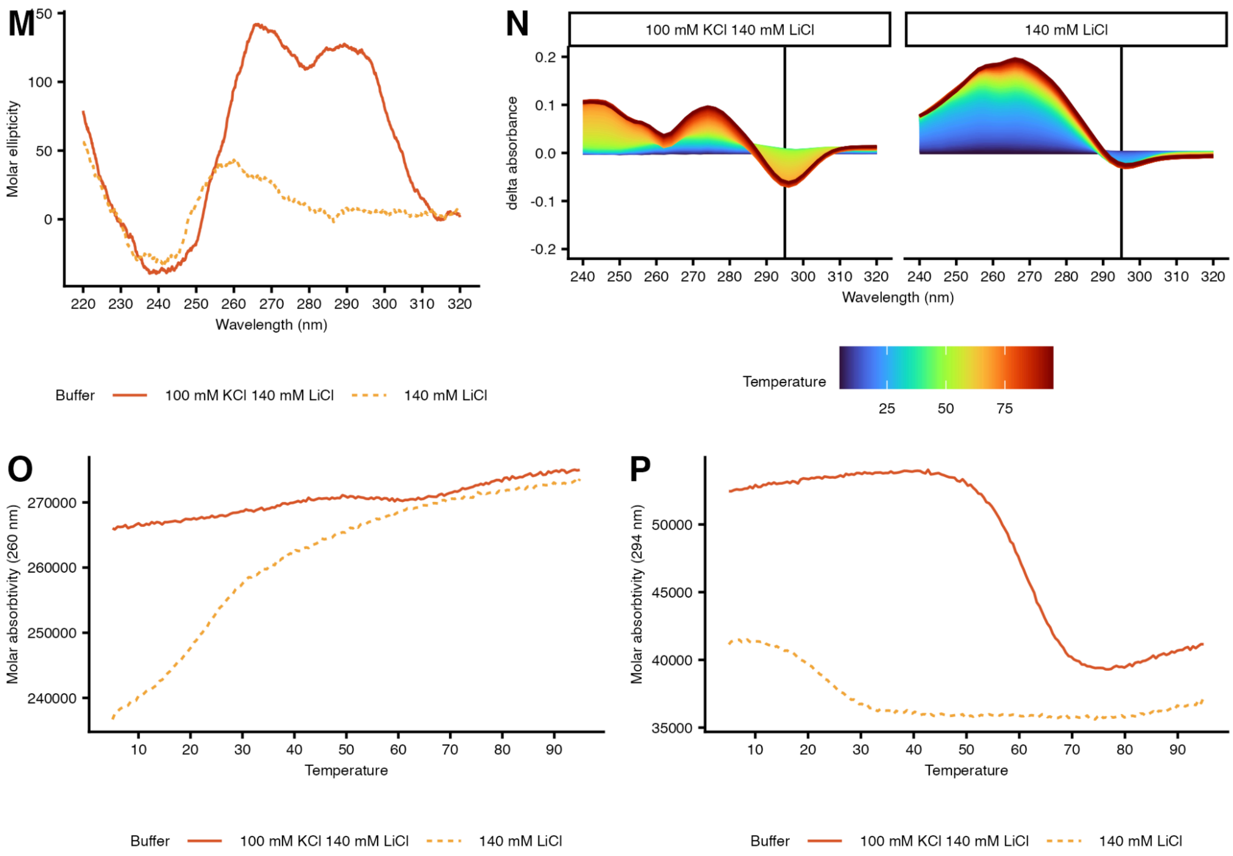
M
N
O
P
Molar ellipticity
Wavelength (nm)
Buffer
100 mM KCl 140 mM LiCl
140 mM LiCl
delta absorbance
Temperature
Molar absorbtivity (260 nm)
Molar absorbtivity (294 nm)

**Figure S20.** Native gel analysis of DNA structure

Samples were fractionated on a 15% native gel supplemented with 10 mM KCl and stained with SYBR gold. The native gel was run at 4°C. Tgut368A (5S rDNA): GGTGGGGGGCAGGAGGGAAGAGG, LINE/CR1 (second from left): GGGACTGGGGACAAGGGATGGAGGG, TR/promoter: GGGCTGGGCTGGGCTGGG, LINE/CR1 (second from right): GGGCAGGGATGGATGGGATATTGGG.

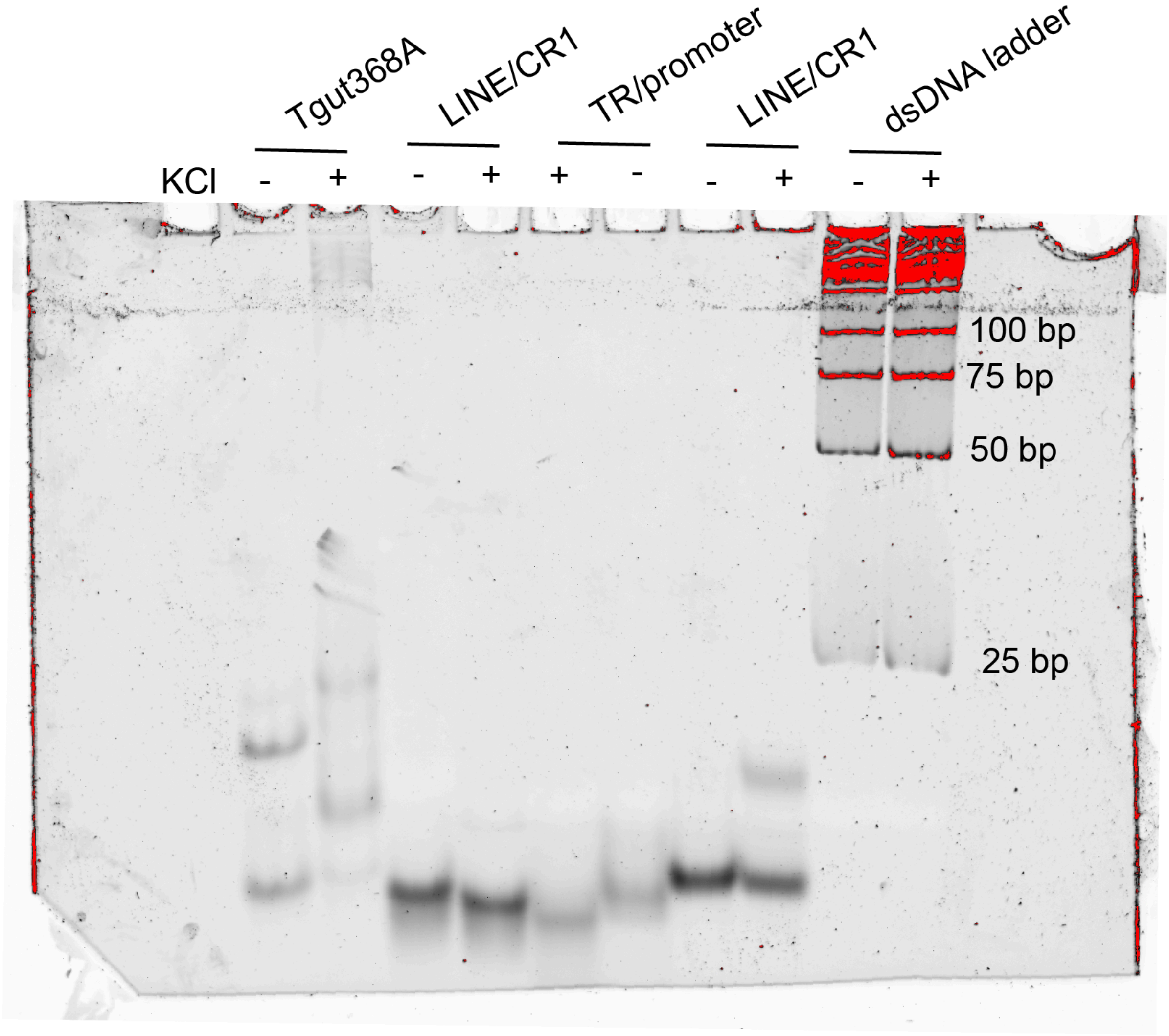

**Figure S21.** ChIP-seq enrichment around chicken transcription start sites (TSS)
Genome-wide ChIP-seq enrichment was quantified as the RPKM-normalized ratio of ChIP to input control signal across a 6-kb window centered on the TSS (±3 kb, 10-bp bins) of all annotated chicken genes. Heatmaps are grouped after chromosome categories: *(A)* genes on macrochromosomes, *(B)* genes on microchromosomes and *(C)* genes on dot chromosomes, sorted by mean signal across the plotted window (highest to lowest). Color scale is shared across all three heatmaps and capped at the 99th percentile of signal genome-wide to allow direct visual comparison of enrichment intensity between chromosome classes. A ratio of 1 indicates no enrichment relative to control.

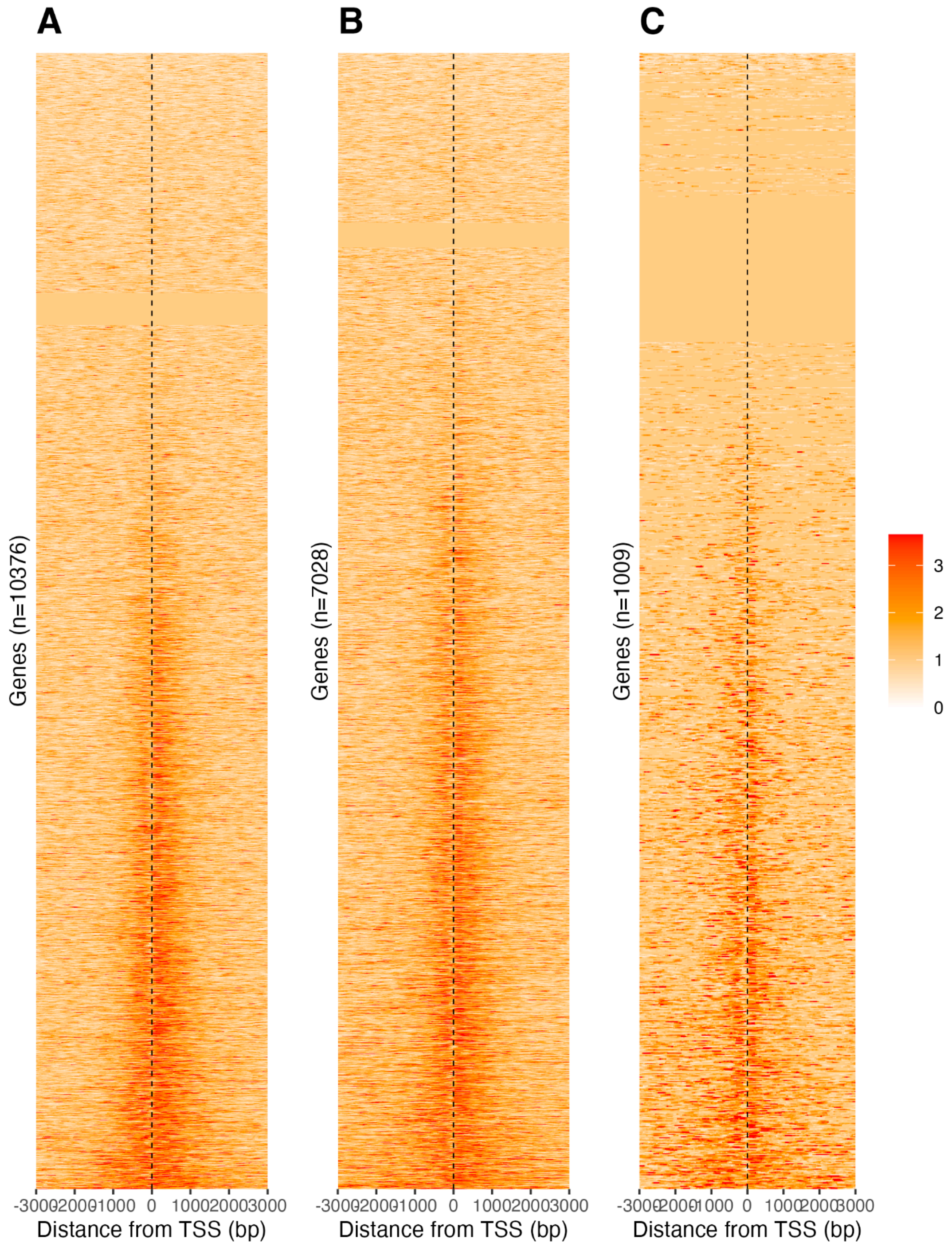

**Figure S22.** G4 ChIP-seq peak enrichment in chicken functional regions
Fold enrichment of G4 ChIP-seq peak density in functional regions compared to *(A)* genome-wide peak density and *(B)* category specific peak density (eg. different baselines for macro, micro and dot chromosomes respectively).

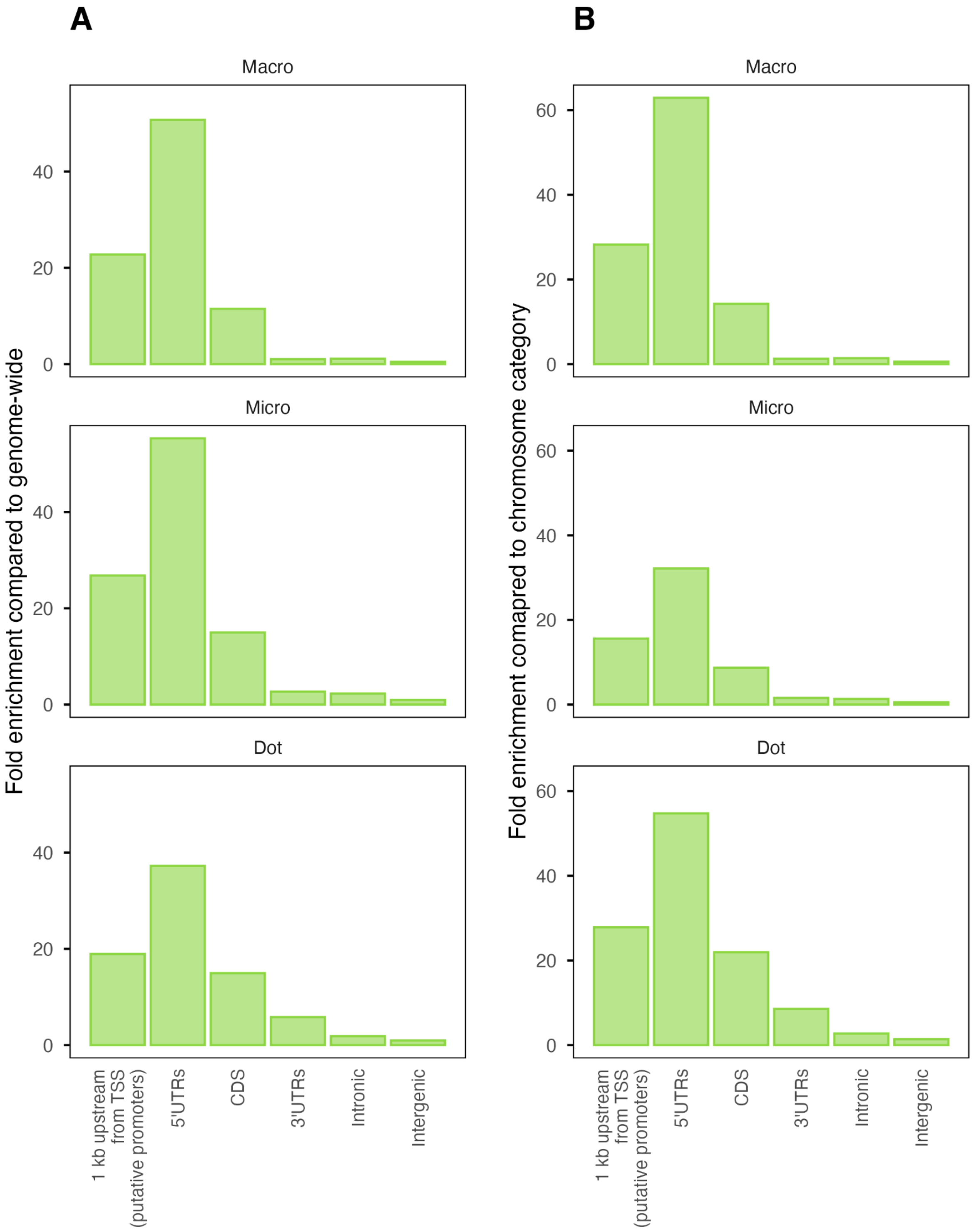

**Figure S23.** PDAL-Seq signal is depleted at APR, DR, STR, and IR non-B DNA motifs in non-repetitive regions of the genome and HMM model selection.

*(A)* CFS414 cell line PDAL-Seq mean signals around non-B DNA motifs in uniquely mapping, non-repetitive regions. PDAL-Seq data are compared to PCR-free whole-genome DNA sequencing data to control for sequencing, mapping, and copy number artifacts. Mean read coverage (CPM per 100 bp bin, both strands summed) is plotted against distance from the feature centre (bp). *(B)* Model selection for the Gaussian HMM fitted to z-scored combined PDAL-Seq coverage in 1 kbp windows (blacklisted, sex-chromosome, chrMT and rDNA windows removed). Bayesian information criterion versus number 2–30 hidden states in each model. The vertical red line represents the HMM (k = 5) presented in the main manuscript.

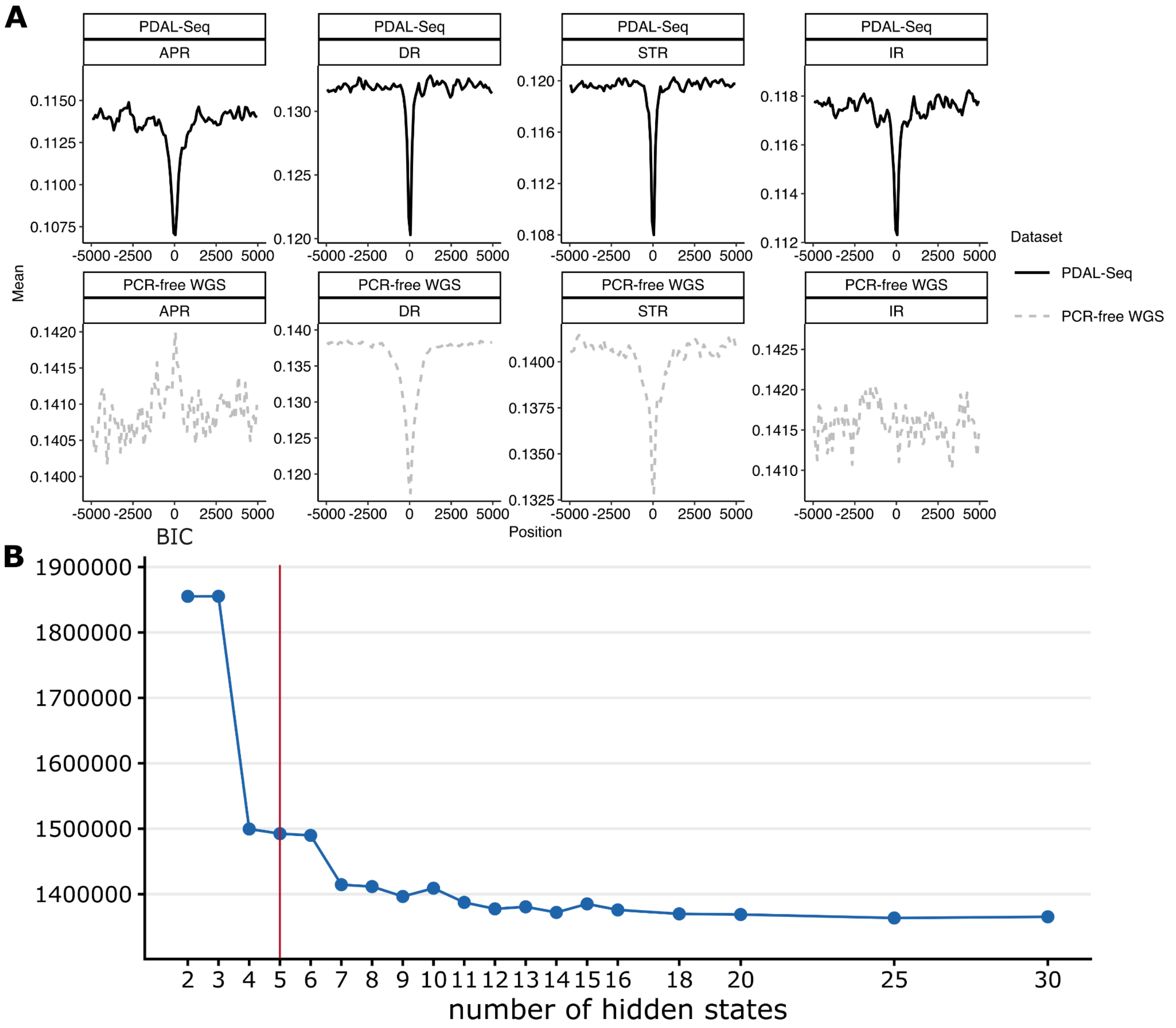

**Figure S24.** Non-B DNA motif coverage in zebra finch A and B compartments

Violin plot of Non-B DNA motif coverage distribution at the A compartments (euchromatic) and the B compartments (heterochromatic) for *(A)* macrochromosomes, *(B)* microchromosomes and **C** dot chromosomes (averaged across the compartments for each chromosome and haplotype separately). Individual chromosomes are seen as dots, and gray lines between violins connect the compartments from the same chromosome. *$P$ <0.05; *$P$ <0.01; ***$P$ <0.001; ns not significant. Wilcoxon test adjusted for multiple testing using FDR.

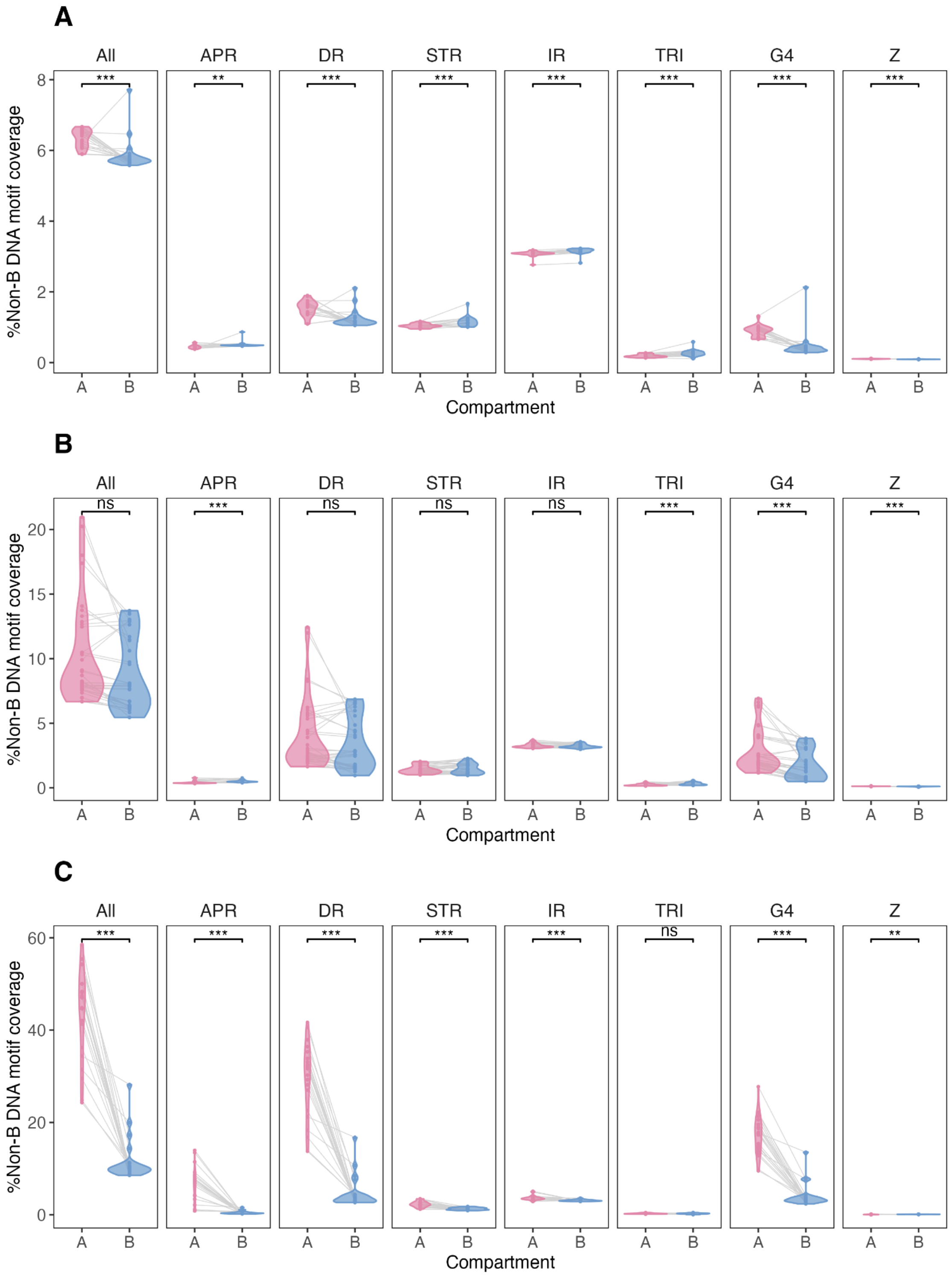

**Figure S25.** Enrichment of non-B DNA at vs. outside of TRs in introns in zebra finch

*(A)* Fold enrichment (as compared to genome-wide) of non-B DNA motifs at introns in zebra finch dot chromosomes (A and B compartments separated), comparing tandem repetitive sequence with the rest of the introns ("outside tandem repeat"). Inset pie charts show the fraction of intronic base pairs that is also annotated as tandem repetitive, in dot chromosome A compartments (pink) and B compartments (blue). *(B)* Number of annotated tandem repeats on dot chromosome introns with different unit lengths, separated by compartment.

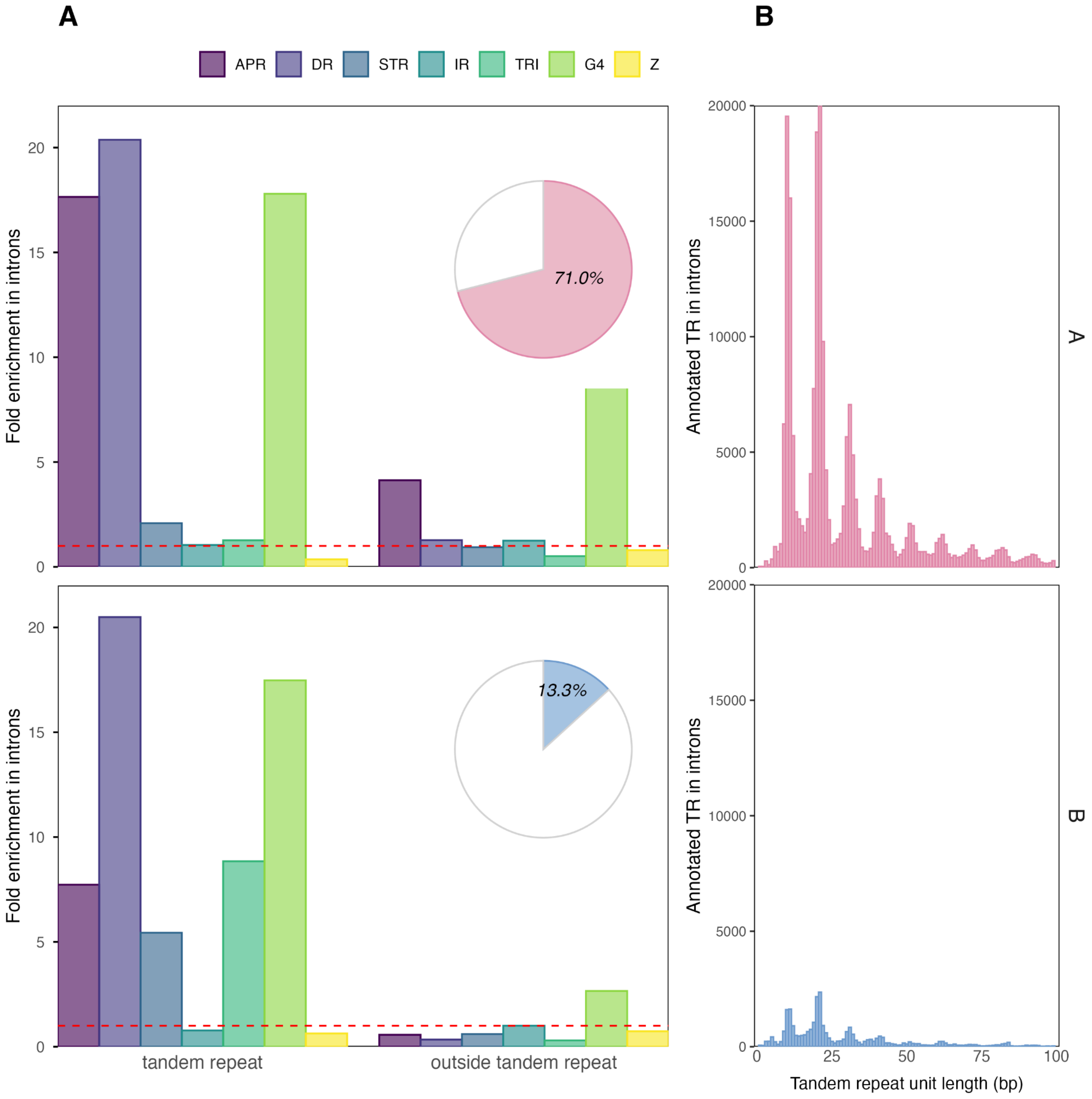

**Figure S26.** Non-B DNA motif coverage and HiFi sequence coverage for zebra finch macro and microchromosomes

Correlation between non-B DNA motif coverage and PacBio HiFi sequencing coverage in zebra finch, on macro- and microchromosomes, in 1,024-bp non-overlapping windows. Dark red dashed lines show linear regression fits, and Pearson's $R^2$ (red) and Spearman's $\rho^2$ (blue) are shown for each correlation. Windows with sequence coverage greater than 100× are not displayed for visualization purposes. Abbreviations are as in Figure S1.

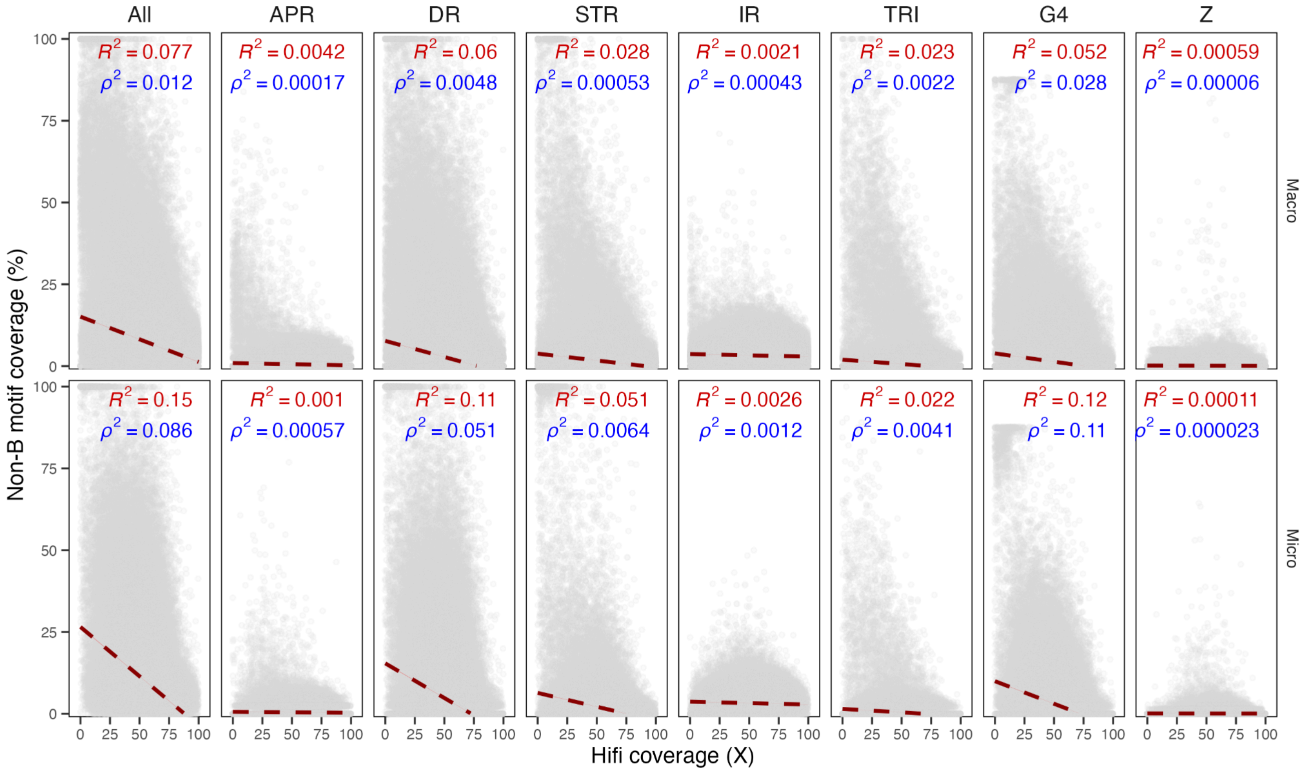

**Figure S27.** Non-B DNA motif coverage and ONT sequence coverage for zebra finch macro and microchromosomes

Same as Figure S22 but for ONT data.

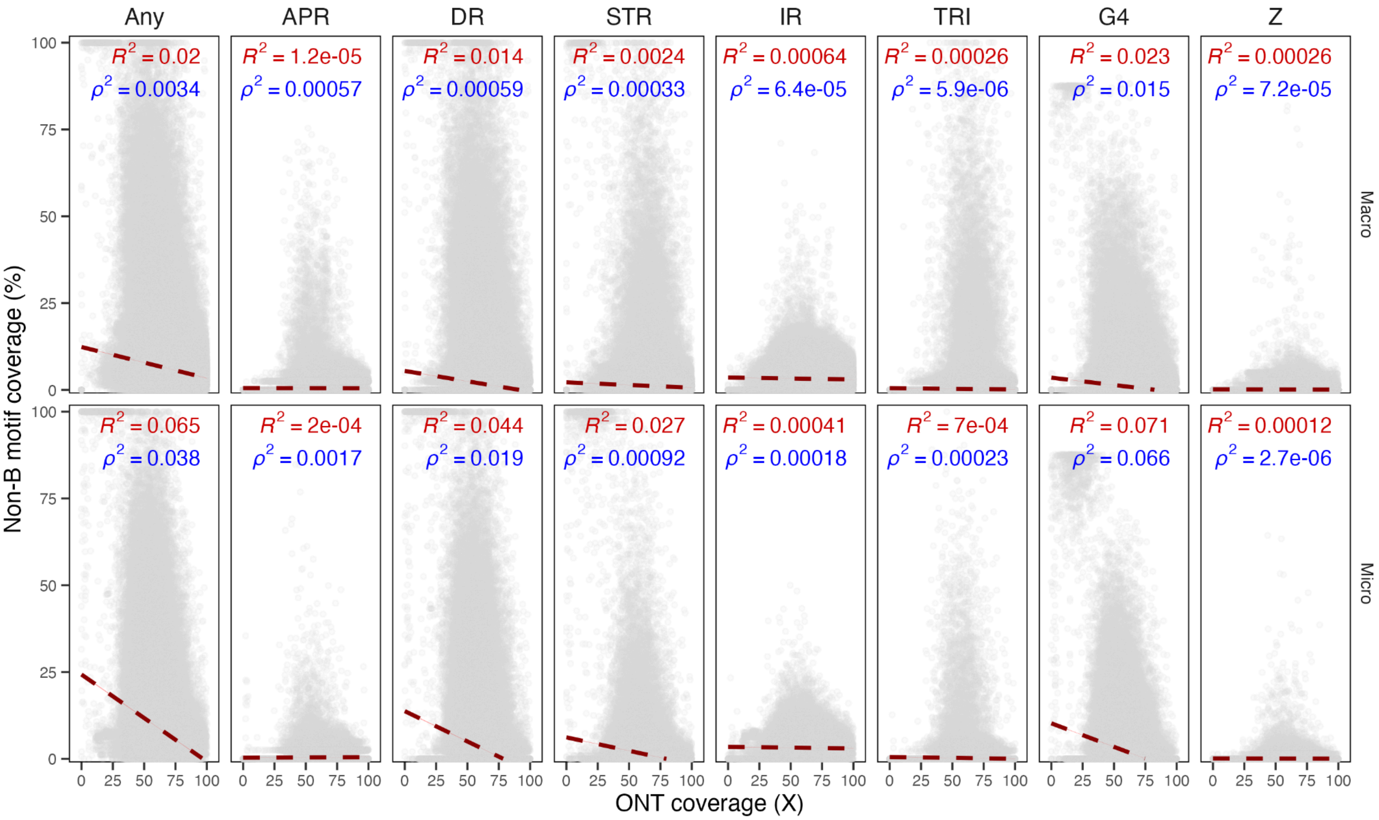

**Figure S28.** Spacer and repeat length distributions

Spacer and repeat length distributions for direct repeats, inverted repeats, and mirror repeats found by gfa. Red dashed lines show the thresholds used in this study.

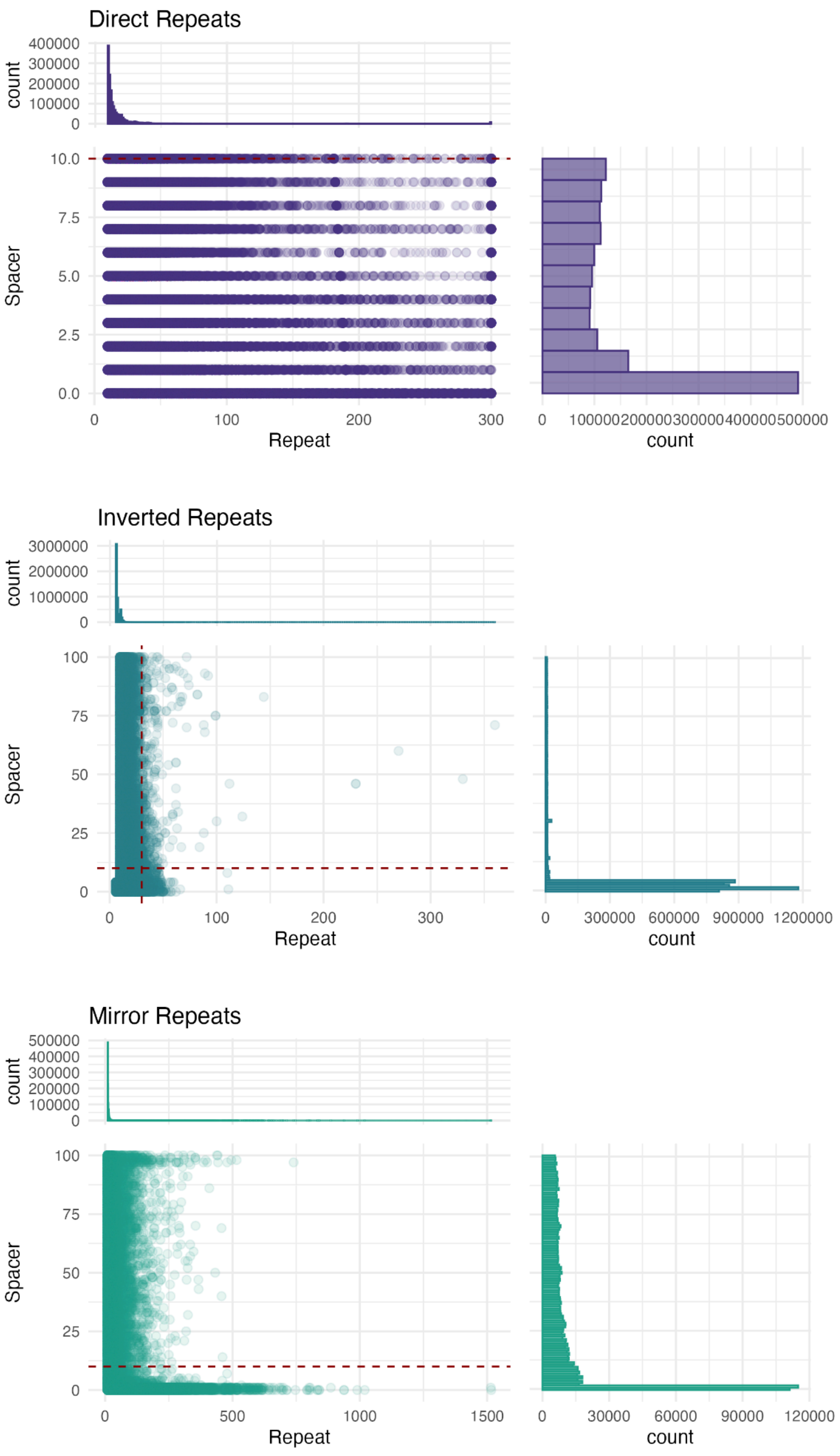

**Figure S29.** Comparisons between annotation software

Comparisons between g4discovery and Quadron for G4 motifs and gfa, Z-DNA Hunter (two models) and ZSeeker for Z-DNA, as visualized for three chromosomes as examples. G4discovery was chosen for G4, and Z-DNA Hunter model2 were chosen for the study.

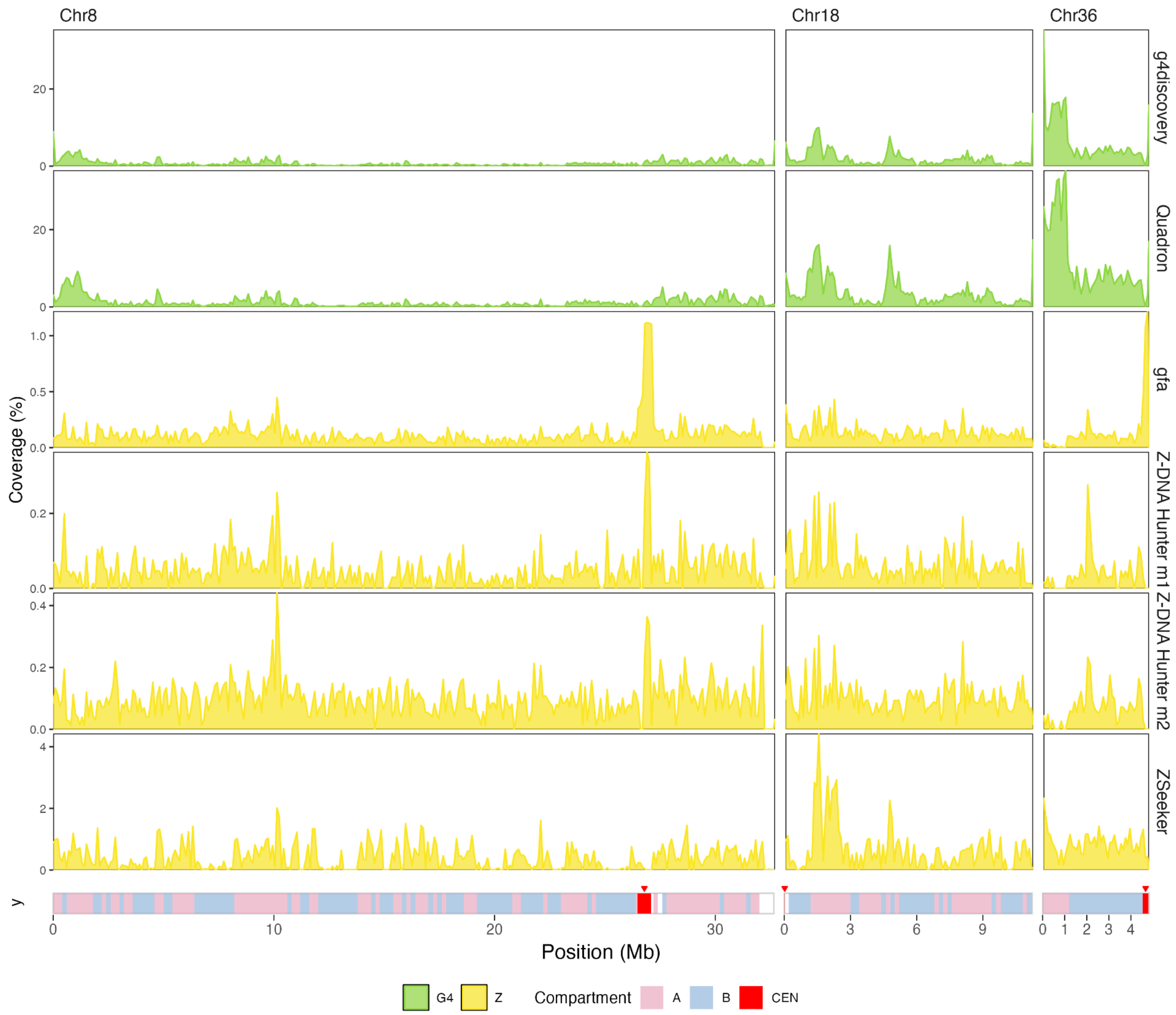
